# Exploring Adenosine Analogues for Chondrosarcoma Therapy: In Vitro and In Vivo Insights

**DOI:** 10.1101/2024.12.23.630093

**Authors:** Marion Lenté, Juliette Aury-Landas, Mahdia Taieb, Benoît Bernay, Eva Lhuissier, Karim Boumédiene, Catherine Baugé

## Abstract

Chondrosarcoma (CS) is described as resistant to conventional chemotherapy and radiotherapy. The development of new therapeutic approaches is necessary. The aim of the present study is to validate the use of adenosine analogues as a new therapeutic strategy in the treatment of CS.

Five adenosine analogues (aristeromycin, cladribine, clofarabine, formycin, and pentostatin) were evaluated *in vitro* on several chondrosarcoma cell lines using both 2D cultures and 3D alginate bead models. Cell viability was assessed using Acridine Orange and DAPI staining, or ATP assay. Apoptosis was measured via Annexin V and Propidium Iodide staining, while cell cycle progression was analyzed with DAPI staining. The most promising compounds were further tested *in vivo* using a xenograft chondrosarcoma model in nude mice.

Results showed that four analogues (aristeromycin, formycin, cladribine, and clofarabine) significantly reduced cell viability in 2D cell cultures. Of these, cladribine and clofarabine demonstrated potent efficacy in both 2D and 3D models by inducing apoptosis. Cladribine was further found to induce cell cycle arrest, leading to apoptosis-mediated cell death. *In vivo*, both cladribine and clofarabine exhibited substantial antitumor effects in a xenograft model.

In conclusion, cladribine and clofarabine, which are already approved for clinical use in leukemia and multiple sclerosis, show promise as potential candidates for chondrosarcoma treatment. Their efficacy in preclinical models suggests these molecules could be repurposed for Phase II clinical trials in CS patients.

## Introduction

Chondrosarcomas (CS) are primary malignant bone tumors characterized by the production of a cartilaginous matrix. They rank as the third most common primary malignant tumors after myelomas and osteosarcomas, accounting for 20-27% of all malignant bone tumors in adults. Annually, they represent 2.1 to 4 new cases per million people [1,2].

According to the European Society for Medical Oncology (ESMO) guidelines, the gold standard for treating conventional CS, which constitutes approximately 85% of all CS, is surgical intervention. This may involve either bone curettage (a minimally invasive technique) or extensive ablation, which can include amputation of bone and surrounding soft tissues, leading to increased morbidity. Furthermore, the feasibility of surgery depends on the tumor’s location and stage; in some cases, surgical options may be limited. Conventional CS are notably resistant to radiotherapy and chemotherapy; while radiotherapy can serve as a palliative measure, the overall evidence for its effectiveness in patient studies is sparse due to the rarity of the disease. Consequently, ongoing research aims to discover new therapeutic approaches and enhance the efficacy of existing chemotherapies to provide better patient care. In this context, scientists are focusing on developing innovative anti-cancer therapies, particularly through the design of new or repositioned drugs, with purine analogues representing a significant category.

Purines are essential components of deoxyribonucleic acid (DNA), ribonucleic acid (RNA), and other biomolecules, including adenosine triphosphate (ATP) and S-adenosyl-methionine (SAM). Due to their critical roles in various cellular processes, such as energy production, signaling, DNA synthesis, and repair, purines and their metabolites have been implicated in cancer development and in the mechanisms of radiotherapy resistance [3,4]. Therefore, targeting purine metabolism presents a promising therapeutic avenue for treating cancers, particularly bone cancers as well as other malignancies [5,6].

Since the 1950s, purine antimetabolites have been developed to inhibit purine synthesis, including a variety of purine analogues. These compounds share structural similarities with purines and can incorporate themselves into purine nucleotides and DNA, thus competing for essential cellular processes. They comprise a diverse group of molecules with varying mechanisms of action, indications, and pharmacokinetics. Notably, adenosine analogues have been developed over time to enhance their antineoplastic efficacy [7]. The discovery of 3-Deazaneplanocin A (DZNep) in the 1980s marked a significant advancement in this field. Numerous studies have since demonstrated its anticancer effects across a wide range of cancers including chondrosarcomas, as evidenced by our own *in vitro* and *in vivo* research [8–10]. DZNep has also been shown to enhance chemosenstivity specifically in CS *in vitro* [11].

Given these findings, we hypothesized that other adenosine analogues may also exert therapeutic effects on chondrosarcomas. To investigate this potential, we selected five additional adenosine analogues: two natural analogues, aristeromycin and formycin, along with three clinically used analogues for treating various leukemias: pentostatin, cladribine, and the cladribine derivative clofarabine [12–15]. Aristeromycin, a unique carbocyclic nucleoside antibiotic derived from *Streptomyces citricolor*, has been shown to reduce cell viability in prostate cancer cells and, like DZNep, inhibits S-adenosylhomocysteine hydrolase (SAHH), an enzyme critical to the transmethylation cycle and epigenetic processes [16]. Formycin A, isolated from Streptomyces kaniharaensis, is noted for its potent antibiotic properties and has also been recognized as a ribonucleoside analogue with significant anti-HIV-1 activity without cytotoxic effects [17,18]. Pentostatin, cladribine, and clofarabine are established adenosine analogues used clinically for treating various leukemias [19,20]. Pentostatin acts as an adenosine deaminase inhibitor with demonstrated anti-tumor activity [21,22]. Cladribine and clofarabine, in their active forms, inhibit ribonucleotide reductase (RR), an enzyme crucial for synthesizing deoxyribonucleotide triphosphates (dNTPs). This inhibition results in reduced dNTP pools, impairing DNA synthesis and repair, ultimately leading to increased DNA damage and triggering programmed cell death. Clofarabine has also shown antitumoral effects in Ewing’s sarcoma and other sarcomas [23].

Given these insights, we have undertaken investigations into the effects of these adenosine analogues on cell viability, cell death, and overall anti-tumor efficacy in chondrosarcoma.

## Materials and Methods

### Cell culture

The human chondrosarcoma cell lines JJ012 and FS090 were generously provided by Dr. J.A. Block from Rush University Medical Center [24]. The CH2879 cell line was kindly donated by Professor A. Llombart-Bosch from the University of Valencia, Spain [25]. The chondrosarcoma cell line SW1353 was obtained from the American Type Culture Collection (ATCC, Manassas, VA, USA). Human chondrocytes were isolated, as previously described [26], from the femoral heads of patients undergoing arthroplasty, following the acquisition of informed consent in accordance with local legislation and ethical guidelines established by the Comité de Protection des Personnes Nord Ouest III.

All cells were cultured in Dulbecco’s Modified Eagle Medium (DMEM; LONZA, Levallois Perret, France), supplemented with 10% fetal bovine serum (FBS; Dutscher, Bernolsheim, France) and antibiotics (penicillin and streptomycin; LONZA). The cultures were maintained at 37°C in a humidified atmosphere with 5% CO_2_. Regular testing for mycoplasma contamination was conducted using PCR techniques.

For three-dimensional (3D) cultures, the SW1353 and JJ012 cell lines were encapsulated in alginate beads, as previously described [10]. Briefly, 2 million cells per mL were suspended in 1 mL of a 1.2% sodium alginate solution (Merck, Saint-Quentin-Fallavier, France). Beads were formed by dripping the cell suspension into a sterile 100 mM CaCl₂ solution (VWR, Fontenay-sous-Bois, France). The beads were subsequently washed in 0.15 M NaCl solution (Sigma Aldrich) and incubated in DMEM for two weeks prior to further treatments.

### Adenosine analogue drugs

Five molecules have been selected for to their analogy with adenosine (figure 1). Aristeromycin was provided by Santa Cruz Biotechnology (Clinisciences, Nanterre, France). Pentostatin and formycin were provided by Sigma Aldrich (Merck, Saint-Quentin-Fallavier, France). Cladribine and clofarabine were provided by MedChem (Clinisciences). Aristeromycin, cladribine and clofarabine were resuspended in dimethyl sulfoxide (DMSO, Dutscher). Pentostatin and formycin were dissolved in DMEM and Dulbecco’s Phosphate Buffered Saline with no Calcium or Magnesium (DPBS; LONZA) respectively.

**Figure 1:**
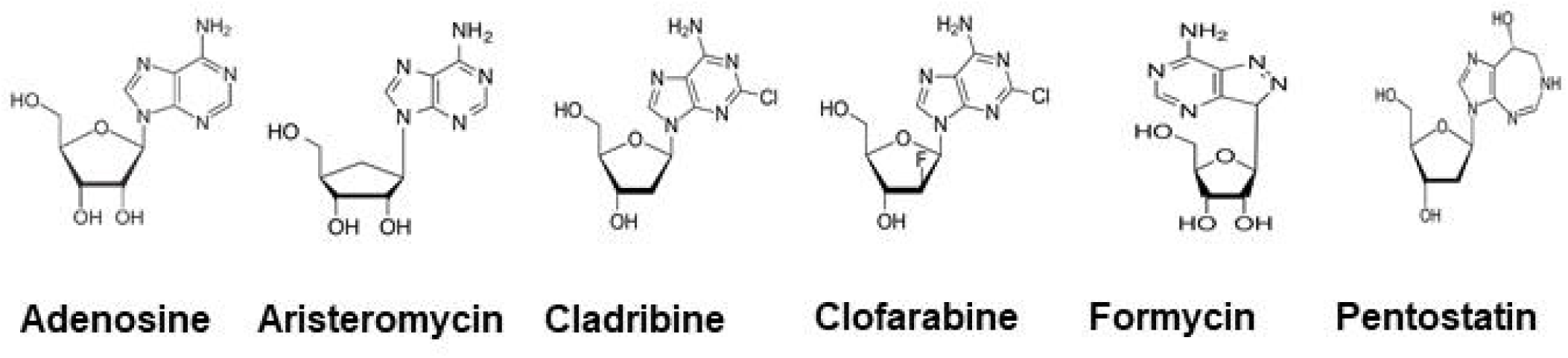
Chemical structures of adenosine and its analogues used in the study. Chemical structure of adenosine and of the five analogues used: aristeromycin, cladribine, clofarabine, formycin and pentostatin.

### Cell Viability Assessment

Two methods were employed to measure cell viability: cell staining assay using Acridine Orange (AO) and 4’,6-diamidino-2-phenylindole (DAPI) (Chemometec, Denmark), and the CellTiter-Glo Luminescence Viability Assay kit (Promega Corporation, Lyon, France).

#### Cell Staining Assay

The viability and cell count assay were conducted following the protocol recommended by the supplier (Chemometec). Cells were trypsinized to detach them from the culture surface and then resuspended in a suitable medium. A cell suspension was loaded into the Via2-Cassette™ tip for analysis. Inside the cassette, the cells were stained with two fluorescent dyes: AO, which stains all nucleated cells, and DAPI, which selectively stains non-viable cells. Samples were analyzed using an image-based cytometer, the NucleoCounter® NC-3000™ (Chemometec), utilizing the “Viability and Cell Count Assay” program.

#### ATP Measurement

The CellTiter-Glo Luminescence Viability Assay kit was used to quantify cell viability according to the manufacturer’s instructions. Briefly, the CellTiter-Glo reagent was added directly to the cell culture. This reagent lyses the cells and generates a luminescent signal through a reaction between luciferase and oxyluciferin, with the conversion to oxyluciferin being catalyzed by ATP produced by living cells. The resulting luminescent signal was captured and quantified to measure the amount of ATP present, which is directly correlated with the number of viable cells. Luminescence was measured using a Varioskan LUX (Thermo Fisher, Illkirch Graffenstaden, France). Results were presented as dose-response inhibition curves, illustrating the relationship between treatment concentration and cell viability.

### Apoptosis Assay

To assess apoptosis, cells were harvested and counted to achieve a concentration of 2 × 10^5^ cells per sample. Following centrifugation, the cells were resuspended and stained with 100 µL of Annexin V binding buffer, 2 µL of Hoechst 33342, and 2 µL of Annexin V-CF488A conjugate. After incubation (at 37°C for 15 minutes), the cells were washed twice to remove excess dye. Subsequently, they were stained again with 100 µL of Annexin V binding buffer and 2 µL of Propidium Iodide to differentiate between viable and non-viable cells. The samples were analyzed using an image-based cytometer, the NucleoCounter® NC-3000™ (Chemometec), employing the “Annexin V Assay” program. The analysis provided the percentage of cells exhibiting apoptosis, which included both early and late apoptotic stages. The results were plotted to visualize the apoptotic response of the treated cells.

### Chondrosarcoma xenograft model in nude mice

Animal research was conducted in compliance with the current European Directive (2010/63/EU), which has been incorporated into national legislation (Decree 87/848). All animal experiments adhered to internationally accepted ethical principles for laboratory animal use and care to minimize suffering, and all procedures were approved by the Ministry of Education and Research as well as the regional ethics committee (CENOMEXA, France; APAFIS number #37292).

Twenty-two NMRI nude mice (6 weeks old, males), obtained from Janvier Labs (Le Genest-Saint-Isle, France), were housed in ventilated cages under controlled conditions. The environment was specific pathogen-free, with a 12-hour reversed light-dark cycle and a temperature maintained at 23 ± 2°C at the Centre Universitaire de Ressources Biologiques (CURB, Caen, France; accreditation number: C14118015). Mice had ad libitum access to food and water.

Subcutaneous xenografts were established by injecting 1 million JJ012 cells suspended in 150 µL of Matrigel (Fisher Scientific, Illkirch, France) into the right flank of each mouse. When tumors were palpable (*i.e.* 19 days post-implantation), the mice were then divided into three groups for treatment: 5 mice treated with DMSO (vehicle control); 10 mice treated with cladribine at a dose of 20 mg/kg; and 7 mice treated with clofarabine at a dose of 20 mg/kg. Treatment was administered via intraperitoneal injection three times a week. Tumor dimensions were regularly measured using calipers, and tumor volume was calculated using the formula: (L x w^2^)/2, where L is the length and w is the width of the tumor. After 42 days, mice were euthanized by cervical dislocation following inhalation of 5% CO_2_. Four mice previously treated with clofarabine were kept alive for an additional 10 days post-treatment to monitor tumor regrowth. Tumors were subsequently collected, measured, photographed, and weighed.

### Proteomic experiment

#### Sample preparation and analysis

Total protein extraction were performed using RIPA buffer as previously described [27]. The buffer consisting of 50 mM Tris-HCl, 1% Igepal, 150 mM NaCl, 1 mM EGTA, 1 mM NaF, and 0.25% sodium deoxycholate. was supplemented with protease inhibitors to prevent protein degradation during the extraction process. Samples were transferred to Proteogen platform (UNICAEN, Caen) for proteomic experiment.

Five µg of each protein extract were prepared using a modified Gel-aided Sample Preparation protocol [28]. Samples were digested with trypsin/Lys-C overnight at 37°C. For nano-LC fragmentation, peptide samples were first desalted and concentrated onto a µC18 Omix (Agilent) before analysis.

The chromatography step was performed on a NanoElute (Bruker Daltonics) ultra-high-pressure nano flow chromatography system. Approximatively 100ng of each peptide sample were concentrated onto a C18 pepmap 100 (5mm x 300µm i.d.) precolumn (Thermo Scientific) and separated at 50°C onto a reversed phase Reprosil column (25cm x 75μm i.d.) packed with 1.6μm C18 coated porous silica beads (Ionopticks). Mobile phases consisted of 0.1% formic acid, 99.9% water (v/v) (A) and 0.1% formic acid in 99.9% ACN (v/v) (B). The nanoflow rate was set at 250 nL/min, and the gradient profile was as follows: from 2 to 30% B within 70 min, followed by an increase to 37% B within 5 min and further to 85% B within 5 min and reequilibration.

MS experiments were carried out on an TIMS-TOF pro mass spectrometer (Bruker Daltonics) with a modified nano electrospray ion source (CaptiveSpray, Bruker Daltonics). A 1400 spray voltage with a capillary temperature of 180°C was typically employed for ionizing. MS spectra were acquired in the positive mode in the mass range from 100 to 1700 m/z and 0.60 to 1.60 1/k0 window. In the experiments described here, the mass spectrometer was operated in PASEF DIA mode with exclusion of single charged peptides. The DIA acquisition scheme consisted of 16 variable windows ranging from 300 to 1300 m/z.

#### Protein identification

Database searching and LFQ quantification (using XIC) was performed using DIA-NN (version 1.8.2) [56]. An updated UniProt Homo sapiens database was used for library-free search / library generation. For RT prediction and extraction mass accuracy, the default parameter 0.0 was used, which means DIA-NN performed automatic mass and RT correction. Top six fragments (ranked by their library intensities) were used for peptide identification and quantification. The variable modifications allowed were as follows: Nterm-acetylation and Oxidation (M). In addition, C-Propionoamide was set as fix modification. “Trypsin/P” was selected. Data were filtering according to a FDR of 1%. Cross-run normalisation was performed using RT-dependent.

#### Identification of differentially expressed proteins

To quantify the relative levels of protein abundance between different groups, data from DIA-NN were then analysed using DEP package from R. Briefly, proteins that are identified in 2 out of 3 replicates of at least one condition were filtered, missing data were imputed using a normal distribution from PERSEUS and differential enrichment analysis was based on linear models and empherical Bayes statistic. A 1.2-fold increase in relative abundance and a 0.05 false discovery rate (FDR) were used to determine enriched proteins. ANOVA were performed from PERSEUS to determine those enriched proteins.

### Functional Annotation and Pathway Enrichment Analysis

Pathway enrichment analysis and functional annotation were conducted using ShinyGO (v0.8), an online bioinformatics tool available at http://bioinformatics.sdstate.edu/go/ [29]. The analysis encompassed Gene Ontology (GO) functional annotation, including Biological Processes (BP) and Molecular Functions (MF). Additionally, pathways were analyzed using Reactome, WikiPathways, and KEGG databases. Enrichment p-values were calculated using the hypergeometric test, and to account for multiple testing, FDR was computed using the Benjamini-Hochberg method. Data were analyzed by using the “sort by average ranks (FDR & Fold Enrichment)” option permitting to initially filter pathways and GO terms by an FDR threshold below 0.05, and only those deemed significant were subsequently ranked. The top 15 pathways were selected for further analysis and discussion.

### Cell Cycle Analysis

Cell cycle distribution was assessed using the NucleoCounter® NC-3000™ (Chemometec) in accordance with the manufacturer’s “Fixed Cell Cycle-DAPI Assay” protocol.

A total of 18,000 cells per cm² were seeded and incubated for 24 hours with the respective treatments. Following this incubation, both floating and adherent cells (which were harvested via trypsinization) were combined and washed with phosphate-buffered saline (PBS). Next, approximately 2 x 10⁶ cells were fixed in ice-cold 70% ethanol for a minimum of 24 hours at 4°C. After fixation, the cells were washed again with cold PBS. The cell pellet was then resuspended in 0.5 mL of DAPI solution (comprising 1 µg/mL DAPI and 0.1% Triton X-100 in PBS) and incubated for 5 minutes at 37°C. The percentages of cells arrested in the sub-G1, G0/G1, S, and G2/M phases of the cell cycle were quantified using the manufacturer’s software.

### Statistics analysis

All statistical analyses were conducted using Prism version 10.1.2. Data from both *in vitro* and *in vivo* experiments are presented as mean ± SEM. Statistical comparisons were made using a two-way ANOVA, with Dunnett’s or Sidak’s multiple comparison tests as appropriate. For tumor weight analysis, a t-test was employed. Statistical significance was defined as a p-value of less than 0.05.

## Results

### Cytotoxic Effects of Adenosine Analogues in SW1353 Chondrosarcoma Cell Line

To evaluate the efficacy of various adenosine analogues, we first investigated their cytotoxic effects on the widely studied chondrosarcoma cell line SW1353. Cells were treated with increasing concentrations of aristeromycin, cladribine, formycin, or pentostatin for a period of seven days. Microscopic examination of treated SW1353 cells revealed significant morphological changes, including reduced cell density and noticeable cell shrinkage at 1 µM concentrations of aristeromycin, cladribine, and formycin compared to control cells (figure 2A).

**Figure 2:**
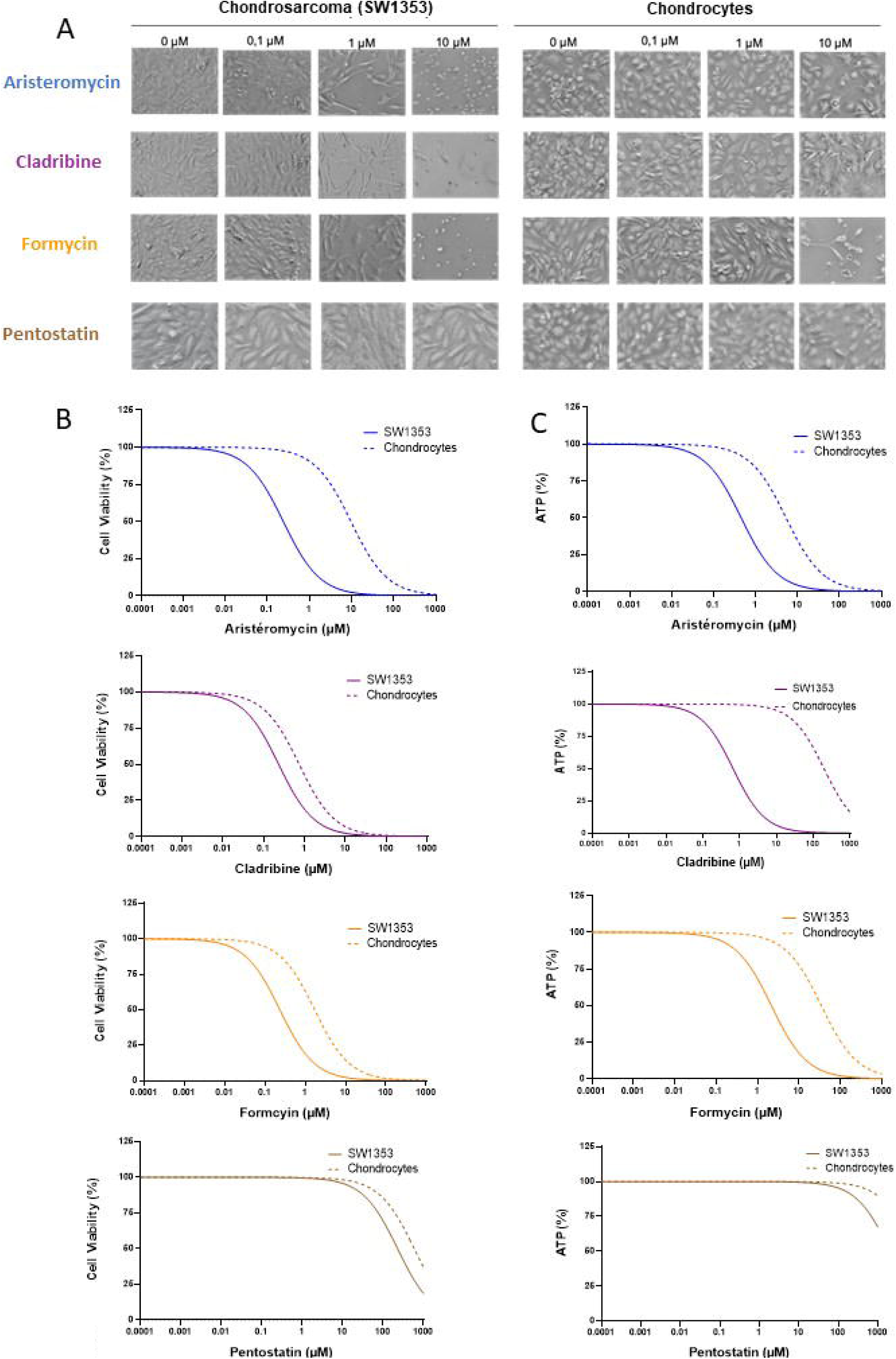
Cytotoxic Effects of Adenosine Analogues in SW1353 Chondrosarcoma Cell Line. SW1353 chondrosarcoma cells and chondrocytes were treated with increasing doses of aristeromycin, cladribine, formycin, or pentostatin (0 to 1000 µM). After 7 days of treatment, microscopic observations were performed (A), and cell viability was assessed by counting viable cells (B) or measuring ATP levels (C). Experiments were independently repeated at least 3 times for each cell line and treatment. Graphs represent the mean of the experiments.

Cell viability was further assessed using acridine orange and DAPI staining. This analysis demonstrated that aristeromycin, cladribine, and formycin decreased cell viability in a concentration-dependent manner in the SW1353 line (figure 2B). In contrast, primary chondrocytes exhibited significantly less sensitivity to these three analogues. ATP quantification using the CellTiter reagent corroborated the cytotoxic effects of aristeromycin, cladribine, and formycin on chondrosarcoma cells (figure 2C).

In contrast, pentostatin did not induce any cytotoxic effects in either chondrosarcoma cells or chondrocytes after seven days of treatment.

### Efficacy of Adenosine Analogues Across Multiple Chondrosarcoma Cell Lines

To better represent tumor heterogeneity, we tested the three effective adenosine analogues (aristeromycin, cladribine, and formycin) on three additional human chondrosarcoma lines: JJ012, CH2879, and FS090. After 48 hours of treatment, the cells were stained with acridine orange and DAPI, and cell viability was quantified (figure 3). As anticipated, aristeromycin, cladribine, and formycin effectively reduced viability in SW1353 without affecting chondrocyte viability. Importantly, similar responses were observed in the other chondrosarcoma cell lines. Specifically, treatment with aristeromycin (10 µM), cladribine (1 µM), or formycin (1 µM) resulted in reduced viability across all four chondrosarcoma lines tested.

**Figure 3:**
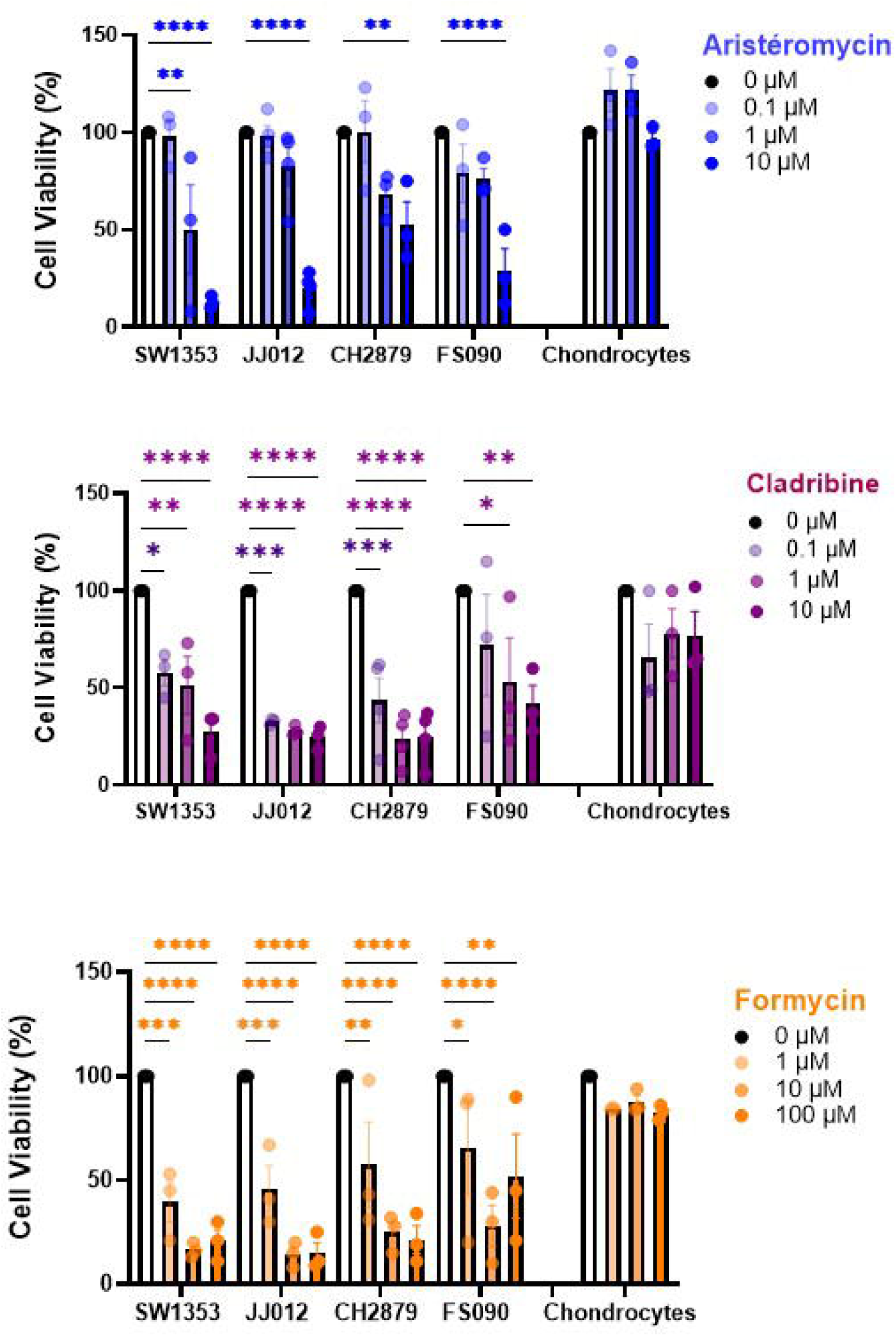
Efficacy of Adenosine Analogues Across Multiple Chondrosarcoma Cell Lines. Four chondrosarcoma cell lines (SW1353, JJ012, FS090, and CH2879) and chondrocytes were treated with increasing concentrations of aristeromycin, cladribine, or formycin. After 2 days of treatment, cell viability was assessed by counting viable cells. The results of three independent experiments are shown. Histograms represent the percentage of live cells after treatment compared to controls. Data are expressed as means ± SEM. *: p-value < 0.05, **: p-value < 0.01, ***: p-value < 0.001, ****: p-value < 0.0001.

### Apoptosis Induced by Aristeromycin and Cladribine

Given the comparable effects previously observed, we opted to focus subsequent analyses on the two most common chondrosarcoma lines: SW1353 and JJ012. To investigate whether the observed decrease in cell viability resulted from apoptotic cell death, we exposed the chondrosarcoma cell lines, as well as chondrocytes, to the three adenosine analogues for 48 hours. Apoptosis was quantified using Annexin V and Propidium Iodide staining. Both aristeromycin and cladribine significantly increased apoptosis in SW1353 and JJ012 cells (figures 4A and B). In contrast, formycin did not induce apoptosis in chondrosarcoma cell lines, suggesting that its cytotoxicity may be mediated through a different mechanism. Importantly, there was no increase in apoptosis observed in chondrocytes regardless of the analogue tested.

**Figure 4:**
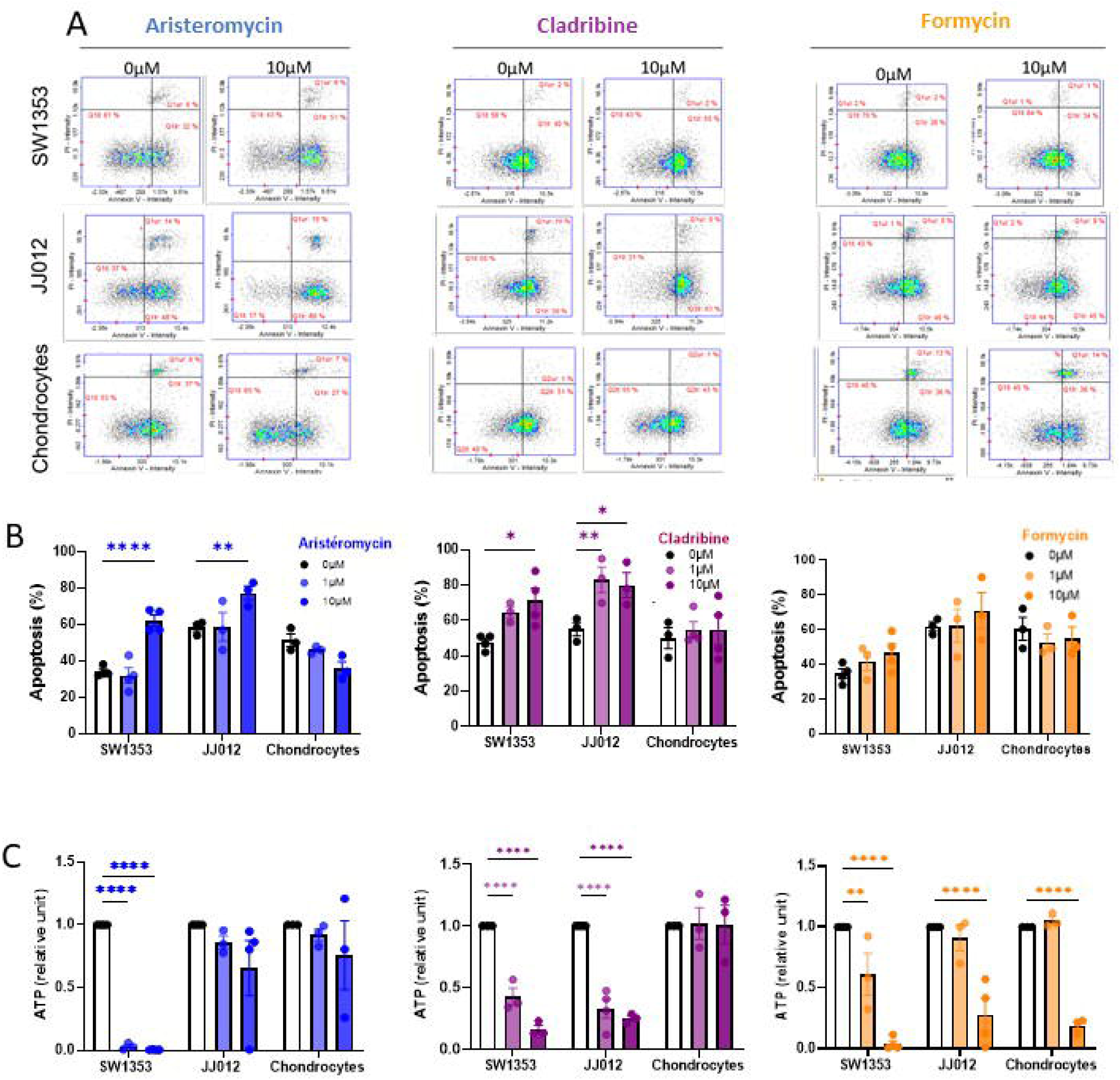
Apoptosis Induced by Aristeromycin and Cladribine and Efficacy in 3D Chondrosarcoma Models. SW1353 and JJ012 chondrosarcoma cells and chondrocytes were treated with aristeromycin, cladribine, or formycin (0, 1, or 10 µM). After 2 days of treatment, apoptotic cells were identified by Annexin V and Propidium Iodide staining. Cytograms show a representative result from three or four independent experiments (A). Histograms represent the percentage of apoptotic cells (early and late) obtained (B). Alginate beads containing JJ012 or SW1353 chondrosarcoma cells, or chondrocytes, were treated with aristeromycin, cladribine, or formycin (0, 1, or 10 µM) 15 days after formation. After 7 days of treatment, cell viability was assessed by ATP measurement (C). The results of three independent experiments are shown. Data are expressed as means ± SEM. *: p-value < 0.05, **: p-value < 0.01, ****: p-value < 0.0001.

### Cladribine’s Efficacy in 3D Chondrosarcoma Models

To enhance the translational relevance of our findings, we evaluated the effects of aristeromycin, cladribine, and formycin in a 3D alginate bead model that more closely mimics the tumor microenvironment [58]. After 14 days of culture, alginate beads were treated with the adenosine analogues for an additional seven days (figure 4C). Cladribine was the only analogue to significantly reduce viability in both chondrosarcoma cell lines without affecting chondrocytes. Aristeromycin showed a reduction in viability exclusively in the SW1353 line, while formycin reduced viability across all tested cell types, including chondrocytes. These findings prompted us to focus further investigations on cladribine.

### Proteomic Analysis Following Cladribine Treatment

To elucidate the mechanisms underlying cladribine’s action, we conducted a proteomic study. After 48 hours of treatment, 36 proteins were found to be deregulated in the SW1353 cell line (FDR < 0.05), with 12 downregulated (33.3%) and 24 upregulated (66.7%) proteins identified (figure 5A, suppl table 1). In the JJ012 cell line, a total of 1,275 proteins were deregulated after cladribine treatment (FDR < 0.05), of which 938 were highly significant (FDR < 0.001). This included 685 downregulated (53.8%) and 590 upregulated (46.2%) proteins (suppl fig 1A and suppl table 1).

**Figure 5:**
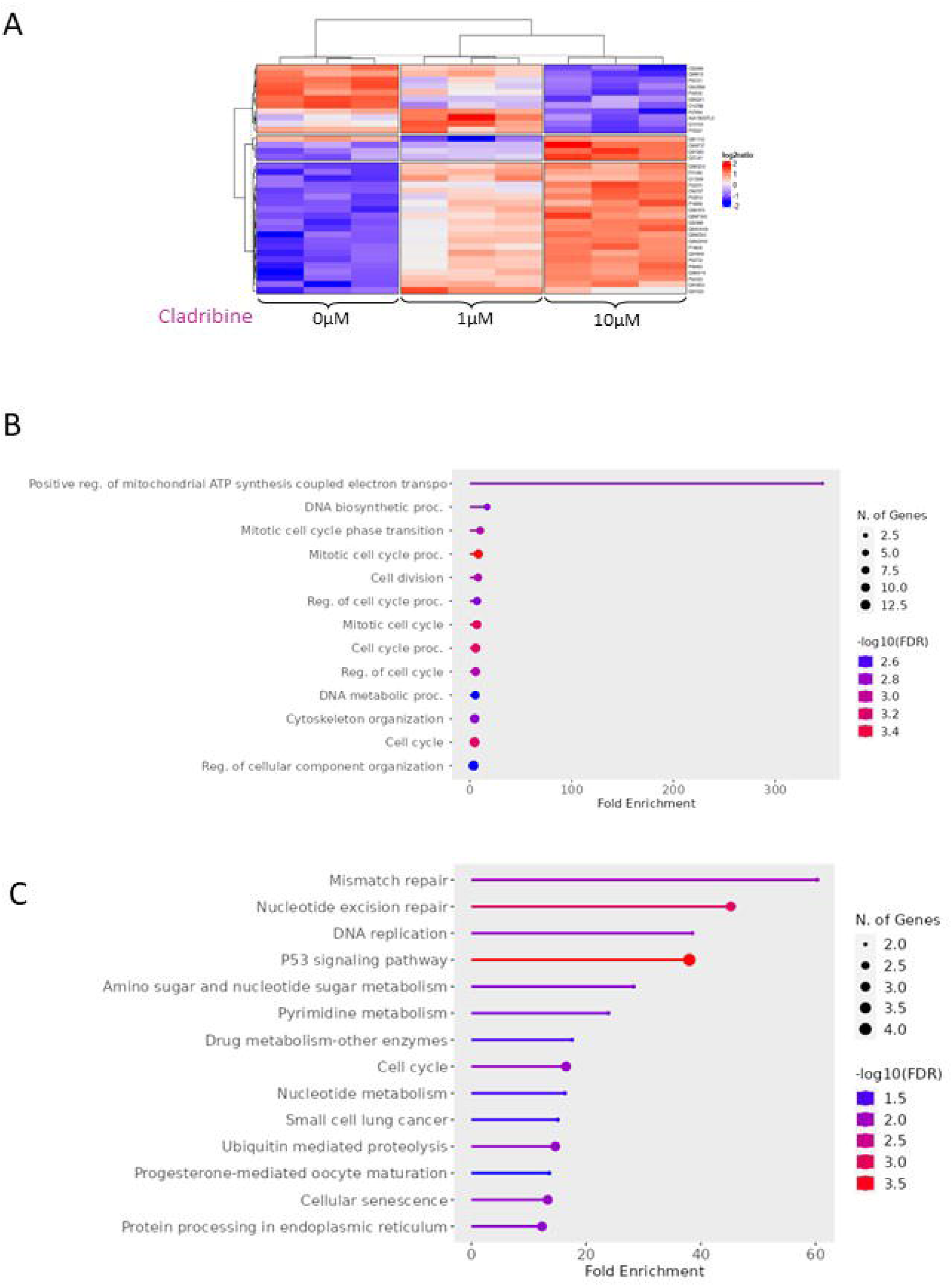
Proteomic Analysis Following Cladribine Treatment in SW1353. Heatmap showing the effects of cladribine treatment (0, 1, or 10 µM) on the SW1353 chondrosarcoma cell line after 48 hours (A). Gene enrichment analysis was performed using ShinyGO. The bubble plot represents the top 15 significantly enriched terms for deregulated proteins in chondrosarcoma cells after 48 hours of cladribine treatment. Gene Ontology (GO) terms are shown for Biological Process (BP) (B) and KEGG pathways (C). An FDR < 0.05 was considered significant, and fold enrichment scores were used to evaluate the relevance of GO terms and the enriched pathways.

Analysis of these deregulated proteins revealed significant enrichment in pathways related to the cell cycle, purine metabolism, and overall metabolism, as identified through GO Biological Process and KEGG database analyses (figure 5B-C, suppl fig 1B-C and suppl tables 2 and 3). Additional evaluations using GO term Molecular Function, Reactome, and WikiPathways databases confirmed enrichment in cell cycle and metabolic pathways, including ribosomal and ribonucleoprotein activities, as well as pathways associated with DNA repair and the p53 signaling pathway, often linked to cell death (suppl tables 4 and 5). Collectively, these results suggest that cladribine may induce apoptosis by disrupting the cell cycle and influencing the p53 pathway.

### Alteration of Cell Cycle Induced by Cladribine

Given the proteomic findings indicating a potential effect on the cell cycle, we examined the various phases of the cell cycle in SW1353 and JJ012 cell lines following 48 hours of cladribine treatment. In SW1353, cladribine treatment resulted in an increase in S-phase cells and a decrease in G2/M-phase cells, indicating S-phase cell cycle arrest (figure 6). Conversely, in JJ012, cladribine treatment led to an increase in G1-phase cells and a decrease in G2/M-phase cells, suggesting either a block in the transition from G1 to S phase or an acceleration of the transition from G2/M to G1 phase (Suppl figure 2).

**Figure 6:**
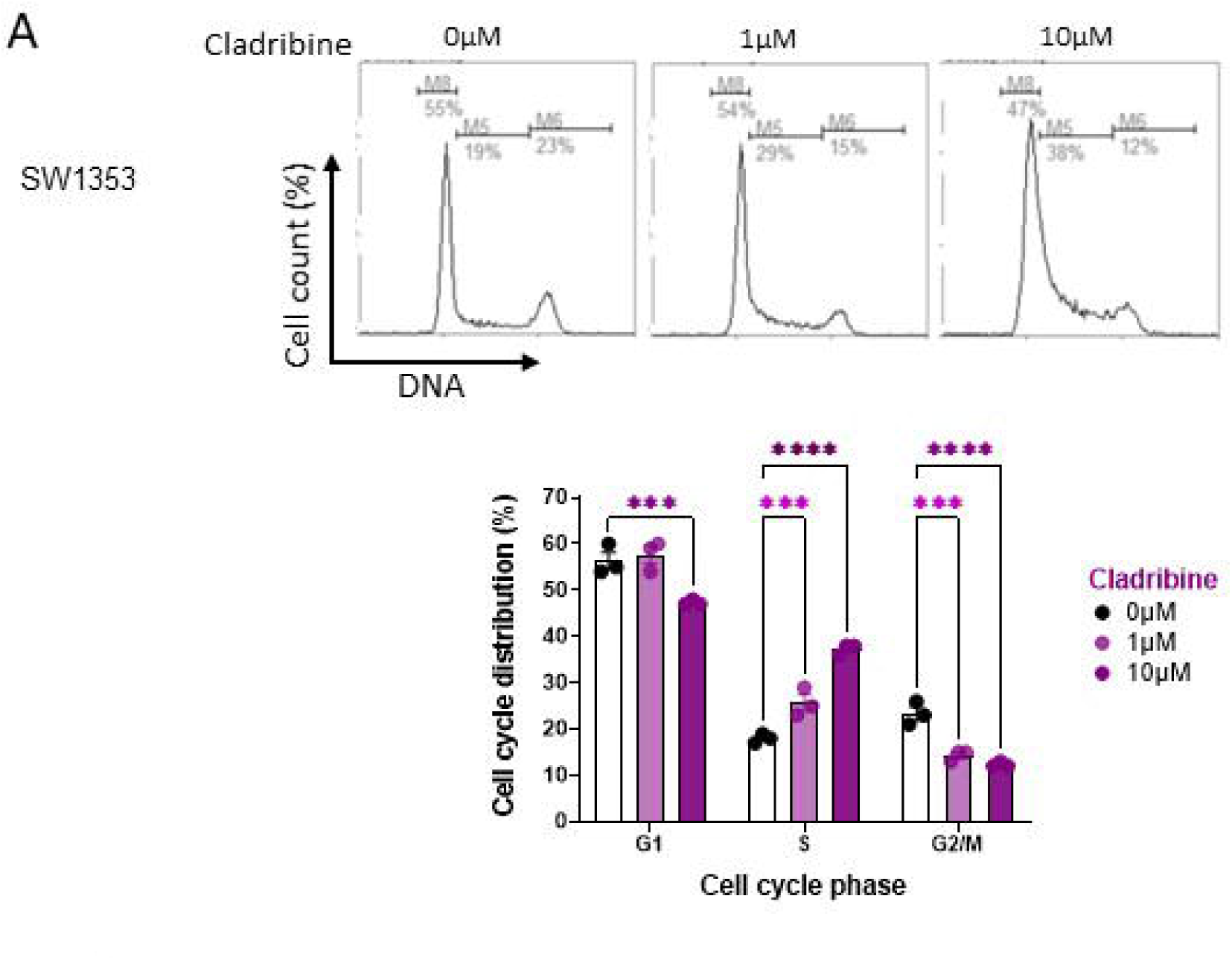
Alteration of Cell Cycle Induced by Cladribine. After 24 hours of treatment with cladribine (0, 1, or 10 µM), SW1353 cells were harvested, and the cell cycle was analyzed using image-based cytometry. Cytograms show a representative result from three independent experiments. Histograms represent the distribution of cells across the different cell cycle phases. Data are expressed as means ± SEM. *: p-value < 0.05, **: p-value < 0.01.

### In Vivo Efficacy of Cladribine in Tumor Growth Reduction

To assess the *in vivo* efficacy of cladribine, we established heterotopic chondrosarcoma xenografts using JJ012 cells in nude mice. Once tumors were palpable, cladribine (20 mg/kg) or vehicle (DMSO) were intraperitoneally administrated three times per week for six weeks in mice (figure 7A). The treatment was well-tolerated, with no significant reductions in body weight observed (Suppl fig 3A) or signs of discomfort. Cladribine significantly slowed tumor growth, as evidenced by decreased tumor volumes compared to vehicle-treated mice (figure 7B). Additionally, tumor weight was significantly reduced in cladribine-treated mice (figure 7C).

**Figure 7:**
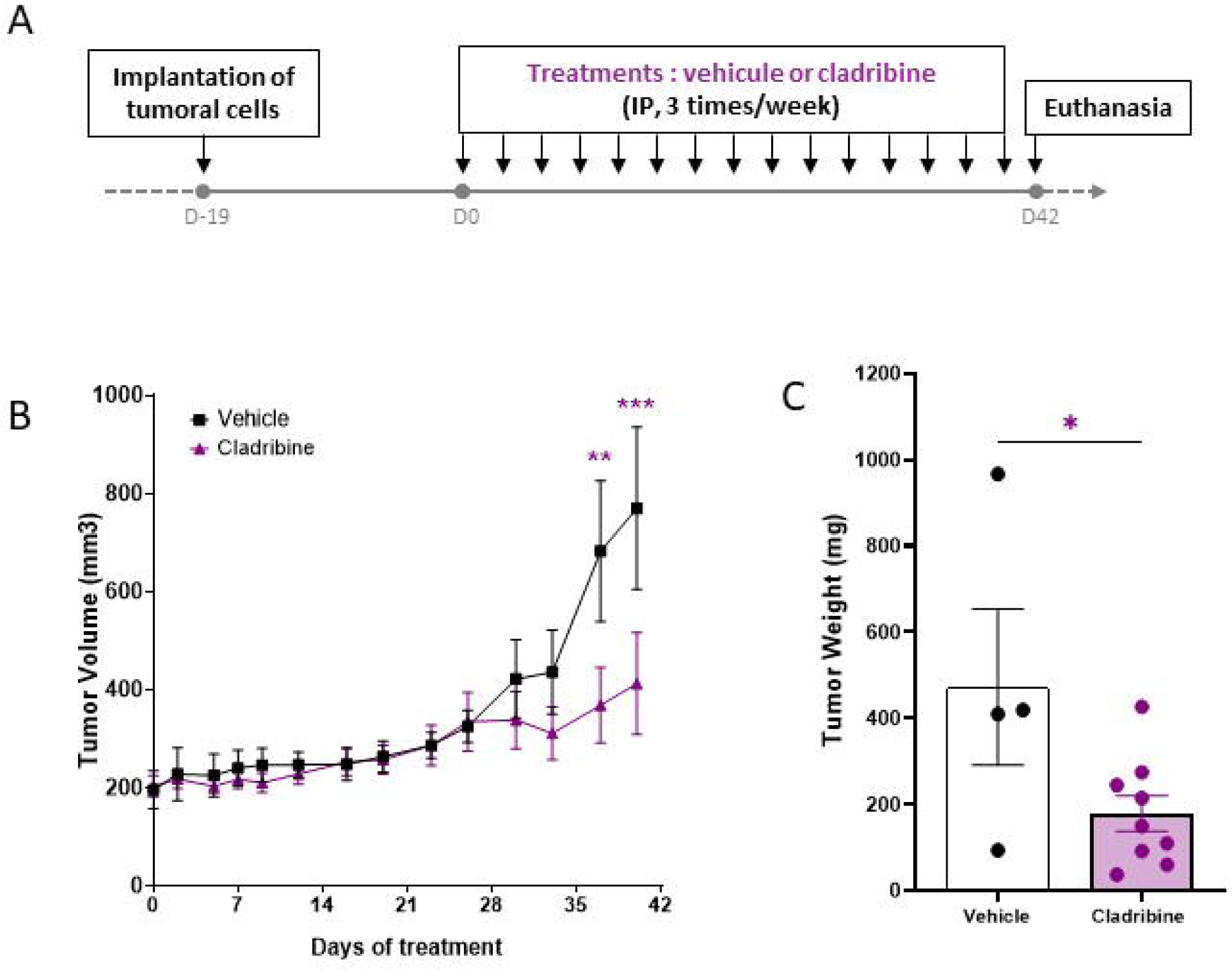
In Vivo Efficacy of Cladribine in Tumor Growth Reduction. A xenograft of JJ012 chondrosarcoma cells was subcutaneously implanted into nude mice, which were treated intraperitoneally with 20 mg/kg cladribine (n=10) or vehicle (n=5), three times per week for 42 days (A). Tumor volume was measured three times per week (B). After 42 days, tumors were collected and weighed (C). Data are expressed as means ± SEM. **: p-value < 0.01, ***: p-value < 0.001.

### Evaluation of Antitumor Effects of Clofarabine, a Cladribine Derivative, in chondrosarcomas

Next, we were interested in a derivative of cladribine, the clofarabine, that has demonstrated enhanced efficacy in Ewing’s sarcoma mouse models [30].

*In vitro*, clofarabine significantly decreased the viability of chondrosarcoma cells in both monolayer and 3D cultures without affecting the viability of chondrocytes. Furthermore, clofarabine induced apoptosis in chondrosarcoma cells (figure 8).

**Figure 8:**
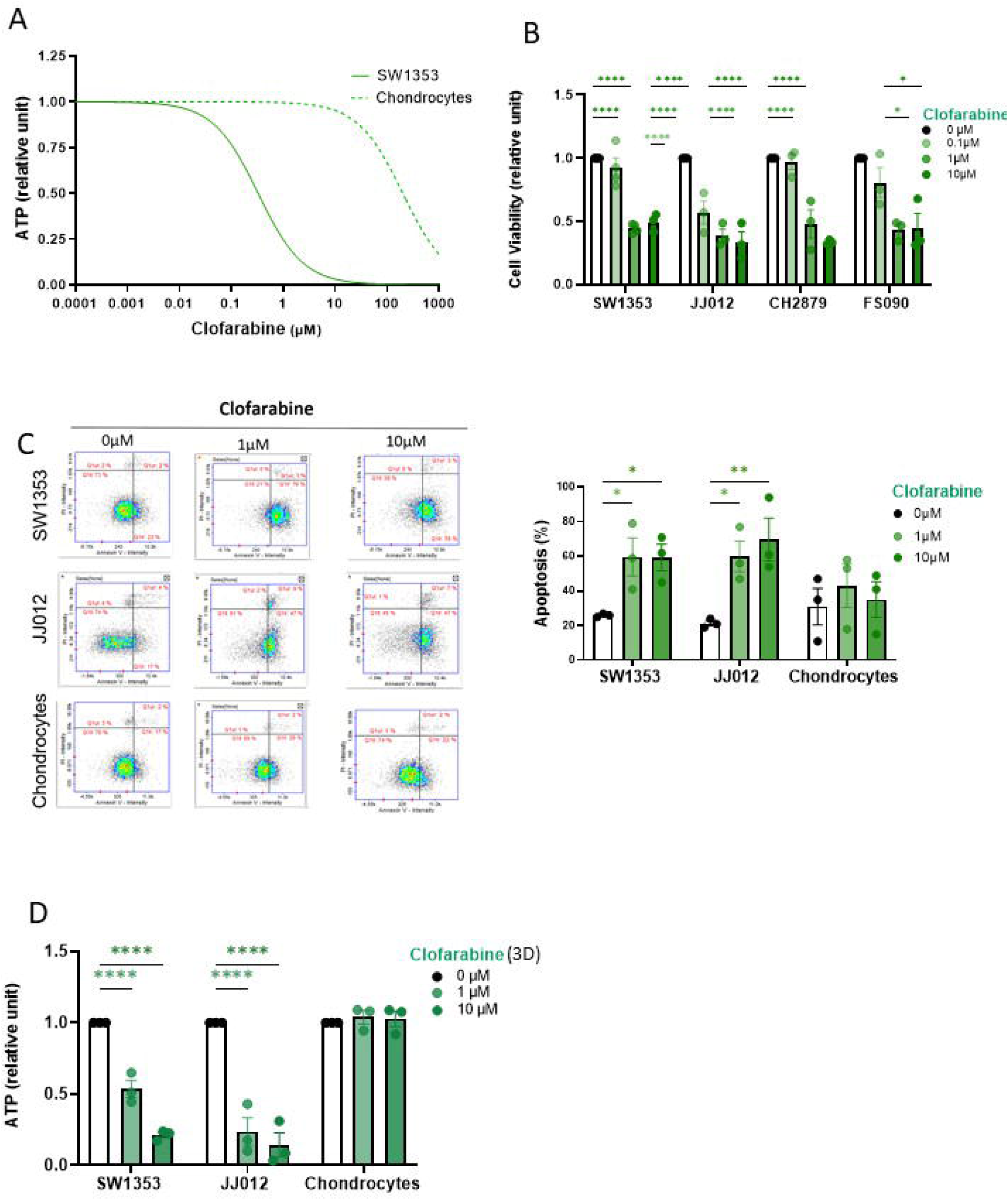
*In Vitro* Evaluation of Antitumor Effects of Clofarabine, a Cladribine Derivative, in chondrosarcomas. Chondrosarcoma cell lines and chondrocytes were treated with increasing doses of clofarabine. Cell viability was assessed after 7 days of treatment by ATP measurement using the Celltiter-Glo assay (A), or after 2 days of treatment by counting viable cells (B). Apoptosis was determined by Annexin V and Propidium Iodide staining after 48 hours of treatment. Cytograms show a representative result from three independent experiments, and histograms represent the percentage of apoptotic cells (early and late) observed (C). Additionally, alginate beads containing chondrosarcoma cells or chondrocytes were treated with clofarabine (0, 1, or 10 µM), and cell viability was evaluated after 7 days of treatment by ATP measurement (D). Data are expressed as means ± SEM. *: p-value < 0.05, **: p-value < 0.01, ***: p-value < 0.001, ****: p-value < 0.0001.

Furthermore, we assessed the antitumor effects of clofarabine in a chondrosarcoma xenograft model in nude mice. Following JJ012 cell implantation, mice received either vehicle (DMSO) or clofarabine (20 mg/kg) intraperitoneally (figure 9A). Mice tolerated the treatment well, with no significant changes in body weight (suppl fig 3B), and interestingly, clofarabine reduced tumor growth. Tumor volume in clofarabine-treated mice remained stable throughout the 42-day treatment period and continued to be maintained for up to 10 days after treatment cessation (figure 9B). Moreover, clofarabine treatment tended to decrease tumor weight compared to vehicle treatment, with final weights recorded at 100.7 ± 34.8 mg for clofarabine-treated mice and 473 ± 181 mg for vehicle-treated mice (p = 0.07; figure 9C).

**Figure 9:**
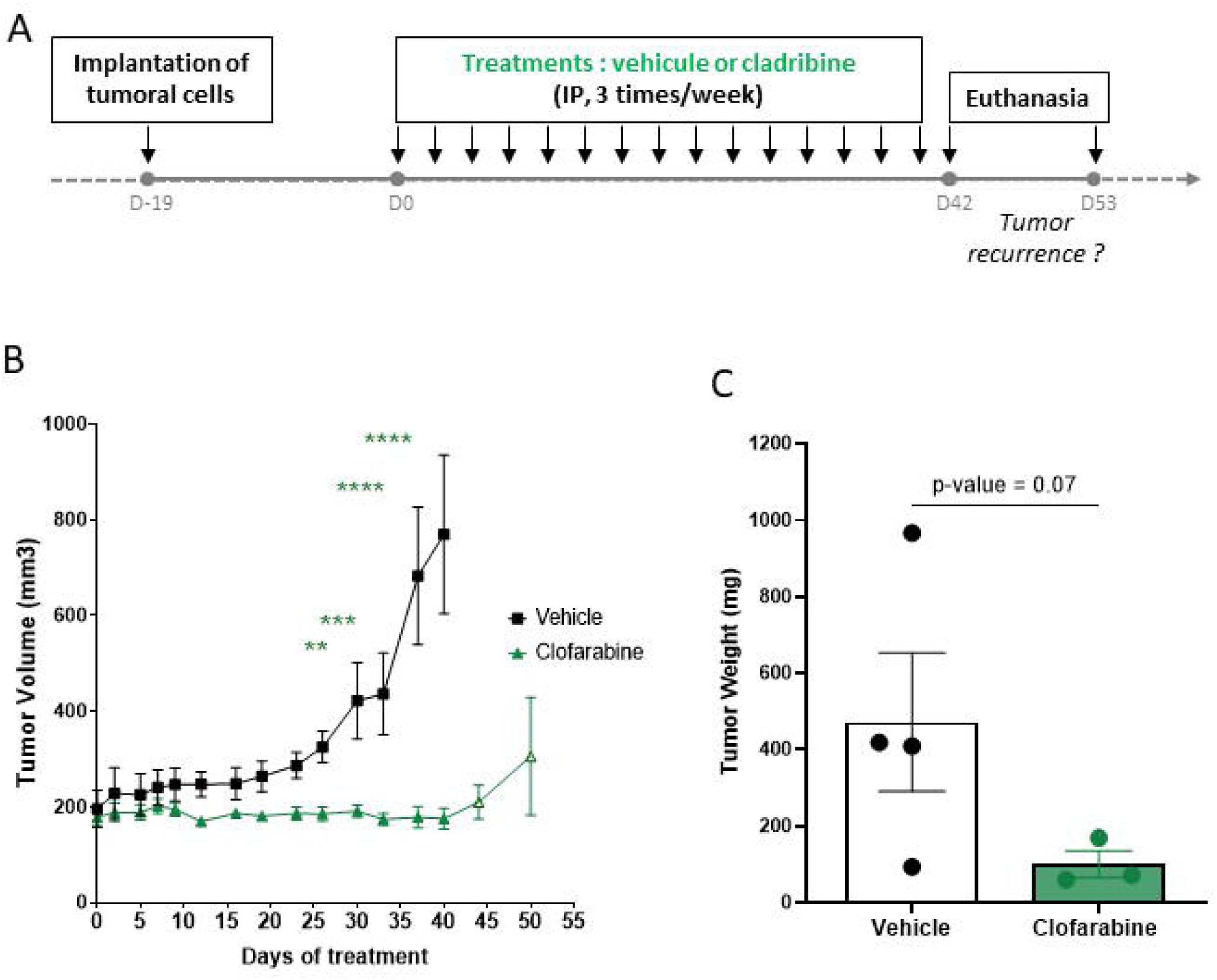
*In Vivo* Evaluation of Antitumor Effects of Clofarabine. JJ012 chondrosarcoma cells were implanted as subcutaneous xenografts into nude mice. Once tumors became palpable, intraperitoneal injections of 20 mg/kg clofarabine (n = 7) or vehicle (n = 5) were administered three times per week for 40 days. Tumor volume was measured with a caliper three times per week throughout the treatment period (B). On day 42, four mice treated with clofarabine were kept alive and monitored for an additional 10 days after treatment cessation to evaluate potential tumor recurrence. The remaining mice were sacrificed, and tumors were weighed. **: p-value < 0.01, ***: p-value < 0.001, ****: p-value < 0.0001.

## Discussion

Prior studies have demonstrated that the adenosine analogue 3-dezaneplanocin (DZNep) effectively induces apoptosis both *in vitro* and *in vivo* in chondrosarcoma models. However, DZNep is not available for clinical use, posing significant challenges for human application in terms of time and rentability/cost. In this study, we identified two additional adenosine analogues —cladribine and clofarabine— that are currently utilized clinically for other malignancies. Both analogues significantly decreased chondrosarcoma viability *in vitro* and in vivo, promoted apoptotic activity, and induced cell cycle arrest, positioning them as promising candidate drugs for the treatment of patients with chondrosarcomas.

In our i*n vitro* studies, cladribine and clofarabine emerged as adenosine analogues capable of effectively reducing cell viability and inducing apoptosis in all tested chondrosarcoma cell lines, while sparing normal chondrocytes. In contrast, other analogues either failed to induce apoptosis across all chondrosarcoma lines or exhibited toxicity towards chondrocytes. The variability in the efficacy of these adenosine analogues can be attributed to several factors. For instance, it is noteworthy that pentostatin did not reduce survival in the SW1353 cell line, despite its proven effectiveness in Hairy cell leukemia, where it is reported to have similar efficacy to cladribine [31,32]. Pentostatin specifically inhibits adenosine deaminase (ADA), while cladribine is described as ADA-resistant. Nevertheless, both agents result in the accumulation of deoxyadenosine triphosphate (dATP), leading to DNA strand breaks and the activation of apoptotic pathways via p53 and cytochrome c release from mitochondria. The enhanced efficacy of cladribine can be attributed to its additional mechanism of action: its active triphosphorylated form (CdATP) integrates into DNA, inhibiting DNA polymerase β and disrupting DNA repair processes. This further DNA damage activates poly(ADP-ribose) polymerase (PARP), resulting in the depletion of cellular nicotinamide adenine dinucleotide (NAD) and ATP [33]. This PARP activation-induced necrosis could explain cladribine’s superior effectiveness in treating chondrosarcoma. While pentostatin-treated cells may evade cell cycle control, those exposed to cladribine experience halted cell cycle progression, rendering them incapable of DNA synthesis or repair, ultimately leading to apoptosis.

Additionally, Uchiyama et al. found that DZNep and its analogue aristeromycin exhibit strong inhibitory activity against SAHH in prostate cancer cells, resulting in reduced cell proliferation [16]. However, unlike DZNep, aristeromycin did not consistently inhibit growth across all tested chondrosarcoma lines, particularly in 3D models. This observation aligns with findings in prostate cancer cell lines, suggesting that aristeromycin’s efficacy may vary significantly [16]. Aristeromycin’s structural similarity to ATP, with a cyclopentyl modification replacing the furanose sugar, could contribute to its increased toxicity, particularly towards non-tumor cells such as chondrocytes, as our results indicate. A similar toxicity profile was noted with formycin, which significantly compromised chondrocyte viability. Consequently, both aristeromycin and formycin may require the development of novel derivatives to reduce their adverse effects while maintaining therapeutic efficacy [21].

In contrast to the previously mentioned analogues, cladribine and clofarabine demonstrated promising results in reducing cell viability by inducing S-phase arrest and blocking the G1/S transition, which in turn promotes apoptosis in chondrosarcomas. This mechanism is consistent with the established effects of these agents in leukemia and Ewing’s sarcoma [5,14,23,30,34,35]. Importantly, both adenosine analogues exhibited selective toxicity, sparing non-tumor cells.

*In vivo*, both cladribine and clofarabine significantly reduced tumor volume. Notably, clofarabine showed superior efficacy. Tumors in clofarabine-treated mice did not increase in size throughout the 42-day treatment period, whereas tumors treated with cladribine, although growing more slowly than those in control groups, still exhibited some growth. Despite this, complete tumor regression was not achieved with clofarabine within the study’s dose and duration. Upon discontinuation of treatment, initial tumor regrowth was observed within ten days, indicating a need for continuous treatment or optimization of the administration protocol.

Both cladribine and clofarabine were administered *via* intraperitoneal injection, a method that may not be optimal. Previous studies have shown that oral administration of clofarabine can be more effective than the same dosage given intraperitoneally [30,36]. Additionally, higher doses of clofarabine might be necessary, as its effects in leukemia blast cells are dose-dependent. For example, higher doses resulted in sustained inhibition of DNA synthesis, whereas lower doses only allowed for partial recovery [37]. In our study, we utilized a dose of 20 mg/kg, which may have been insufficient. Therefore, further efforts are needed to optimize the administration and dosing protocols for clofarabine in the treatment of chondrosarcoma. Additionally, investigating combinations of clofarabine with other cytotoxic agents could yield synergistic effects, enhancing therapeutic outcomes for patients with chondrosarcoma.

In summary, we have identified three adenosine analogues—DZNep (in a previous report), cladribine, and clofarabine—that show efficacy against chondrosarcoma *in vitro* and *in vivo*. These agents hold promise as potential therapeutic options, as they effectively inhibit tumor growth with minimal toxicity. The findings suggest that cladribine, clofarabine and their derivatives could be valuable treatments for chondrosarcoma, a largely underexplored area in oncology. Further research is needed to clarify their molecular mechanisms and evaluate their therapeutic potential in clinical settings.

## Acknowledgment

We thank Sylvain Leclercq, Baptiste Picard and collaborators (Service orthopédique de la Clinique Saint-Martin, Caen France) for the femoral head samples, as well as Pierre Ruggeri (MSc Student, BioConnect, Caen), Julien Pontin (Proteogen, Unicaen, France) and Palma Pro and Charlotte Marie (CURB, Unicaen, France) for technical assistance.

