## Supplementary figures and images for "Exploring Adenosine Analogues for Chondrosarcoma Therapy: In Vitro and In Vivo Insights"

### suppl fig 1

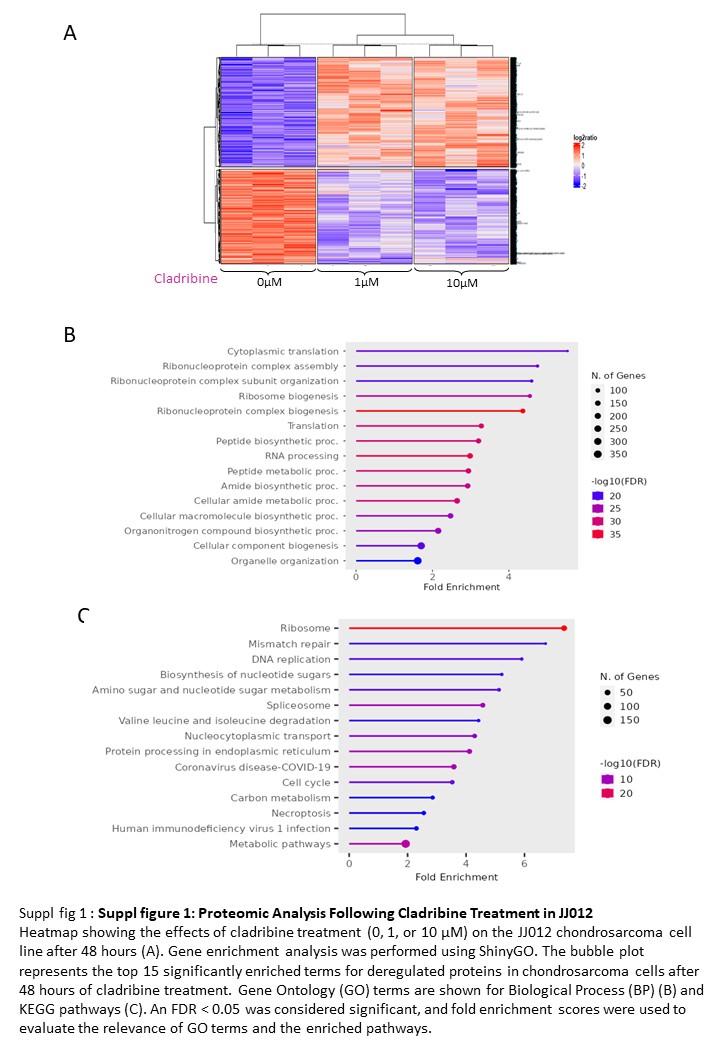

### suppl fig 2

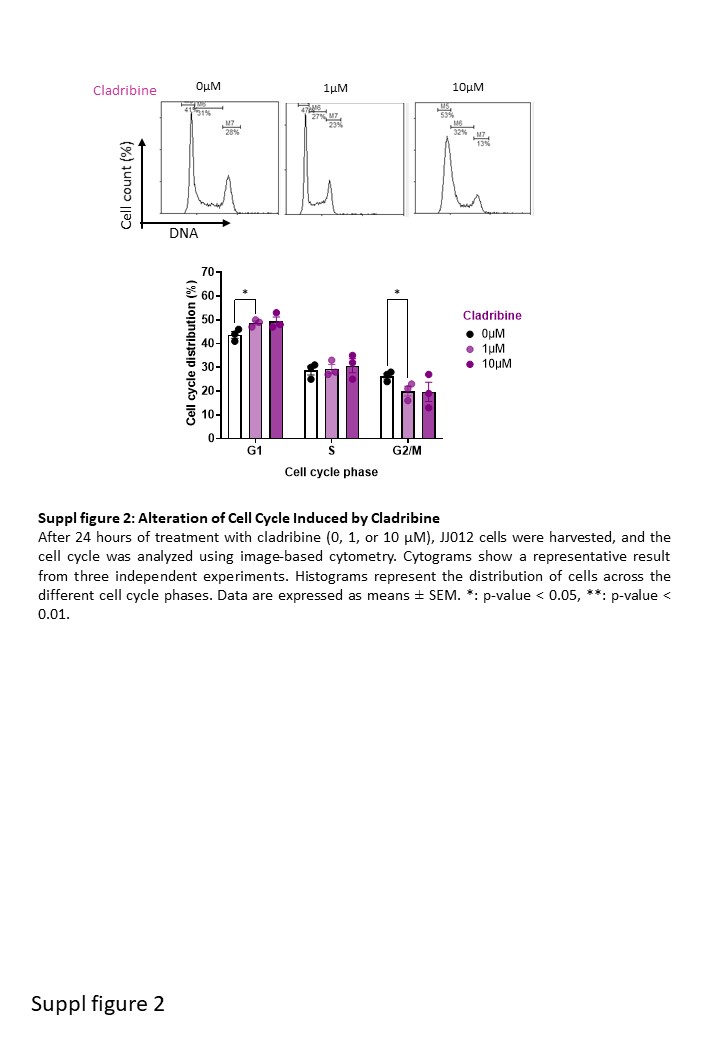

### suppl fig 3

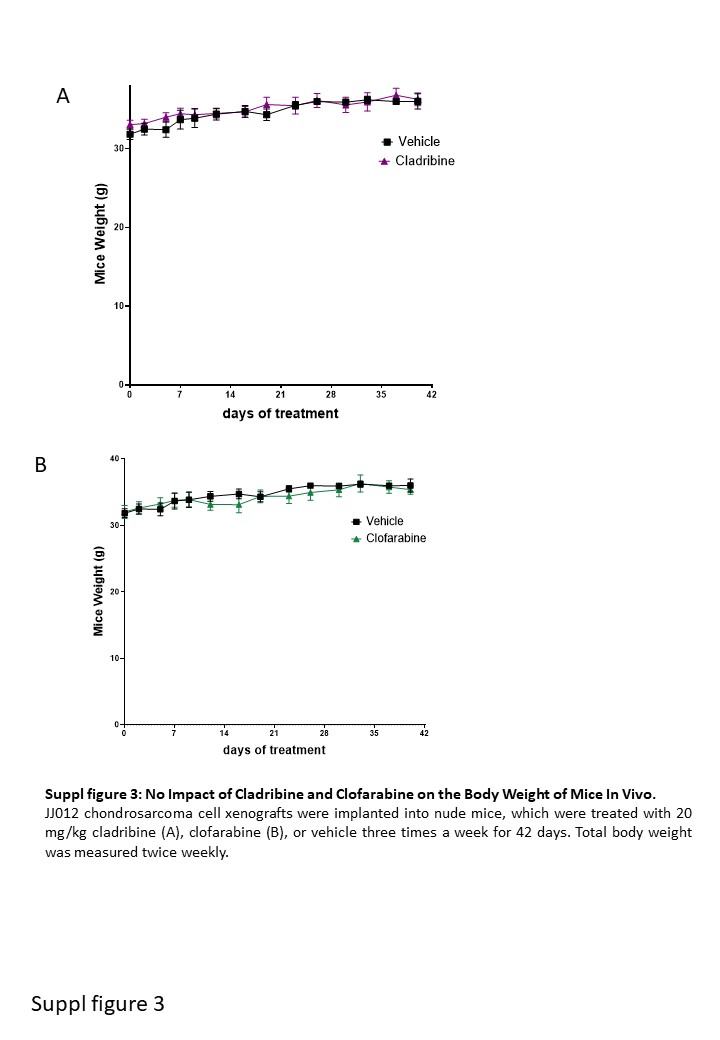
