## Supplementary material for "Exploring Adenosine Analogues for Chondrosarcoma Therapy: In Vitro and In Vivo Insights": suppl table

**Suppl table 1: List of proteins significantly deregulated after cladribine treatment (1 and 10 µM) in cell line SW1353 and JJ012. A 0.05 FDR was used.**

| Cell line | Ids Protein | Genes Name | Ids Protein | Genes Name | Ids Protein | Genes Name | Ids Protein | Genes Name |
|---|---|---|---|---|---|---|---|---|
| **JJ012** | Q8N5N7 | MRPL50 | Q9UBB6 | NCDN | Q9UID3 | VPS51 | Q9NZB2 | FAM120A |
| | O14757 | CHEK1 | O75828 | CBR3 | P10412 | H1-2 | P55786 | NPEPPS |
| | P50443 | SLC26A2 | Q15459 | SF3A1 | Q9BVJ6 | UTP14A | Q96I25 | RBM17 |
| | P49711 | CTCF | Q9P015 | MRPL15 | Q2NL82 | TSR1 | Q9UHN6 | CEMIP2 |
| | P11216 | PYGB | P53618 | COPB1 | P07237 | P4HB | Q9NSU2 | TREX1 |
| | Q9UBE0 | SAE1 | O43488 | AKR7A2 | Q15910 | EZH2 | Q16527 | CSRP2 |
| | P30530 | AXL | P14635 | CCNB1 | P46777 | RPL5 | O00203 | AP3B1 |
| | P43121 | MCAM | O43709 | BUD23 | P54098 | POLG | O94842 | TOX4 |
| | Q32P28 | P3H1 | Q7Z4G1 | COMMD6 | Q9P2E9 | RRBP1 | Q9UBR5 | CKLF |
| | Q9C0D5 | TANC1 | Q15003 | NCAPH | Q02297 | NRG1 | O43684 | BUB3 |
| | P30740 | SERPINB1 | Q9ULW0 | TPX2 | Q4U2R6 | MRPL51 | P25205 | MCM3 |
| | Q9HBF4 | ZFYVE1 | P06493 | CDK1 | P15509 | CSF2RA | Q8TB52 | FBXO30 |
| | P04181 | OAT | O14657 | TOR1B | P53814 | SMTN | P56199 | ITGA1 |
| | Q96PZ2 | FAM111A | P54136 | RARS1 | Q02878 | RPL6 | Q7Z569 | BRAP |
| | Q13885 | TUBB2A | P40763 | STAT3 | O75051 | PLXNA2 | Q9NXF1 | TEX10 |
| | P80723 | BASP1 | Q8WUX1 | SLC38A5 | O00411 | POLRMT | Q99569 | PKP4 |
| | O14640 | DVL1 | Q9UJY1 | HSPB8 | P23229 | ITGA6 | P49902 | NT5C2 |
| | Q9UBI6 | GNG12 | Q12872 | SFSWAP | P62699 | YPEL5 | Q10589 | BST2 |
| | Q13315 | ATM | Q9NY93 | DDX56 | Q70UQ0 | IKBIP | O95831 | AIFM1 |
| | P07305 | H1-0 | P48735 | IDH2 | Q9UMY1 | NOL7 | A0A499FIZ0 | WDR26 |
| | Q9NR45 | NANS | Q15036 | SNX17 | Q96EX3 | DYNC2I2 | Q9UBM7 | DHCR7 |
| | P21912 | SDHB | Q9BZJ0 | CRNKL1 | Q9UJ70 | NAGK | Q99661 | KIF2C |
| | Q9NQ88 | TIGAR | Q14331 | FRG1 | Q96M96 | FGD4 | A0A1B0GW05 | DPY19L1 |
| | Q96TA2 | YME1L1 | P04818 | TYMS | O15118 | NPC1 | Q5JTJ3 | COA6 |
| | Q9NXG2 | THUMPD1 | Q02241 | KIF23 | Q9UHB6 | LIMA1 | Q9BYD2 | MRPL9 |
| | Q6NUQ4 | TMEM214 | Q9ULX3 | NOB1 | Q99707 | MTR | O60869 | EDF1 |
| | Q96GX2 | ATXN7L3B | P50225 | SULT1A1 | P55268 | LAMB2 | Q9BV29 | CCDC32 |
| | P18887 | XRCC1 | Q9BY42 | RTF2 | P43246 | MSH2 | Q15833 | STXBP2 |
| | P40925 | MDH1 | Q13243 | SRSF5 | Q9UJM3 | ERRFI1 | P14324 | FDPS |
| | O14880 | MGST3 | Q9H0H5 | RACGAP1 | Q99567 | NUP88 | Q9UKX7 | NUP50 |
| | Q8WYP5 | AHCTF1 | P15311 | EZR | P35249 | RFC4 | Q15386 | UBE3C |
| | Q8N129 | CNPY4 | Q16254 | E2F4 | P28749 | RBL1 | O43759 | SYNGR1 |
| | P10909 | CLU | Q9BW19 | KIFC1 | Q9BYK8 | HELZ2 | Q9NWT1 | PAK1IP1 |
| | Q9Y6V7 | DDX49 | P33316 | DUT | P52701 | MSH6 | O95210 | STBD1 |
| | O75821 | EIF3G | P12235 | SLC25A4 | Q9BQE5 | APOL2 | Q9UHQ9 | CYB5R1 |
| | P52732 | KIF11 | Q5VTB9 | RNF220 | P13797 | PLS3 | Q14186 | TFDP1 |

| | | | | | | | | |
|---|---|---|---|---|---|---|---|---|
| | Q9NQ86 | TRIM36 | P11940 | PABPC1 | Q86UP2 | KTN1 | Q13405 | MRPL49 |
| | Q6P1L8 | MRPL14 | Q9NV06 | DCAF13 | Q6KCM7 | SLC25A25 | Q66PJ3 | ARL6IP4 |
| | Q9BZM5 | ULBP2 | Q86WX3 | RPS19BP1 | Q8IWX8 | CHERP | P28161 | GSTM2 |
| | P60228 | EIF3E | O75794 | CDC123 | Q92688 | ANP32B | O95835 | LATS1 |
| | Q9UGY1 | NOL12 | P52815 | MRPL12 | Q9NRZ9 | HELLS | P12004 | PCNA |
| | P41273 | TNFSF9 | P53992 | SEC24C | P24928 | POLR2A | Q9ULJ7 | ANKRD50 |
| | Q96DG6 | CMBL | Q9HCE1 | MOV10 | Q13185 | CBX3 | Q9ULW3 | ABT1 |
| | Q96IG2 | FBXL20 | P38606 | ATP6V1A | Q16762 | TST | P53004 | BLVRA |
| | Q8N488 | RYBP | Q5T5P2 | KIAA1217 | P49247 | RPIA | P11169 | SLC2A3 |
| | Q9H4G0 | EPB41L1 | P31153 | MAT2A | P51151 | RAB9A | Q12849 | GRSF1 |
| | Q5T653 | MRPL2 | Q8WUF5 | PPP1R13L | Q96TA1 | NIBAN2 | Q9P2I0 | CPSF2 |
| | P39880 | CUX1 | O00469 | PLOD2 | Q09161 | NCBP1 | O60831 | PRAF2 |
| | Q96GX5 | MASTL | P21266 | GSTM3 | O75616 | ERAL1 | Q96EQ0 | SGTB |
| | P60842 | EIF4A1 | Q8N1F7 | NUP93 | Q9Y5V3 | MAGED1 | Q8NFC6 | BOD1L1 |
| | Q8N257 | H2BU1 | Q9H7D7 | WDR26 | P78559 | MAP1A | Q9Y6M7 | SLC4A7 |
| | Q6P2Q9 | PRPF8 | O43663 | PRC1 | O00748 | CES2 | Q9NTJ5 | SACM1L |
| | Q86V21 | AACS | Q96PU8 | QKI | Q9HAW4 | CLSPN | P20290 | BTF3 |
| | O75503 | CLN5 | P23786 | CPT2 | P13073 | COX4I1 | O95848 | NUDT14 |
| | Q9BQ13 | KCTD14 | P13674 | P4HA1 | O95861 | BPNT1 | H7C2K6 | EPB41L1 |
| | P53396 | ACLY | P62888 | RPL30 | Q9NVX2 | NLE1 | P52943 | CRIP2 |
| | P19440 | GGT1 | Q9NYV6 | RRN3 | Q9P2T1 | GMPR2 | O75179 | ANKRD17 |
| | O15127 | SCAMP2 | P52565 | ARHGDIA | Q9GZZ1 | NAA50 | Q9HBR0 | SLC38A10 |
| | P18031 | PTPN1 | Q15021 | NCAPD2 | Q9UBT2 | UBA2 | Q8NBF2 | NHLRC2 |
| | O00762 | UBE2C | Q13257 | MAD2L1 | P23193 | TCEA1 | P41134 | ID1 |
| | P42566 | EPS15 | Q7L5L3 | GDPD3 | Q04828 | AKR1C1 | P57075 | UBASH3A |
| | Q96P11 | NSUN5 | O60506 | SYNCRIP | Q9GZU8 | PSME3IP1 | P43155 | CRAT |
| | Q9P253 | VPS18 | Q9NQS7 | INCENP | Q92597 | NDRG1 | Q13217 | DNAJC3 |
| | P78344 | EIF4G2 | Q86U42 | PABPN1 | P04424 | ASL | Q6P2C8 | MED27 |
| | Q8TB40 | ABHD4 | Q9Y6N5 | SQOR | Q8NFZ0 | FBH1 | Q8N5M4 | TTC9C |
| | O94766 | B3GAT3 | Q15366 | PCBP2 | Q9BXI9 | C1QTNF6 | Q9Y6Y0 | IVNS1ABP |
| | Q6YHK3 | CD109 | P28300 | LOX | P33992 | MCM5 | O95169 | NDUFB8 |
| | P11234 | RALB | P26358 | DNMT1 | Q9UHI6 | DDX20 | Q16540 | MRPL23 |
| | Q7Z434 | MAVS | P13051 | UNG | P62263 | RPS14 | O75385 | ULK1 |
| | Q5C9Z4 | NOM1 | Q9BXS6 | NUSAP1 | Q9BTC8 | MTA3 | O96005 | CLPTM1 |
| | P31040 | SDHA | Q15404 | RSU1 | P13667 | PDIA4 | P0CG29 | GSTT2 |
| | P06280 | GLA | Q9Y678 | COPG1 | Q9UI30 | TRMT112 | P33993 | MCM7 |
| | O94906 | PRPF6 | Q5T8P6 | RBM26 | P28715 | ERCC5 | P50552 | VASP |
| | P35555 | FBN1 | Q99633 | PRPF18 | P49642 | PRIM1 | Q13620 | CUL4B |
| | P24385 | CCND1 | Q12874 | SF3A3 | O95486 | SEC24A | Q9UFN0 | NIPSNAP3A |
| | P48444 | ARCN1 | P17405 | SMPD1 | A0A140T997 | HLA-B | Q9H5V9 | STEEP1 |

| | | | | | | | | |
|---|---|---|---|---|---|---|---|---|
| | Q96JC1 | VPS39 | O14578 | CIT | P54105 | CLNS1A | P61165 | TMEM258 |
| | O76021 | RSL1D1 | Q8N5M1 | ATPAF2 | Q9UI26 | IPO11 | Q96EY8 | MMAB |
| | O95208 | EPN2 | O95340 | PAPSS2 | Q03113 | GNA12 | A4D1E9 | GTPBP10 |
| | P29373 | CRABP2 | P02462 | COL4A1 | Q6UB35 | MTHFD1L | P56211 | ARPP19 |
| | Q9P035 | HACD3 | P17931 | LGALS3 | Q9H3H3 | C11orf68 | Q92544 | TM9SF4 |
| | P50148 | GNAQ | Q9BSV6 | TSEN34 | P53634 | CTSC | Q7Z3U7 | MON2 |
| | Q9BW62 | KATNAL1 | Q9BRK3 | MXRA8 | Q13347 | EIF3I | P67809 | YBX1 |
| | O94808 | GFPT2 | Q15436 | SEC23A | O43633 | CHMP2A | P84243 | H3-3A |
| | O95757 | HSPA4L | P49366 | DHPS | O43776 | NARS1 | Q92797 | SYMPK |
| | P18615 | NELFE | Q93050 | ATP6V0A1 | O43166 | SIPA1L1 | P43034 | PAFAH1B1 |
| | Q9Y263 | PLAA | Q9NPA0 | EMC7 | P51530 | DNA2 | Q13619 | CUL4A |
| | A0AV96 | RBM47 | P52292 | KPNA2 | Q96HE7 | ERO1A | Q5JXI8 | FHL1 |
| | O15042 | U2SURP | Q14914 | PTGR1 | P24941 | CDK2 | Q02818 | NUCB1 |
| | O60271 | SPAG9 | P17813 | ENG | P19838 | NFKB1 | P20585 | MSH3 |
| | P99999 | CYCS | P50213 | IDH3A | E9PAV3 | NACA | P32970 | CD70 |
| | Q96HE9 | PRR11 | P42224 | STAT1 | Q9UIG0 | BAZ1B | P41227 | NAA10 |
| | P19387 | POLR2C | P09874 | PARP1 | Q9UKU7 | ACAD8 | Q8IVD9 | NUDCD3 |
| | Q6ZWI9 | RFPL4B | P61011 | SRP54 | Q6PCE3 | PGM2L1 | Q9BYG3 | NIFK |
| | Q9NY33 | DPP3 | Q15645 | TRIP13 | P35658 | NUP214 | Q9Y223 | GNE |
| | Q03135 | CAV1 | P19525 | EIF2AK2 | P61353 | RPL27 | O75718 | CRTAP |
| | Q6NXE6 | ARMC6 | O94979 | SEC31A | Q969Z3 | MTARC2 | P22570 | FDXR |
| | Q96EU6 | RRP36 | P30044 | PRDX5 | Q9Y3R5 | DOP1B | P50914 | RPL14 |
| | Q9Y570 | PPME1 | Q7L1V2 | MON1B | O94992 | HEXIM1 | P13686 | ACP5 |
| | Q9H0U6 | MRPL18 | Q8N556 | AFAP1 | Q9H9P8 | L2HGDH | P53701 | HCCS |
| | Q9Y5Q8 | GTF3C5 | P52823 | STC1 | P46939 | UTRN | Q9BZK7 | TBL1XR1 |
| | Q13586 | STIM1 | Q8NEM2 | SHCBP1 | Q15652 | JMJD1C | Q9Y285 | FARSA |
| | Q9BXI6 | TBC1D10A | Q16851 | UGP2 | Q70Z53 | FRA10AC1 | Q03518 | TAP1 |
| | Q99519 | NEU1 | Q9P2N5 | RBM27 | Q9H6S3 | EPS8L2 | O95817 | BAG3 |
| | Q96PZ0 | PUS7 | Q9P0V9 | SEPTIN10 | Q00610 | CLTC | Q9HC21 | SLC25A19 |
| | Q9H3P7 | ACBD3 | O95833 | CLIC3 | Q9UKF6 | CPSF3 | Q99613 | EIF3C |
| | O75683 | SURF6 | O43251 | RBFOX2 | Q9NZ01 | TECR | Q14008 | CKAP5 |
| | Q15738 | NSDHL | Q9H0U9 | TSPYL1 | Q9UBT7 | CTNNAL1 | P40939 | HADHA |
| | O75396 | SEC22B | Q13895 | BYSL | Q9NVP1 | DDX18 | Q15113 | PCOLCE |
| | X5CMH5 | TAP2 | Q6PK04 | CCDC137 | P36543 | ATP6V1E1 | O43813 | LANCL1 |
| | Q96S97 | MYADM | P09110 | ACAA1 | P08238 | HSP90AB1 | P62241 | RPS8 |
| | P49189 | ALDH9A1 | O60763 | USO1 | O95801 | TTC4 | Q8N5C7 | DTWD1 |
| | Q16576 | RBBP7 | Q96EK4 | THAP11 | Q9NX24 | NHP2 | P55769 | SNU13 |
| | P16070 | CD44 | P49790 | NUP153 | Q9BV44 | THUMPD3 | Q13148 | TARDBP |
| | Q9NUL7 | DDX28 | O75153 | CLUH | P05388 | RPLP0 | Q96EK7 | FAM120B |
| | O14776 | TCERG1 | Q6AI08 | HEATR6 | P28065 | PSMB9 | Q96KC8 | DNAJC1 |

| | | | | | | | | |
|---|---|---|---|---|---|---|---|---|
| | P41240 | CSK | Q08AF3 | SLFN5 | Q9UNF0 | PACSIN2 | P55265 | ADAR |
| | Q6UVJ0 | SASS6 | Q8WWY3 | PRPF31 | P78316 | NOP14 | Q15750 | TAB1 |
| | Q5JTH9 | RRP12 | Q9NQX7 | ITM2C | Q15052 | ARHGEF6 | Q86V88 | MDP1 |
| | O96028 | NSD2 | Q9NZE8 | MRPL35 | Q5JTZ9 | AARS2 | Q4L235 | AASDH |
| | O00429 | DNM1L | Q9NQW6 | ANLN | Q15014 | MORF4L2 | P23368 | ME2 |
| | O14979 | HNRNPDL | Q8N3U4 | STAG2 | P39019 | RPS19 | Q01105 | SET |
| | P78524 | DENND2B | O75175 | CNOT3 | P51668 | UBE2D1 | P17987 | TCP1 |
| | Q6P1K8 | GTF2H2C | P82675 | MRPS5 | Q9BUJ2 | HNRNPUL1 | Q8ND56 | LSM14A |
| | P31146 | CORO1A | Q9Y2K7 | KDM2A | Q5VTR2 | RNF20 | Q96T51 | RUFY1 |
| | Q9UH62 | ARMCX3 | Q9UKJ3 | GPATCH8 | O00303 | EIF3F | P51784 | USP11 |
| | Q99717 | SMAD5 | Q9H0Q0 | CYRIA | Q12802 | AKAP13 | P04439 | HLA-A |
| | O75173 | ADAMTS4 | P62829 | RPL23 | P62906 | RPL10A | O60701 | UGDH |
| | P56545 | CTBP2 | P46776 | RPL27A | P11498 | PC | Q7L014 | DDX46 |
| | Q13907 | IDI1 | P13639 | EEF2 | Q8TDW7 | FAT3 | Q96PD2 | DCBLD2 |
| | P46778 | RPL21 | P52788 | SMS | P07339 | CTSD | Q15058 | KIF14 |
| | Q14692 | BMS1 | P16083 | NQO2 | P08572 | COL4A2 | Q15365 | PCBP1 |
| | Q15233 | NONO | P08397 | HMBS | P45954 | ACADSB | Q16719 | KYNU |
| | A0A494C072 | CHN1 | P17844 | DDX5 | A0A0A6YYJ5 | MACF1 | O75369 | FLNB |
| | Q15392 | DHCR24 | O15031 | PLXNB2 | P06730 | EIF4E | Q16610 | ECM1 |
| | Q9NW13 | RBM28 | P42285 | MTREX | O95394 | PGM3 | Q5K651 | SAMD9 |
| | P61254 | RPL26 | P30084 | ECHS1 | Q9Y223 | GNE | Q14566 | MCM6 |
| | Q9Y2Q3 | GSTK1 | P21283 | ATP6V1C1 | Q86XP3 | DDX42 | Q9H7E9 | C8orf33 |
| | Q13232 | NME3 | Q969R2 | OSBP2 | Q8TDQ7 | GNPDA2 | Q9UHX1 | PUF60 |
| | P04179 | SOD2 | P54725 | RAD23A | Q9Y4D1 | DAAM1 | Q13509 | TUBB3 |
| | Q9UN86 | G3BP2 | P46379 | BAG6 | Q8NEZ2 | VPS37A | Q02543 | RPL18A |
| | Q8TDD1 | DDX54 | Q9Y3B7 | MRPL11 | Q8NFH4 | NUP37 | P83111 | LACTB |
| | O94985 | CLSTN1 | Q9UNQ2 | DIMT1 | Q7KZ85 | SUPT6H | P56377 | AP1S2 |
| | Q96CW5 | TUBGCP3 | A0A7P0Z439 | CUL4B | Q86Y56 | DNAAF5 | Q16611 | BAK1 |
| | P51452 | DUSP3 | O95239 | KIF4A | Q92522 | H1-10 | Q9BRJ6 | C7orf50 |
| | P20340 | RAB6A | P11217 | PYGM | Q15274 | QPRT | Q8N3V7 | SYNPO |
| | P35237 | SERPINB6 | Q56NI9 | ESCO2 | Q9Y4B6 | DCAF1 | P06132 | UROD |
| | Q53EL6 | PDCD4 | Q06210 | GFPT1 | O95810 | CAVIN2 | Q8TCU4 | ALMS1 |
| | Q71RG4 | TMUB2 | Q96N66 | MBOAT7 | P04183 | TK1 | Q9ULI3 | HEG1 |
| | P53365 | ARFIP2 | Q9ULC5 | ACSL5 | Q9NYZ3 | GTSE1 | Q9Y6A5 | TACC3 |
| | O15318 | POLR3G | Q9BY50 | SEC11C | P30041 | PRDX6 | O60307 | MAST3 |
| | P35754 | GLRX | Q16134 | ETFDH | Q99615 | DNAJC7 | Q8TEX9 | IPO4 |
| | O94760 | DDAH1 | Q86VN1 | VPS36 | O75330 | HMMR | P00750 | PLAT |
| | P42677 | RPS27 | P62140 | PPP1CB | Q8NFW8 | CMAS | P62316 | SNRPD2 |
| | Q13435 | SF3B2 | Q9BTE3 | MCMBP | Q6NUM9 | RETSAT | O94967 | WDR47 |
| | P0C0S8 | H2AC11 | Q12965 | MYO1E | P36952 | SERPINB5 | Q13428 | TCOF1 |

| | | | | | | | | |
|---|---|---|---|---|---|---|---|---|
| | P40261 | NNMT | P46779 | RPL28 | P06756 | ITGAV | O00487 | PSMD14 |
| | Q9P031 | CCDC59 | P50991 | CCT4 | P13928 | ANXA8 | O43504 | LAMTOR5 |
| | Q9UH65 | SWAP70 | P35790 | CHKA | P46821 | MAP1B | O94925 | GLS |
| | O76031 | CLPX | Q9Y487 | ATP6V0A2 | P11182 | DBT | Q9UNF1 | MAGED2 |
| | Q03519 | TAP2 | Q12933 | TRAF2 | Q9BZI7 | UPF3B | Q9NVA2 | SEPTIN11 |
| | Q969X5 | ERGIC1 | Q8TF42 | UBASH3B | Q9UHC9 | NPC1L1 | Q86TS9 | MRPL52 |
| | P36578 | RPL4 | Q9BYD1 | MRPL13 | Q9UI42 | CPA4 | Q9UHY1 | NRBP1 |
| | Q9NRX2 | MRPL17 | O60814 | H2BC12 | P38919 | EIF4A3 | Q9H8V3 | ECT2 |
| | P32455 | GBP1 | Q96ET8 | TVP23B | Q96T88 | UHRF1 | Q13671 | RIN1 |
| | Q8IV08 | PLD3 | Q15050 | RRS1 | O14965 | AURKA | Q8N9N8 | EIF1AD |
| | Q6UW78 | UQCC3 | Q9NWT8 | AURKAIP1 | Q8WWI1 | LMO7 | P49792 | RANBP2 |
| | P51159 | RAB27A | Q8N983 | MRPL43 | E9PMS6 | LMO7 | Q13618 | CUL3 |
| | O95400 | CD2BP2 | P61619 | SEC61A1 | P37268 | FDFT1 | Q96PU4 | UHRF2 |
| | Q8TF05 | PPP4R1 | Q9Y6X9 | MORC2 | Q6NZI2 | CAVIN1 | Q9UMX1 | SUFU |
| | Q96E29 | MTERF3 | P07384 | CAPN1 | O95259 | KCNH1 | P46013 | MKI67 |
| | Q05086 | UBE3A | Q9BVG4 | PBDC1 | Q2VPK5 | CTU2 | P10915 | HAPLN1 |
| | Q9UPN7 | PPP6R1 | O14684 | PTGES | Q9H0D6 | XRN2 | Q49AR2 | C5orf22 |
| | Q92930 | RAB8B | Q08380 | LGALS3BP | P12270 | TPR | Q8IVT2 | MISP |
| | Q9H0X9 | OSBPL5 | Q14139 | UBE4A | Q9H967 | WDR76 | Q96T37 | RBM15 |
| | Q9NPD8 | UBE2T | Q9NYK5 | MRPL39 | Q16881 | TXNRD1 | P51398 | DAP3 |
| | Q9NW82 | WDR70 | Q587I9 | SFT2D3 | P83876 | TXNL4A | O75521 | ECI2 |
| | P07108 | DBI | P00374 | DHFR | P13716 | ALAD | Q03169 | TNFAIP2 |
| | Q8IXM3 | MRPL41 | Q8IY17 | PNPLA6 | O43390 | HNRNPR | Q9NZM5 | NOP53 |
| | Q15005 | SPCS2 | O75815 | BCAR3 | P05161 | ISG15 | Q04323 | UBXN1 |
| | Q9UNS1 | TIMELESS | Q8NCN5 | PDPR | Q8TCJ2 | STT3B | Q8N6H7 | ARFGAP2 |
| | Q8TAE8 | GADD45GIP1 | P83731 | RPL24 | Q8TAQ2 | SMARCC2 | Q9BYD6 | MRPL1 |
| | P07738 | BPGM | P98170 | XIAP | Q460N5 | PARP14 | P62805 | H4C1 |
| | Q8NC60 | NOA1 | O43149 | ZZEF1 | F8VRH0 | PCBP2 | Q96B97 | SH3KBP1 |
| | Q9NXK8 | FBXL12 | O43314 | PPIP5K2 | Q9BW85 | YJU2 | P27635 | RPL10 |
| | Q8N7H5 | PAF1 | O43290 | SART1 | P62899 | RPL31 | Q9Y6K5 | OAS3 |
| | Q12834 | CDC20 | P17655 | CAPN2 | P35914 | HMGCL | Q7Z3C6 | ATG9A |
| | Q9H6E5 | TUT1 | Q86U38 | NOP9 | Q96MF7 | NSMCE2 | O75691 | UTP20 |
| | P35606 | COPB2 | P26373 | RPL13 | P53007 | SLC25A1 | Q00653 | NFKB2 |
| | Q969E8 | TSR2 | Q15029 | EFTUD2 | Q92841 | DDX17 | O15460 | P4HA2 |
| | Q6P179 | ERAP2 | Q15056 | EIF4H | Q8WWH5 | TRUB1 | P47914 | RPL29 |
| | O43172 | PRPF4 | Q02750 | MAP2K1 | Q14554 | PDIA5 | P63096 | GNAI1 |
| | O75348 | ATP6V1G1 | Q9Y4W6 | AFG3L2 | O76074 | PDE5A | P27694 | RPA1 |
| | Q16850 | CYP51A1 | Q14999 | CUL7 | Q8TCT7 | SPPL2B | Q6ZN18 | AEBP2 |
| | Q9Y3U8 | RPL36 | P14649 | MYL6B | Q8TCB0 | IFI44 | Q8N163 | CCAR2 |
| | Q9Y2W6 | TDRKH | Q86WV6 | STING1 | Q01844 | EWSR1 | Q5SRE5 | NUP188 |

| | | | | | | | | |
|---|---|---|---|---|---|---|---|---|
| | O94901 | SUN1 | O75223 | GGCT | Q8IWC1 | MAP7D3 | Q9UNE7 | STUB1 |
| | Q9NRG9 | AAAS | Q9P0J1 | PDP1 | P52306 | RAP1GDS1 | O75534 | CSDE1 |
| | Q8N567 | ZCCHC9 | Q15637 | SF1 | P22033 | MMUT | Q9NSI2 | SLX9 |
| | P21281 | ATP6V1B2 | Q05932 | FPGS | Q10471 | GALNT2 | Q14498 | RBM39 |
| | Q9BVS4 | RIOK2 | P29590 | PML | O00488 | ZNF593 | Q9NYU2 | UGGT1 |
| | Q12769 | NUP160 | P12268 | IMPDH2 | Q02338 | BDH1 | Q02218 | OGDH |
| | Q9BVI4 | NOC4L | Q9NTM9 | CUTC | P41226 | UBA7 | Q9UQE7 | SMC3 |
| | Q9H6F5 | CCDC86 | A6NCE7 | MAP1LC3B | Q8TF72 | SHROOM3 | Q15397 | PUM3 |
| | P04062 | GBA | Q9Y4X5 | ARIH1 | P62913 | RPL11 | P05362 | ICAM1 |
| | P23588 | EIF4B | Q9NP87 | POLM | P50151 | GNG10 | Q6P996 | PDXDC1 |
| | P28288 | ABCD3 | Q9UHG3 | PCYOX1 | Q9BSA9 | TMEM175 | Q9NYY8 | FASTKD2 |
| | Q9BTX1 | NDC1 | Q9Y4P1 | ATG4B | Q9H6S0 | YTHDC2 | P13693 | TPT1 |
| | Q86XI2 | NCAPG2 | Q6P1Q0 | LETMD1 | Q9NRN7 | AASDHPPT | Q08431 | MFGE8 |
| | P27797 | CALR | A0A494C0R8 | CLUH | Q9NR56 | MBNL1 | Q13268 | DHRS2 |
| | P26022 | PTX3 | P55789 | GFER | Q6PIU2 | NCEH1 | Q92925 | SMARCD2 |
| | O43854 | EDIL3 | P27701 | CD82 | P63244 | RACK1 | B7Z4M1 | RTN3 |
| | Q9BSJ2 | TUBGCP2 | P16435 | POR | Q5T4S7 | UBR4 | Q5TC84 | OGFRL1 |
| | P49184 | DNASE1L1 | Q92879 | CELF1 | Q9BQC3 | DPH2 | Q9UIC8 | LCMT1 |
| | Q14C86 | GAPVD1 | Q03252 | LMNB2 | O43731 | KDELR3 | P08253 | MMP2 |
| | Q9UHD8 | SEPTIN9 | P40123 | CAP2 | P48449 | LSS | O75695 | RP2 |
| | Q96AH0 | NABP1 | P18124 | RPL7 | Q8NBI2 | CYB561A3 | Q8WU10 | PYROXD1 |
| | Q96GQ7 | DDX27 | Q13283 | G3BP1 | Q13228 | SELENBP1 | Q9GZR7 | DDX24 |
| | P09001 | MRPL3 | Q96EY5 | MVB12A | Q9BQ52 | ELAC2 | Q53EZ4 | CEP55 |
| | Q9BVP2 | GNL3 | Q9Y399 | MRPS2 | Q14493 | SLBP | Q9UQN3 | CHMP2B |
| | O75400 | PRPF40A | Q9NQT4 | EXOSC5 | Q9BQG0 | MYBBP1A | Q9BZ67 | FRMD8 |
| | Q8TDZ2 | MICAL1 | Q8IU85 | CAMK1D | Q7L1Q6 | BZW1 | O60216 | RAD21 |
| | Q14147 | DHX34 | Q2TAY7 | SMU1 | O95721 | SNAP29 | P49257 | LMAN1 |
| | Q86SQ0 | PHLDB2 | P28702 | RXRB | P30876 | POLR2B | Q7L8W6 | DPH6 |
| | Q86SK9 | SCD5 | P46060 | RANGAP1 | P55081 | MFAP1 | O60921 | HUS1 |
| | Q68CQ4 | UTP25 | Q15746 | MYLK | P49005 | POLD2 | O14929 | HAT1 |
| | Q9Y5V0 | ZNF706 | Q8TF66 | LRRC15 | O95772 | STARD3NL | P29466 | CASP1 |
| | P50995 | ANXA11 | Q96CS2 | HAUS1 | Q8TCT9 | HM13 | Q9H8W3 | FAM204A |
| | Q9BXK5 | BCL2L13 | Q9Y2R9 | MRPS7 | P61221 | ABCE1 | Q15714 | TSC22D1 |
| | P35250 | RFC2 | P62072 | TIMM10 | E9PMP7 | LMO7 | Q7Z3T8 | ZFYVE16 |
| | Q13393 | PLD1 | Q13685 | AAMP | Q9UKD1 | GMEB2 | P15085 | CPA1 |
| | Q5T8D3 | ACBD5 | Q2TB90 | HKDC1 | Q5VT79 | ANXA8L1 | P04843 | RPN1 |
| | O75382 | TRIM3 | Q9BUQ8 | DDX23 | Q15554 | TERF2 | P14678 | SNRPB |
| | O15371 | EIF3D | Q96JJ3 | ELMO2 | Q96DV4 | MRPL38 | P61964 | WDR5 |
| | O76024 | WFS1 | Q8NFQ8 | TOR1AIP2 | P30043 | BLVRB | P61086 | UBE2K |
| | P51571 | SSR4 | Q8IYS1 | PM20D2 | O00159 | MYO1C | O43301 | HSPA12A |

| | | | | | | | | |
|---|---|---|---|---|---|---|---|---|
| | P17096 | HMGA1 | Q6P5R6 | RPL22L1 | Q16270 | IGFBP7 | P18440 | NAT1 |
| | Q9NVS2 | MRPS18A | P40429 | RPL13A | O95905 | ECD | Q8NE86 | MCU |
| | Q9BXY0 | MAK16 | Q15771 | RAB30 | Q6PD62 | CTR9 | P00813 | ADA |
| | P49419 | ALDH7A1 | O96011 | PEX11B | Q13356 | PPIL2 | Q5JSH3 | WDR44 |
| | P31943 | HNRNPH1 | Q9NP77 | SSU72 | Q6NZY4 | ZCCHC8 | Q8IYS2 | KIAA2013 |
| | Q2M2I8 | AAK1 | Q9NPQ8 | RIC8A | Q9NZN8 | CNOT2 | Q13614 | MTMR2 |
| | Q5T3I0 | GPATCH4 | P34741 | SDC2 | O95900 | TRUB2 | P09417 | QDPR |
| | Q01082 | SPTBN1 | Q8NBM8 | PCYOX1L | P19634 | SLC9A1 | Q9BXP5 | SRRT |
| | O15357 | INPPL1 | Q96E39 | RBMXL1 | P24534 | EEF1B2 | P11802 | CDK4 |
| | P62854 | RPS26 | Q9Y3E5 | PTRH2 | Q6ZW76 | ANKS3 | P78346 | RPP30 |
| | Q96GD4 | AURKB | Q8IWZ8 | SUGP1 | Q8IY67 | RAVER1 | O75376 | NCOR1 |
| | P46063 | RECQL | Q9Y5N6 | ORC6 | Q96P16 | RPRD1A | O43427 | FIBP |
| | Q9H0S4 | DDX47 | P46782 | RPS5 | Q9Y6I4 | USP3 | P14921 | ETS1 |
| | O43772 | SLC25A20 | Q99551 | MTERF1 | Q14980 | NUMA1 | Q9BZQ6 | EDEM3 |
| | Q96I51 | RCC1L | O00193 | SMAP | Q8NC54 | KCT2 | Q9NSG2 | C1orf112 |
| | P01889 | HLA-B | O60343 | TBC1D4 | P19793 | RXRA | Q7L590 | MCM10 |
| | P53621 | COPA | O60784 | TOM1 | Q8IWA4 | MFN1 | P62249 | RPS16 |
| | P30046 | DDT | Q6PID8 | KLHDC10 | Q96HH9 | GRAMD2B | Q9Y618 | NCOR2 |
| | Q99988 | GDF15 | Q9HAF1 | MEAF6 | P86790 | CCZ1 | O43805 | SSNA1 |
| | Q9Y2R0 | COA3 | Q13418 | ILK | O14879 | IFIT3 | Q9UM54 | MYO6 |
| | Q14116 | IL18 | Q9BT40 | INPP5K | P00966 | ASS1 | O75970 | MPDZ |
| | P08133 | ANXA6 | O60832 | DKC1 | P07947 | YES1 | Q9Y5S1 | TRPV2 |
| | Q86U90 | YRDC | Q9BQP7 | MGME1 | A5PLL7 | PEDS1 | Q9BZE1 | MRPL37 |
| | Q7Z3B4 | NUP54 | Q6PKG0 | LARP1 | J3KQL8 | APOL2 | Q92900 | UPF1 |
| | Q9Y4C8 | RBM19 | Q7Z2T5 | TRMT1L | Q6Y7W6 | GIGYF2 | Q14012 | CAMK1 |
| | O00194 | RAB27B | Q86V48 | LUZP1 | Q13112 | CHAF1B | P50452 | SERPINB8 |
| | Q99584 | S100A13 | Q13371 | PDCL | O14735 | CDIPT | P50454 | SERPINH1 |
| | Q86WJ1 | CHD1L | Q9UH17 | APOBEC3B | O00161 | SNAP23 | Q9BUF5 | TUBB6 |
| | P53990 | IST1 | Q8NC44 | RETREG2 | Q8NBT2 | SPC24 | Q8N2G8 | GHDC |
| | O15254 | ACOX3 | Q9Y5P4 | CERT1 | P20700 | LMNB1 | Q96Q15 | SMG1 |
| | O75947 | ATP5PD | Q99536 | VAT1 | Q14644 | RASA3 | O00267 | SUPT5H |
| | P01033 | TIMP1 | Q9NTJ3 | SMC4 | Q86X02 | CDR2L | Q8NI60 | COQ8A |
| | P00367 | GLUD1 | P07093 | SERPINE2 | Q8TD55 | PLEKHO2 | P42226 | STAT6 |
| | Q14011 | CIRBP | Q96FK6 | WDR89 | O43395 | PRPF3 | Q9P2R3 | ANKFY1 |
| | Q9Y4Z0 | LSM4 | P68104 | EEF1A1 | Q5JPH6 | EARS2 | P49454 | CENPF |
| | P16298 | PPP3CB | P12111 | COL6A3 | Q92851 | CASP10 | Q9BTY7 | HGH1 |
| | Q96SI9 | STRBP | O94804 | STK10 | Q07065 | CKAP4 | Q5TAQ9 | DCAF8 |
| | Q8N4V1 | MMGT1 | Q9NWZ5 | UCKL1 | Q9Y4C2 | TCAF1 | Q8TDJ6 | DMXL2 |
| | Q96IU4 | ABHD14B | Q9Y4C1 | KDM3A | Q86V81 | ALYREF | Q15545 | TAF7 |
| | Q9GZY8 | MFF | Q9UBQ5 | EIF3K | Q9BT22 | ALG1 | Q8IXH7 | NELFCD |

| | | | | | | | | |
|---|---|---|---|---|---|---|---|---|
| | P78357 | CNTNAP1 | P30519 | HMOX2 | P39023 | RPL3 | Q3ZCW2 | LGALSL |
| | Q96DZ1 | ERLEC1 | Q9NRK6 | ABCB10 | Q9Y314 | NOSIP | Q8N3Z3 | GTPBP8 |
| | Q9NR19 | ACSS2 | P01040 | CSTA | P36954 | POLR2I | Q7L592 | NDUFAF7 |
| | Q9BTL3 | RAMAC | P35052 | GPC1 | P62841 | RPS15 | Q9Y606 | PUS1 |
| | Q7Z7F7 | MRPL55 | P51003 | PAPOLA | P56937 | HSD17B7 | O94782 | USP1 |
| | Q6GMV2 | SMYD5 | Q9NSP4 | CENPM | Q9Y3X0 | CCDC9 | Q86VP1 | TAX1BP1 |
| | Q13813 | SPTAN1 | P62424 | RPL7A | P07311 | ACYP1 | P35269 | GTF2F1 |
| | Q96BW5 | PTER | Q8N5S9 | CAMKK1 | O43187 | IRAK2 | P62942 | FKBP1A |
| | Q16514 | TAF12 | A0A8C8KBL6 | ITGA6 | Q8WVF1 | OSCP1 | Q92973 | TNPO1 |
| | Q9NUE0 | ZDHHC18 | Q8WWK9 | CKAP2 | Q9NZU5 | LMCD1 | Q02252 | ALDH6A1 |
| | P49207 | RPL34 | Q9Y5J9 | TIMM8B | Q13488 | TCIRG1 | Q96M27 | PRRC1 |
| | Q9BZD2 | SLC29A3 | Q9UPY8 | MAPRE3 | Q9Y296 | TRAPPC4 | Q6SZW1 | SARM1 |
| | Q9NUW8 | TDP1 | A0A140T913 | HLA-A | P30626 | SRI | Q9H8M7 | MINDY3 |
| | P52597 | HNRNPF | Q9H477 | RBKS | P98082 | DAB2 | Q9ULG6 | CCPG1 |
| | O43747 | AP1G1 | Q8WWV3 | RTN4IP1 | Q13308 | PTK7 | Q96GA3 | LTV1 |
| | P51659 | HSD17B4 | Q9UPN3 | MACF1 | Q15629 | TRAM1 | P34059 | GALNS |
| | P62879 | GNB2 | Q12979 | ABR | P55084 | HADHB | P60510 | PPP4C |
| | Q8NBZ7 | UXS1 | Q07960 | ARHGAP1 | P11388 | TOP2A | Q8N128 | FAM177A1 |
| | Q96CM8 | ACSF2 | Q8WUP2 | FBLIM1 | Q9UBS4 | DNAJB11 | P05114 | HMGN1 |
| | Q9BSC4 | NOL10 | Q7Z4W1 | DCXR | P23634 | ATP2B4 | Q9Y2Z0 | SUGT1 |
| | P08842 | STS | Q96CV9 | OPTN | Q9UIW2 | PLXNA1 | A5D8V6 | VPS37C |
| | Q96B96 | LDAF1 | Q8NC26 | ZNF114 | Q9Y2D5 | AKAP2 | O43731 | KDELR3 |
| | Q6P4F2 | FDX2 | P32189 | GK | Q14674 | ESPL1 | Q8IVL5 | P3H2 |
| | Q14683 | SMC1A | O75915 | ARL6IP5 | P11166 | SLC2A1 | P09525 | ANXA4 |
| | Q495W5 | FUT11 | P23258 | TUBG1 | Q92917 | GPKOW | Q92830 | KAT2A |
| | P49406 | MRPL19 | Q9HAN9 | NMNAT1 | Q9NX20 | MRPL16 | Q9UBU8 | MORF4L1 |
| | Q92614 | MYO18A | P46459 | NSF | P29218 | IMPA1 | Q8IYQ7 | THNSL1 |
| | P00973 | OAS1 | Q8TCC3 | MRPL30 | P55145 | MANF | Q16342 | PDCD2 |
| | P60903 | S100A10 | P16260 | SLC25A16 | P16278 | GLB1 | P49023 | PXN |
| | Q9Y383 | LUC7L2 | Q7LGA3 | HS2ST1 | Q8WZ82 | OVCA2 | Q9UEE5 | STK17A |
| | Q9Y6C2 | EMILIN1 | Q16629 | SRSF7 | P10301 | RRAS | Q6P1J9 | CDC73 |
| | Q969M3 | YIPF5 | Q6P1A2 | LPCAT3 | Q9NTZ6 | RBM12 | Q6RFH5 | WDR74 |
| | Q9BPW9 | DHRS9 | Q9UBF2 | COPG2 | P61803 | DAD1 | O00330 | PDHX |
| | Q9UP95 | SLC12A4 | Q9Y484 | WDR45 | A0A1B0GTL5 | RAB11FIP5 | P00390 | GSR |
| | Q9NP81 | SARS2 | Q9H330 | TMEM245 | Q6PI78 | TMEM65 | Q13423 | NNT |
| | Q16790 | CA9 | Q8IYB7 | DIS3L2 | P41970 | ELK3 | P52657 | GTF2A2 |
| | Q66K74 | MAP1S | Q96AE4 | FUBP1 | Q6P1Q9 | METTL2B | Q969P0 | IGSF8 |
| | Q8N2K0 | ABHD12 | Q9BXF6 | RAB11FIP5 | P50613 | CDK7 | Q96N67 | DOCK7 |
| | Q08722 | CD47 | A0A0G2JJL1 | WDR46 | Q6DKI1 | RPL7L1 | O15431 | SLC31A1 |
| | Q9NP74 | PALMD | Q8NFH5 | NUP35 | Q9BZL1 | UBL5 | Q86TI2 | DPP9 |

| | | | | | | | | |
|---|---|---|---|---|---|---|---|---|
| | P62312 | LSM6 | Q9NUV7 | SPTLC3 | Q8ND24 | RNF214 | Q5VV42 | CDKAL1 |
| | Q9NRH1 | YAE1 | Q13330 | MTA1 | P13984 | GTF2F2 | | |
| | Q92871 | PMM1 | Q5T200 | ZC3H13 | P48163 | ME1 | | |
| | O75643 | SNRNP200 | Q9BW71 | HIRIP3 | Q9H9C1 | VIPAS39 | | |
| **SW1353** | O00499 | BIN1 | Q9BW19 | KIFC1 | P14635 | CCNB1 | P04183 | TK1 |
| | Q8NF37 | LPCAT1 | Q9Y223 | GNE | P31350 | RRM2 | P05121 | SERPINE1 |
| | Q27J81 | INF2 | Q13309 | SKP2 | Q96GD4 | AURKB | P53814 | SMTN |
| | Q8NFW8 | CMAS | Q8N2K1 | UBE2J2 | Q9NQW6 | ANLN | Q9Y263 | PLAA |
| | Q14103 | HNRNPD | P78347 | GTF2I | P22570 | FDXR | Q8WWK9 | CKAP2 |
| | Q96HE9 | PRR11 | Q9H6S3 | EPS8L2 | Q92466 | DDB2 | P27694 | RPA1 |
| | Q8NEM2 | SHCBP1 | A0A1B0GTL5 | RAB11FIP5 | O95757 | HSPA4L | P52732 | KIF11 |
| | O14786 | NRP1 | P18858 | LIG1 | Q99615 | DNAJC7 | P06493 | CDK1 |
| | P30530 | AXL | Q9Y6A5 | TACC3 | Q9UBB4 | ATXN10 | Q6Y1H2 | HACD2 |

**Suppl table 2: Enrichment analysis of proteins whose expression is significantly deregulated by cladribine in SW1353 chondrosarcoma cells.** Analysis considering the GO Biological Process (BP) term and KEGGs using Shinygo.

| Data base | FDR | nGenes | Pathway Genes | Fold Enrichment | Terms | Genes |
|---|---|---|---|---|---|---|
| GO BP | 0.0019 | 2 | 4 | 346.68 | GO:1905448 positive reg. of mitochondrial ATP synthesis coupled electron transport | CCNB1 CDK1 |
| | 0.0019 | 5 | 204 | 16.99 | GO:0071897 DNA biosynthetic proc. | RPA1 LIG1 HNRNPD TK1 AURKB |
| | 0.0012 | 7 | 476 | 10.20 | GO:0044772 mitotic cell cycle phase transition | CCNB1 SKP2 CDK1 AURKB RRM2 TACC3 ANLN |
| | 0.0003 | 10 | 825 | 8.40 | GO:1903047 mitotic cell cycle proc. | ANLN LIG1 CCNB1 KIF11 SKP2 CDK1 AURKB RRM2 TACC3 CKAP2 |
| | 0.0012 | 8 | 689 | 8.05 | GO:0051301 cell division | ANLN AURKB TACC3 LIG1 CCNB1 KIF11 CDK1 CKAP2 |
| | 0.0019 | 8 | 781 | 7.10 | GO:0010564 reg. of cell cycle proc. | CDK1 AURKB CCNB1 BIN1 RRM2 TACC3 KIF11 ANLN |
| | 0.0006 | 10 | 996 | 6.96 | GO:0000278 mitotic cell cycle | ANLN LIG1 CCNB1 KIF11 SKP2 CDK1 AURKB RRM2 TACC3 CKAP2 |
| | 0.0006 | 11 | 1321 | 5.77 | GO:0022402 cell cycle proc. | ANLN LIG1 CCNB1 KIF11 SKP2 CDK1 AURKB BIN1 RRM2 TACC3 CKAP2 |
| | 0.0012 | 10 | 1213 | 5.72 | GO:0051726 reg. of cell cycle | CCNB1 SKP2 CDK1 AURKB PRR11 BIN1 RRM2 TACC3 KIF11 ANLN |
| | 0.0028 | 9 | 1138 | 5.48 | GO:0006259 DNA metabolic proc. | LIG1 RPA1 DDB2 GTF2I HNRNPD TK1 CDK1 RRM2 AURKB |
| | 0.0018 | 11 | 1594 | 4.78 | GO:0007010 cytoskeleton organization | ANLN TACC3 CKAP2 KIF11 AURKB SMTN BIN1 INF2 NRP1 CCNB1 CDK1 |

| | | | | | | |
|---|---|---|---|---|---|---|
| | 0.0006 | 13 | 1937 | 4.65 | GO:0007049 cell cycle | ANLN LIG1 RPA1 CCNB1 KIF11 SKP2 CDK1 AURKB PRR11 BIN1 RRM2 TACC3 CKAP2 |
| | 0.0028 | 13 | 2509 | 3.59 | GO:0051128 reg. of cellular component organization | CKAP2 EPS8L2 AURKB NRP1 SERPINE1 BIN1 PLAA HNRNPD ANLN TACC3 CCNB1 AXL INF2 |
| **KEGG** | 0.0103 | 2 | 23 | 60.29 | Path:hsa03430 Mismatch repair | LIG1 RPA1 |
| | 0.0012 | 3 | 46 | 45.22 | Path:hsa03420 Nucleotide excision repair | DDB2 LIG1 RPA1 |
| | 0.0127 | 2 | 36 | 38.52 | Path:hsa03030 DNA replication | LIG1 RPA1 |
| | 0.0002 | 4 | 73 | 37.99 | Path:hsa04115 p53 signaling pathway | DDB2 SERPINE1 RRM2 CCNB1 |
| | 0.0157 | 2 | 49 | 28.30 | Path:hsa00520 Amino sugar and nucleotide sugar metabolism | GNE CMAS |
| | 0.0197 | 2 | 58 | 23.91 | Path:hsa00240 Pyrimidine metabolism | RRM2 TK1 |
| | 0.0327 | 2 | 79 | 17.55 | Path:hsa00983 Drug metabolism-other enzymes | RRM2 TK1 |
| | 0.0122 | 3 | 126 | 16.51 | Path:hsa04110 Cell cycle | SKP2 CCNB1 CDK1 |
| | 0.0345 | 2 | 85 | 16.31 | Path:hsa01232 Nucleotide metabolism | RRM2 TK1 |
| | 0.0371 | 2 | 92 | 15.07 | Path:hsa05222 Small cell lung cancer | DDB2 SKP2 |
| | 0.0127 | 3 | 142 | 14.65 | Path:hsa04120 Ubiquitin mediated proteolysis | UBE2J2 DDB2 SKP2 |
| | 0.0421 | 2 | 102 | 13.60 | Path:hsa04914 Progesterone-mediated oocyte maturation | CCNB1 CDK1 |
| | 0.0129 | 3 | 156 | 13.33 | Path:hsa04218 Cellular senescence | SERPINE1 CCNB1 CDK1 |
| | 0.0142 | 3 | 169 | 12.31 | Path:hsa04141 Protein processing in endoplasmic reticulum | UBE2J2 HSPA4L PLAA |

**Suppl table 3: Enrichment analysis of proteins whose expression is significantly deregulated by cladribine in JJ012 chondrosarcoma cells.** Analysis considering the GO Biological Process (BP) term. and KEGGs using Shinygo

| Data base | FDR | nGenes | Pathway Genes | Fold Enrichment | Terms | Genes |
|---|---|---|---|---|---|---|
| GO BP | 1.35E-21 | 51 | 178 | 5.54 | GO:0002181 cytoplasmic translation | EIF4B RPL31 EIF3I RPL6 RPLP0 EIF4H RPL24 RPL36 DPH2 DPH6 EIF4A1 RPL26 RPL29 RPL22L1 RPS5 RPL3 EIF3D EIF3E RPS16 RPS19 RPL18A RPL28 RPL34 RPS15 RPL21 RPL5 RPL23 EIF3G RPL27 NCBP1 RPL13A RPL11 RPS8 RPL7 RPL30 RPS14 RPL27A RPL13 RPL4 EIF3F RPS27 EIF3C RPL14 RPS26 RPL10A RACK1 CSDE1 YBX1 PABPC1 SYNCRIP SLBP |
| | 1.86E-21 | 59 | 240 | 4.75 | GO:0022618 ribonucleoprotein complex assembly | SFSWAP EIF4B DDX20 NLE1 RPS5 RPL6 RPLP0 RPL3 PRPF6 CRNKL1 RPS19 NOP53 EIF4H RPS15 RPL5 MRPS7 SNRPD2 LSM4 ERAL1 CDC73 SNRNP200 ABT1 LUC7L2 CELF1 RPS14 PRPF18 RAMAC RPS27 |

| | | | | | | |
|---|---|---|---|---|---|---|
| | | | | | | DDX28 CLNS1A EIF3I SF3B2 SF3A1 EIF3D EIF3E SNRPB EIF3G DDX46 DDX23 EIF3F SF3A3 EIF3C HSP90AB1 PRPF3 RPL13A ADAR SNU13 YJU2 RPL24 FASTKD2 NCBP1 RPL11 ATM RRS1 SF1 CD2BP2 SRSF5 TXNL4A SART1 |
| | 9.72E-21 | 59 | 248 | 4.60 | GO:0071826 ribonucleoprotein complex subunit organization | SFSWAP EIF4B DDX20 NLE1 RPS5 RPL6 RPLP0 RPL3 PRPF6 CRNKL1 RPS19 NOP53 EIF4H RPS15 RPL5 MRPS7 SNRPD2 LSM4 ERAL1 CDC73 SNRNP200 ABT1 LUC7L2 CELF1 RPS14 PRPF18 RAMAC RPS27 DDX28 CLNS1A EIF3I SF3B2 SF3A1 EIF3D EIF3E SNRPB EIF3G DDX46 DDX23 EIF3F SF3A3 EIF3C HSP90AB1 PRPF3 RPL13A ADAR SNU13 YJU2 RPL24 FASTKD2 NCBP1 RPL11 ATM RRS1 SF1 CD2BP2 SRSF5 TXNL4A SART1 |
| | 5.50E-27 | 77 | 327 | 4.55 | GO:0042254 ribosome biogenesis | MTREX RRP12 RIOK2 BUD23 NLE1 RPS5 DIMT1 NOP14 DDX18 RPL6 RPLP0 SNU13 RPL3 RPS16 RPS19 NOP53 DDX49 NUP88 BYSL RPS15 NOL10 UTP25 RPL5 DDX54 RRP36 MRPS7 NSUN5 DKC1 ERAL1 WDR74 LTV1 NOB1 RPS8 NHP2 ABT1 RPL7L1 NOM1 RPL7 NIFK MTERF3 TSR2 RPL26 ABCE1 LSM6 RPS14 DCAF13 BMS1 TSR1 MRPL1 RSL1D1 TRMT112 SART1 RPS27 RRS1 DDX28 RPL14 MAK16 RPL10A DDX47 EXOSC5 DDX17 GTPBP10 RPL24 FASTKD2 UTP20 MYBBP1A DDX56 CUL4A EIF4A3 RPP30 UTP14A CUL4B SLX9 MRPS2 RPL27 RPL11 XRN2 |
| | 1.25E-38 | 115 | 509 | 4.37 | GO:0022613 ribonucleoprotein complex biogenesis | MTREX RRP12 RIOK2 SFSWAP EIF4B DDX20 BUD23 NLE1 RPS5 DIMT1 NOP14 DDX18 RPL6 RPLP0 SNU13 RPL3 PRPF6 CRNKL1 RPS16 RPS19 NOP53 DDX49 EIF4H NUP88 BYSL RPS15 NOL10 UTP25 RPL5 DDX54 RRP36 MRPS7 SNRPD2 NSUN5 LSM4 DKC1 ERAL1 WDR74 CDC73 LTV1 NOB1 RPS8 SNRNP200 NHP2 ABT1 RPL7L1 NOM1 LUC7L2 RPL7 CELF1 NIFK MTERF3 TSR2 RPL26 ABCE1 LSM6 RPS14 DCAF13 PRPF18 BMS1 TSR1 MRPL1 RAMAC RSL1D1 TRMT112 SART1 RPS27 RRS1 DDX28 RPL14 MAK16 RPL10A DDX47 CLNS1A EIF3I SF3B2 SF3A1 EIF3D EIF3E SNRPB EIF3G DDX46 DDX23 EIF3F SF3A3 EIF3C HSP90AB1 PRPF3 RPL13A ADAR EXOSC5 DDX17 YJU2 GTPBP10 RPL24 FASTKD2 UTP20 MYBBP1A DDX56 CUL4A EIF4A3 RPP30 UTP14A CUL4B SLX9 MRPS2 RPL27 NCBP1 RPL11 ATM SF1 CD2BP2 XRN2 SRSF5 TXNL4A |

| | | | | | | |
|---|---|---|---|---|---|---|
| | 1.94E-31 | 133 | 784 | 3.28 | GO:0006412 translation | EIF4B RPL31 PUM3 RPS5 EIF3I RPL6 RPLP0 EIF3D EARS2 EIF3E EIF4H CNOT2 MRPL51 MRPL2 RARS1 RPL24 AARS2 UPF3B MRPS7 RPL36 DPH2 DPH6 NARS1 RPL13A MRPS5 LARP1 EEF1A1 EIF4A1 RPL26 RPL29 MRPL55 RPL22L1 ABCE1 RPS14 MRPL16 EEF2 MRPL13 MRPL52 MRPL11 EIF3F FARSA MRPL41 EIF3C RACK1 MRPL43 MRPS18A RPL3 RPS16 RPS19 RPL18A RPL28 RPL34 MRPL18 MRPL3 RPS15 MRPL19 MRPL37 RPL21 MRPS2 RPL5 RPL23 EIF3G RPL27 MRPL35 DAP3 MRPL50 NCBP1 MRPL15 RPL11 RPS8 MRPL9 RPL7 MRPL49 EIF4E MRPL39 RPL30 MRPL17 RPL27A RPL13 MRPL1 RPL4 AURKAIP1 RPS27 GADD45GIP1 MRPL14 MRPL30 RPL14 RPS26 RPL10A MRPL38 MRPL12 UPF1 CSDE1 EIF2AK2 YBX1 PABPC1 BZW1 PUS7 CIRBP EIF4G2 FASTKD2 TARDBP SYNCRIP PML EIF4A3 MOV10 SLBP TYMS CALR GIGYF2 DHFR NOA1 DNAJC3 QKI ZNF706 DNAJC1 DIS3L2 CDKAL1 EIF1AD TCEA1 TPR TCOF1 PLD1 EXOSC5 NSUN5 TRUB2 COA3 PLXNB2 CELF1 CDC123 DHPS RRBP1 NACA |
| | 5.11E-31 | 135 | 814 | 3.21 | GO:0043043 peptide biosynthetic proc. | EIF4B RPL31 PUM3 RPS5 EIF3I RPL6 RPLP0 EIF3D EARS2 EIF3E EIF4H CNOT2 MRPL51 MRPL2 RARS1 RPL24 AARS2 UPF3B MRPS7 RPL36 DPH2 DPH6 NARS1 RPL13A MRPS5 LARP1 EEF1A1 AASDH EIF4A1 RPL26 RPL29 MRPL55 RPL22L1 ABCE1 RPS14 MRPL16 EEF2 MRPL13 MRPL52 MRPL11 EIF3F FARSA MRPL41 EIF3C RACK1 MRPL43 MRPS18A RPL3 RPS16 RPS19 RPL18A RPL28 RPL34 MRPL18 MRPL3 RPS15 MRPL19 MRPL37 RPL21 MRPS2 RPL5 RPL23 EIF3G RPL27 MRPL35 DAP3 MRPL50 NCBP1 MRPL15 RPL11 RPS8 MRPL9 RPL7 MRPL49 EIF4E MRPL39 RPL30 MRPL17 RPL27A RPL13 MRPL1 RPL4 AURKAIP1 RPS27 GADD45GIP1 MRPL14 MRPL30 RPL14 RPS26 RPL10A MRPL38 MRPL12 UPF1 CSDE1 EIF2AK2 YBX1 PABPC1 BZW1 PUS7 CIRBP GGT1 EIF4G2 FASTKD2 TARDBP SYNCRIP PML EIF4A3 MOV10 SLBP TYMS CALR GIGYF2 DHFR NOA1 DNAJC3 QKI ZNF706 DNAJC1 DIS3L2 CDKAL1 EIF1AD TCEA1 TPR TCOF1 PLD1 EXOSC5 NSUN5 TRUB2 COA3 PLXNB2 CELF1 CDC123 DHPS RRBP1 NACA |
| | 7.84E-35 | 167 | 1082 | 2.98 | GO:0006396 RNA processing | ELAC2 ZCCHC8 MTREX RRP12 RIOK2 SFSWAP DDX20 THUMPD1 GPKOW BUD23 CLNS1A EXOSC5 DIMT1 |

| | | | | | | |
|---|---|---|---|---|---|---|
| | | | | | | SRRT NOP14 SF3B2 DDX18 PAPOLA CIRBP SNU13 DDX17 SRSF5 PRPF6 CRNKL1 DTWD1 RPS16 YJU2 NOP53 DDX49 EFTUD2 QKI BYSL NOL10 SRSF7 PRPF3 UTP25 FASTKD2 CPSF3 TRMT1L SMU1 RBM19 DDX54 RRP36 AARS2 SNRPD2 SYMPK SNRPB NSUN5 LSM4 DKC1 GRSF1 THUMPD3 CDC73 RBM17 PRPF4 NCBP1 MFAP1 NOB1 RPRD1A RPS8 SNRNP200 DDX46 NHP2 CDKAL1 ABT1 RPL7L1 LUC7L2 RPL7 RPP30 TUT1 CELF1 MBNL1 NIFK TSR2 CCAR2 SSU72 ADAR RAVER1 RBM15 SLBP LSM6 RPS14 DCAF13 PRPF18 BMS1 TRUB1 CPSF2 TSR1 SF1 MRPL1 RAMAC HNRNPF RSL1D1 TRMT112 CTU2 DDX23 SART1 AURKAIP1 PUS1 RRS1 SF3A3 PRPF40A MAK16 UBL5 RPL10A RBMXL1 DDX47 RBM12 PABPC1 SF3A1 DDX5 KAT2A TAF12 SYNCRIP EIF4A3 TXNL4A PUS7 TCERG1 TARDBP RBM39 YRDC RPS26 YBX1 CHERP XRN2 RBM27 PABPN1 HNRNPUL1 SUGP1 RBM28 SUPT6H TCP1 UTP20 ECD ZC3H13 RPL23 LGALS3 DDX56 RBM26 NONO UTP14A SLX9 EIF4A1 U2SURP METTL2B TRUB2 CD2BP2 RPL4 RACK1 HMGN1 PRDX6 PAF1 RPS19 RPS15 IVNS1ABP RPL5 RPL27 WDR74 NCOR1 RPL11 RPL26 STAT3 MAP2K1 RPS27 RPL14 NCOR2 HNRNPDL |
| | 5.45E-31 | 152 | 999 | 2.94 | GO:0006518 peptide metabolic proc. | EIF4B RPL31 PUM3 RPS5 EIF3I RPL6 RPLP0 GGT1 EIF3D HM13 EARS2 EIF3E GSR EIF4H CNOT2 MRPL51 MRPL2 RARS1 RPL24 SPCS2 AARS2 UPF3B MRPS7 RPL36 DPH2 DPH6 GSTM3 NARS1 NPEPPS RPL13A MRPS5 LARP1 EEF1A1 AASDH EIF4A1 RPL26 RPL29 MRPL55 RPL22L1 ABCE1 ERAP2 RPS14 SEC11C MRPL16 EEF2 MRPL13 MRPL52 MRPL11 EIF3F FARSA MRPL41 EIF3C GSTK1 RACK1 GSTM2 MRPL43 MRPS18A RPL3 RPS16 RPS19 RPL18A RPL28 RPL34 MRPL18 MRPL3 RPS15 MRPL19 MRPL37 RPL21 MRPS2 RPL5 RPL23 EIF3G RPL27 MRPL35 DAP3 MRPL50 NCBP1 MRPL15 RPL11 RPS8 MRPL9 RPL7 MRPL49 EIF4E MRPL39 RPL30 MRPL17 RPL27A RPL13 MRPL1 RPL4 AURKAIP1 RPS27 GADD45GIP1 MRPL14 MRPL30 RPL14 RPS26 RPL10A MRPL38 MRPL12 UPF1 CSDE1 EIF2AK2 YBX1 PABPC1 BZW1 PUS7 CIRBP CLN5 EIF4G2 FASTKD2 CLU TARDBP RTN3 SYNCRIP PML EIF4A3 MOV10 SLBP TYMS CALR GIGYF2 DHFR NOA1 DNAJC3 SOD2 QKI ZNF706 |

| | | | | | | |
|---|---|---|---|---|---|---|
| | | | | | | DNAJC1 DIS3L2 ARL6IP5 CDKAL1 CLIC3 EIF1AD TCEA1 TPR TCOF1 PLD1 EXOSC5 NSUN5 YIPF5 TRUB2 COA3 PLXNB2 CELF1 CDC123 DHPS RRBP1 NACA ERO1A |
| | 3.03E-29 | 145 | 959 | 2.92 | GO:0043604 amide biosynthetic proc. | EIF4B RPL31 PUM3 RPS5 EIF3I RPL6 RPLP0 EIF3D EARS2 EIF3E EIF4H CNOT2 MRPL51 MRPL2 RARS1 RPL24 AARS2 UPF3B MRPS7 RPL36 ASS1 ACSS2 ACLY DPH2 DPH6 NARS1 RPL13A MRPS5 LARP1 EEF1A1 AASDH EIF4A1 RPL26 RPL29 MRPL55 RPL22L1 ABCE1 RPS14 SMPD1 MRPL16 EEF2 MRPL13 SPTLC3 MRPL52 MRPL11 EIF3F FARSA MRPL41 EIF3C ACSL5 RACK1 MRPL43 MRPS18A RPL3 RPS16 RPS19 RPL18A RPL28 RPL34 PDHX MRPL18 MRPL3 RPS15 MRPL19 MRPL37 RPL21 MRPS2 RPL5 RPL23 EIF3G RPL27 MRPL35 DAP3 MRPL50 NCBP1 MRPL15 RPL11 RPS8 MRPL9 RPL7 MRPL49 EIF4E MRPL39 RPL30 MRPL17 RPL27A RPL13 MRPL1 RPL4 AURKAIP1 RPS27 GADD45GIP1 MRPL14 MRPL30 RPL14 RPS26 RPL10A MRPL38 MRPL12 UPF1 CSDE1 EIF2AK2 YBX1 PABPC1 BZW1 PUS7 CIRBP GGT1 EIF4G2 FASTKD2 TARDBP SYNCRIP PML EIF4A3 MOV10 SLBP TYMS CALR GIGYF2 DHFR NOA1 DNAJC3 QKI ZNF706 ASL DNAJC1 DIS3L2 CDKAL1 EIF1AD TCEA1 TPR TCOF1 PLD1 EXOSC5 NSUN5 TRUB2 COA3 PLXNB2 CELF1 CDC123 DHPS TECR SLC25A1 RRBP1 NACA |
| | 7.10E-31 | 178 | 1301 | 2.64 | GO:0043603 cellular amide metabolic proc. | EIF4B RPL31 TIGAR PUM3 RPS5 EIF3I RPL6 RPLP0 GGT1 EIF3D HM13 GLA EARS2 EIF3E GSR EIF4H CNOT2 MRPL51 MRPL2 RARS1 RPL24 SPCS2 AARS2 UPF3B MRPS7 RPL36 ASS1 ACSS2 ACLY DPH2 DPH6 GSTM3 NARS1 NPEPPS RPL13A MRPS5 LARP1 EEF1A1 AASDH EIF4A1 RPL26 RPL29 MRPL55 GNPDA2 RPL22L1 ABCE1 ERAP2 RPS14 SMPD1 SEC11C MRPL16 EEF2 MRPL13 SPTLC3 MRPL52 MRPL11 EIF3F FARSA MRPL41 EIF3C ACSL5 GSTK1 RACK1 GSTM2 DHFR MRPL43 MRPS18A RPL3 RPS16 RPS19 RPL18A RPL28 RPL34 PDHX MRPL18 MRPL3 RPS15 MRPL19 MRPL37 RPL21 MRPS2 RPL5 RPL23 EIF3G RPL27 MRPL35 DAP3 MRPL50 NCBP1 MRPL15 RPL11 RPS8 MRPL9 RPL7 MRPL49 EIF4E MRPL39 RPL30 MRPL17 RPL27A RPL13 MRPL1 RPL4 AURKAIP1 RPS27 GADD45GIP1 MRPL14 MRPL30 |

| | | | | | | |
|---|---|---|---|---|---|---|
| | | | | | | RPL14 RPS26 RPL10A MRPL38 MRPL12 UPF1 CSDE1 EIF2AK2 YBX1 PABPC1 NT5C2 BZW1 PUS7 CIRBP CLN5 OGDH EIF4G2 FASTKD2 CLU TARDBP RTN3 HSD17B4 SYNCRIP PML EIF4A3 MOV10 SLBP NNMT TYMS CALR GIGYF2 NOA1 DNAJC3 CMAS SOD2 QKI CERT1 ZNF706 NAGK ASL DNAJC1 DIS3L2 ARL6IP5 CDKAL1 CLIC3 EIF1AD TCEA1 ADA TPR TCOF1 PLD1 EXOSC5 NSUN5 YIPF5 TRUB2 COA3 PLXNB2 ABHD4 CELF1 CDC123 DHPS TECR SLC25A1 RRBP1 FPGS AASDHPPT GNE ACSF2 NACA ERO1A |
| | 1.97E-24 | 161 | 1259 | 2.47 | GO:0034645 cellular macromolecule biosynthetic proc. | ALG1 EIF4B RPL31 PUM3 RPS5 EIF3I RPL6 RPLP0 EIF3D PMM1 EARS2 EIF3E EIF4H CNOT2 MRPL51 MRPL2 RARS1 RPL24 AARS2 UPF3B MRPS7 DAD1 RPL36 GFPT2 INPP5K DPH2 DPH6 NARS1 ITM2C UGGT1 RPL13A MRPS5 B3GAT3 HS2ST1 LARP1 EEF1A1 EIF4A1 RPL26 RPL29 MRPL55 STT3B RPL22L1 RPN1 ABCE1 RPS14 MRPL16 EEF2 MRPL13 MRPL52 DPY19L1 MRPL11 EIF3F FARSA MRPL41 EIF3C WDR45 GFPT1 ZDHHC18 RACK1 MRPL43 MRPS18A RPL3 RPS16 RPS19 RPL18A RPL28 RPL34 MRPL18 MRPL3 RPS15 MRPL19 MRPL37 RPL21 MRPS2 RPL5 RPL23 EIF3G RPL27 MRPL35 DAP3 MRPL50 NCBP1 MRPL15 RPL11 RPS8 MRPL9 RPL7 MRPL49 EIF4E MRPL39 RPL30 MRPL17 RPL27A RPL13 MRPL1 RPL4 AURKAIP1 RPS27 GADD45GIP1 MRPL14 MRPL30 RPL14 RPS26 RPL10A MRPL38 MRPL12 UPF1 CSDE1 EIF2AK2 YBX1 PABPC1 BZW1 PUS7 CIRBP EIF4G2 FASTKD2 TARDBP TMEM258 SYNCRIP PML NPC1 EIF4A3 GALNT2 MOV10 DBI SLBP TYMS ULK1 CALR GIGYF2 DHFR NOA1 DNAJC3 UGDH QKI EDEM3 ZNF706 DNAJC1 DIS3L2 CDKAL1 UGP2 EIF1AD TCEA1 FUT11 TREX1 PGM3 TPR TCOF1 PLD1 EXOSC5 NSUN5 ABCB10 TRUB2 COA3 PLXNB2 CELF1 CDC123 DHPS RRBP1 NUDT14 NACA |
| | 1.89E-24 | 210 | 1888 | 2.15 | GO:1901566 organonitrogen compound biosynthetic proc. | ALG1 EIF4B RPL31 HACD3 PUM3 RPS5 EIF3I RPL6 RPLP0 EIF3D PMM1 SMS EARS2 QPRT EIF3E EIF4H UGDH CHKA CNOT2 MRPL51 LPCAT3 MRPL2 RARS1 RPL24 GLS MTR AARS2 UPF3B MRPS7 UROD ASL DUT DAD1 RPL36 ASS1 ACSS2 GFPT2 ACLY DPH2 DPH6 NARS1 ITM2C UGGT1 FPGS RPL13A MRPS5 ALAD AASDHPPT B3GAT3 QDPR HS2ST1 LARP1 EEF1A1 AASDH EIF4A1 RPL26 RPL29 MRPL55 STT3B |

| | | | | | | |
|---|---|---|---|---|---|---|
| | | | | | | RPL22L1 RPN1 ABCE1 RPS14 SMPD1 MRPL16 EEF2 ATP5PD MRPL13 SPTLC3 MRPL52 NMNAT1 DPY19L1 MRPL11 EIF3F TYMS IMPDH2 FARSA MRPL41 EIF3C ADA WDR45 ACSL5 GFPT1 PAPSS2 ZDHHC18 RACK1 DHFR HMBS MRPL43 SDHA MRPS18A RPL3 RPS16 RPS19 RPL18A RPL28 RPL34 PDHX MRPL18 MRPL3 RPS15 MRPL19 MRPL37 SDHB RPL21 MRPS2 RPL5 RPL23 EIF3G RPL27 MRPL35 DAP3 MRPL50 NCBP1 MRPL15 RPL11 RPS8 MRPL9 RPL7 MRPL49 EIF4E MRPL39 RPL30 MRPL17 NDUFB8 RPL27A RPL13 MRPL1 RPL4 AURKAIP1 RPS27 GADD45GIP1 MRPL14 MRPL30 RPL14 RPS26 RPL10A MRPL38 MRPL12 UPF1 CSDE1 EIF2AK2 YBX1 PABPC1 BZW1 PUS7 CIRBP GGT1 NFKB1 EIF4G2 KYNU FASTKD2 MTHFD1L TARDBP TMEM258 SYNCRIP PML CLTC NPC1 EIF4A3 GALNT2 GLUD1 PLOD2 MOV10 DBI SLBP ULK1 CALR GIGYF2 OAT NOA1 DNAJC3 NME3 QKI ATP6V1A EDEM3 ZNF706 ABCB10 DNAJC1 DIS3L2 CDKAL1 CAPN2 ALDH7A1 NNMT EIF1AD TCEA1 FUT11 UCKL1 UQCC3 TREX1 PGM3 TPR TCOF1 PLD1 EXOSC5 NSUN5 PARP1 TRUB2 COA3 PLXNB2 CELF1 CDC123 DHPS TECR SLC25A1 SLC25A16 RRBP1 SLC26A2 NUDT14 NACA |
| | 3.16E-21 | 320 | 3630 | 1.70 | GO:0044085 cellular component biogenesis | NDUFAF7 HCCS SCIN ANLN MTREX TPR LIMA1 RRP12 RIOK2 SFSWAP EIF4B DDX20 BUD23 NLE1 PLD1 RPS5 MAPRE3 DIMT1 NOP14 DDX18 TPX2 RPL6 RPLP0 SNAP23 EZR SNAP29 SNU13 RPL3 PRPF6 CRNKL1 PLS3 RPS16 RPS19 NOP53 DDX49 CAV1 EIF4H NUP88 BYSL RPS15 NOL10 ALMS1 UTP25 HELLS TAF12 RPL5 DDX54 RRP36 MRPS7 SNRPD2 VASP MACF1 NSUN5 LSM4 TUBGCP2 DKC1 TUBG1 ERAL1 WDR74 CDC73 LTV1 MICAL1 KIF23 KIF11 FGD4 GTF2A2 NOB1 CLTC AFG3L2 RPS8 SNRNP200 PKP4 NHP2 ABT1 RPL7L1 NOM1 LUC7L2 RPL7 CELF1 INCENP HAUS1 NIFK MTERF3 AIFM1 SASS6 NRG1 TSR2 CHAF1B RPL26 ABCE1 LSM6 RPS14 RAD21 DNAAF5 DCAF13 PRPF18 BMS1 TSR1 MRPL1 RAMAC RSL1D1 CLSTN1 ATPAF2 SYNPO LAMB2 TRMT112 CSRP2 CKAP5 SART1 EPS8L2 ULK1 RPS27 TAF7 RRS1 TDRKH DDX28 COA3 RPL14 WDR45 H2BC12 MAK16 RPL10A PRC1 ATG9A UQCC3 DDX47 MAP1LC3B2 H4C1 CLNS1A CHMP2B EIF3I SF3B2 SF3A1 EIF3D |

| | | | | | | |
|---|---|---|---|---|---|---|
| | | | | | | VPS18 EIF3E SNRPB CHMP2A EIF3G AP3B1 VPS36 MVB12A DDX46 VIPAS39 VPS37A MED27 AP1G1 VPS39 VPS37C DDX23 EIF3F AURKB AP1S2 SF3A3 EIF3C BAK1 ME1 UBE2K EPS15 DNM1L FKBP1A OAS1 HSP90AB1 NUP93 CTCF STUB1 KAT2A UGDH ECT2 GLS PRPF3 SLC2A1 CLU CBX3 CDK2 MAP7D3 LATS1 LGALS3 INPP5K H3-3B RTN3 MORC2 YME1L1 ANP32B PML RPL13A MCU ADAR RACGAP1 TK1 COA6 MAT2A UBE2C SSNA1 FARSA ISG15 S100A10 RACK1 PAFAH1B1 EIF2AK2 TRIP13 EXOSC5 MTMR2 ICAM1 DDX17 PACSIN2 CORO1A PLAT YJU2 GTPBP10 DVL1 CNTNAP1 TCIRG1 LPCAT3 LOX RPL24 SPTBN1 CDC20 CTSD FASTKD2 STBD1 DYNC2I2 SET UTP20 ID1 TRAF2 TMEM175 DNMT1 MAP1B MYBBP1A SWAP70 DDX56 PDCL EMILIN1 RAP1GDS1 G3BP2 CUL4A EIF4A3 PARP1 RPP30 KCTD14 BAG3 DAB2 UTP14A CUL4B SLX9 INPPL1 ZFYVE1 H1-4 CDK1 HEG1 CALR H1-10 H1-0 TSPYL1 PLXNB2 SPTAN1 CYRIA CTR9 E2F4 EZH2 PHLDB2 CSDE1 NUDCD3 CUL3 NDC1 MYLK RAB27A SMC1A PXN KIF4A AAAS MISP TM9SF4 STAG2 SMC3 DUSP3 CNOT2 KIF14 MRPS2 NUP153 RPL27 ARFIP2 TERF2 NCBP1 NUMA1 NCOR1 RPL11 RALB G3BP1 ATM AHCTF1 FBLIM1 NDUFB8 ATG4B MYADM MYO1C LCMT1 SEC22B SOD2 CUTC ALAD GPC1 SLC9A1 CIRBP SRP54 ARHGEF6 CDC123 AKAP13 CD47 CENPF TCP1 SF1 CD2BP2 XRN2 ITGA6 SRSF5 RP2 DNA2 TXNL4A |
| | 1.17E-19 | 351 | 4206 | 1.61 | GO:0006996 organelle organization | SMARCD2 SMARCC2 LMAN1 NPEPPS SAE1 UBL5 NDUFAF7 RECQL SCIN PAFAH1B1 NCAPD2 ANLN TACC3 CUL7 TPR LIMA1 NDC1 ELMO2 TRAM1 TRIP13 SMC1A NLE1 MCM6 RPS5 MAPRE3 DNM1L TPX2 EPB41L1 RPL6 RPLP0 KIF4A SNAP23 EZR MAST3 SNAP29 PACSIN2 RPL3 SEC23A STAG2 PLS3 KATNAL1 CLN5 CORO1A CTCF VPS18 RPS19 NOP53 SMC3 KAT2A VAT1 SUPT6H SEC24A SMC4 RPS15 SPTBN1 ALMS1 ABCD3 DYNC2I2 HELLS NCAPH NAA50 RPL5 CBX3 CIT OPTN MRPS7 VASP MGME1 MACF1 CCDC32 MAP7D3 MAP1S LSM4 TUBGCP2 TUBG1 MAP1B TOP2A CHD1L PEX11B ARFIP2 CLUH RPA1 ERAL1 TERF2 CCNB1 TIMM10 ESPL1 MICAL1 CKAP2 HUS1 YME1L1 TOR1B TUBB2A NUMA1 |

| | | | | | | |
|---|---|---|---|---|---|---|
| | | | | | | NUSAP1 KIF23 KIF11 SEC31A NUP54 SHROOM3 FGD4 PML POLG PHLDB2 YIPF5 ABT1 NCAPG2 SH3KBP1 MKI67 ARFGAP2 ATM INCENP VPS51 SLC25A4 HAUS1 UTRN BUB3 MTERF3 AIFM1 SASS6 MYO1E CHAF1B NUP35 MAD2L1 RPS14 RAD21 DNAAF5 SUN1 RAB8B MCM7 VPS39 MAP1A H1-4 MFF TOR1AIP2 MFN1 ESCO2 ATPAF2 SYNPO UBE2C CSRP2 CKAP5 TUBB6 SEC24C ULK1 RPS27 AURKB TDRKH DDX28 SMTN COA3 H1-10 H1-0 WDR45 SPTAN1 MYO1C H2BC12 PRC1 ATG9A UQCC3 HMGN1 MAP1LC3B2 TUBB3 ANXA8L1 ANXA8 SEC22B H4C1 BAZ1B BAK1 ATP6V0A1 CHMP2B TCIRG1 ATP6V1A CDC20 CLU CHMP2A DKC1 MYBBP1A AP3B1 VPS36 MVB12A NHP2 TIMM8B VIPAS39 VPS37A NSMCE2 AP1G1 VPS37C AP1S2 UPF1 STEEP1 MTA3 SEC61A1 HSP90AB1 MCM5 NAA10 RBBP7 NUP93 EZH2 CHKA MCM3 CDK2 MTERF1 LATS1 INPP5K H3-3B RTN3 MORC2 EMC7 ETS1 ANP32B DNA2 KIF2C CHEK1 PALM2AKAP2 CCAR2 RACGAP1 RBM15 MMGT1 SSNA1 MTA1 ANKFY1 S100A10 SAMD9 RFC2 LETMD1 BUD23 TIGAR RBL1 SLC9A1 DAAM1 ARMCX3 DVL1 CNTNAP1 NSD2 LPCAT3 SOD2 CAP2 CERT1 ECT2 RPL24 KDM3A HMGCL CTSD STBD1 SET MASTL MORF4L2 ID1 TMEM175 DNMT1 PCNA SWAP70 FBH1 SMYD5 SERPINE2 FLNB TBC1D4 RAP1GDS1 G3BP2 USO1 AEBP2 USP3 NCOR1 AFG3L2 RETREG2 DCAF1 G3BP1 BAG3 RNF20 CAPN2 MEAF6 RFC4 ZFYVE1 SF1 CDK1 JMJD1C KDM2A TBL1XR1 CALR MORF4L1 GTF2F2 TSPYL1 WDR5 AFAP1 SIPA1L1 CYRIA CTR9 E2F4 TREX1 UHRF1 GGCT CSDE1 NUDCD3 CUL3 RIOK2 RAB27A MAVS PXN AAAS MSH2 MISP CNOT2 PTK7 MSH3 LMNB1 CCT4 DOCK7 KIF14 FASTKD2 TCP1 TARDBP MRPS2 NUP153 GNAI1 RAB30 CLTC ARHGDIA RPL11 PARP1 RALB DIS3L2 TRIM36 AHCTF1 MCU SMG1 GNL3 INPPL1 NDUFB8 ATG4B RRS1 MYADM PTPN1 PRPF40A MYO18A MCMBP LCMT1 CIRBP CAV1 BST2 DOP1B AKAP13 CD47 ZFYVE16 CENPF HMGA1 AURKAIP1 EPS15 RP2 HIRIP3 MAP2K1 |
| KEGG | 1.46E-28 | 51 | 134 | 7.36 | Path:hsa03010 Ribosome | MRPL3 RPL22L1 RPL13A RPL36 MRPL13 MRPL18 MRPL15 RPL10A MRPL2 MRPS7 MRPS2 MRPL30 |

| | | | | | | |
|---|---|---|---|---|---|---|
| | | | | | | MRPL35 MRPL16 MRPS18A RPL3 RPL4 RPL5 RPL6 RPL7 RPL11 RPL13 RPL18A RPL21 RPL24 RPL26 RPL27 RPL30 RPL27A RPL28 RPL29 RPL31 RPL34 RPLP0 MRPL12 RPS5 RPS8 RPS14 RPS15 RPS16 RPS19 RPS26 RPS27 MRPL17 MRPL14 MRPS5 MRPL11 MRPL9 MRPL1 RPL14 RPL23 |
| | 3.63E-04 | 8 | 23 | 6.72 | Path:hsa03430 Mismatch repair | MSH6 MSH2 MSH3 PCNA POLD2 RFC2 RFC4 RPA1 |
| | 4.28E-05 | 11 | 36 | 5.90 | Path:hsa03030 DNA replication | DNA2 MCM3 MCM5 MCM6 MCM7 PCNA POLD2 PRIM1 RFC2 RFC4 RPA1 |
| | 3.63E-04 | 10 | 37 | 5.22 | Path:hsa01250 Biosynthesis of nucleotide sugars | GFPT1 PGM3 PMM1 NANS NAGK CMAS UGDH UGP2 UXS1 HKDC1 |
| | 3.20E-05 | 13 | 49 | 5.13 | Path:hsa00520 Amino sugar and nucleotide sugar metabolism | GNE GNPDA2 GFPT1 CYB5R1 PGM3 PMM1 NANS NAGK CMAS UGDH UGP2 UXS1 HKDC1 |
| | 4.27E-11 | 31 | 131 | 4.57 | Path:hsa03040 Spliceosome | SF3A1 CHERP TXNL4A TCERG1 SF3A3 SF3B2 LSM6 DDX42 DDX5 SNRNP200 U2SURP PRPF6 LSM4 NCBP1 SNU13 PCBP1 CRNKL1 PRPF40A SRSF5 SRSF7 SNRPB SNRPD2 RBM17 PRPF18 SART1 PRPF4 PRPF3 EFTUD2 DDX23 EIF4A3 DDX46 |
| | 6.81E-04 | 11 | 48 | 4.43 | Path:hsa00280 Valine leucine and isoleucine degradation | ECHS1 ACAD8 ACAA1 HADHA HADHB HMGCL ACADSB ALDH6A1 MMUT ALDH7A1 AACS |
| | 5.54E-08 | 24 | 108 | 4.29 | Path:hsa03013 Nucleocytoplasmic transport | NUP50 NUP35 EEF1A1 NUP160 NUP188 AHCTF1 KPNA2 TNPO1 NCBP1 NUP88 IPO11 NUP54 NDC1 RANBP2 RANGAP1 UPF1 UPF3B TPR NUP37 NUP214 AAAS NUP93 EIF4A3 NUP153 |
| | 2.30E-11 | 36 | 169 | 4.12 | Path:hsa04141 Protein processing in endoplasmic reticulum | STUB1 SEC23A SEC24A CKAP4 DAD1 STT3B HSPA4L SEC31A TRAM1 ERLEC1 SEC61A1 ERO1A HSP90AB1 LMAN1 P4HB UBXN1 DNAJB11 EIF2AK2 DNAJC3 UGGT1 BAK1 RAD23A RPN1 RRBP1 DNAJC1 SSR4 TRAF2 UBE2D1 WFS1 EDEM3 CALR CAPN1 CAPN2 PLAA PDIA4 SEC24C |
| | 2.30E-11 | 43 | 232 | 3.58 | Path:hsa05171 Coronavirus disease-COVID-19 | ADAR RPL22L1 RPL13A RPL36 RPL10A NFKB1 OAS1 OAS3 EIF2AK2 MAVS RPL3 RPL4 RPL5 RPL6 RPL7 RPL11 RPL13 RPL18A RPL21 RPL24 RPL26 RPL27 RPL30 RPL27A RPL28 RPL29 RPL31 RPL34 RPLP0 RPS5 RPS8 RPS14 RPS15 RPS16 RPS19 RPS26 RPS27 STAT1 STAT3 CASP1 RPL14 RPL23 ISG15 |

| | | | | | | |
|---|---|---|---|---|---|---|
| | 5.27E-06 | 23 | 126 | 3.53 | Path:hsa04110 Cell cycle | CDK4 STAG2 CHEK1 E2F4 ORC6 MAD2L1 MCM3 MCM5 MCM6 MCM7 ATM PCNA RAD21 RBL1 CCND1 TFDP1 SMC1A CCNB1 SMC3 BUB3 ESPL1 CDK1 CDC20 |
| | 1.92E-03 | 17 | 115 | 2.86 | Path:hsa01200 Carbon metabolism | ECHS1 RPIA GLUD1 IDH2 IDH3A MDH1 ME1 ME2 ALDH6A1 MMUT OGDH PC ACSS2 SDHA SDHB HKDC1 ACOX3 |
| | 1.80E-03 | 21 | 159 | 2.55 | Path:hsa04217 Necroptosis | DNM1L PARP1 CHMP2B CHMP2A GLUD1 SLC25A4 XIAP HSP90AB1 EIF2AK2 PYGB PYGM SMPD1 STAT1 STAT3 STAT6 TRAF2 CAPN1 CAPN2 H2AC15 CASP1 AIFM1 |
| | 1.92E-03 | 25 | 210 | 2.30 | Path:hsa05170 Human immunodeficiency virus 1 infection | TAB1 CHEK1 AP1G1 GNAI1 GNAQ GNB2 GNG10 ATM NFKB1 CYCS PPP3CB GNG12 MAP2K1 BAK1 PXN BST2 TRAF2 CALR CUL4B CUL4A AP1S2 CCNB1 APOBEC3B DCAF1 CDK1 |
| | 1.47E-13 | 154 | 1538 | 1.94 | Path:hsa01100 Metabolic pathways | ADA GNE DHRS9 TCIRG1 LANCL1 BPNT1 CDIPT ATP5PD CHKA EARS2 COX4I1 GNPDA2 CMBL CYP51A1 DAD1 AKR1C1 DHCR7 DHCR24 DHFR DNMT1 DUT ECHS1 STT3B ALAD EZH2 FDPS SACM1L RPIA NT5C2 QPRT NNT ATP6V0A2 FPGS PLD3 GALNS GALNT2 MTHFD1L B3GAT3 GFPT1 GGT1 ACAD8 GK GLA GLS GLUD1 PGM2L1 GSR GSTM2 GSTM3 ACAA1 HADHA HADHB HCCS HMBS HMGCL MMAB HSD17B4 IDH2 IDH3A IDI1 ACADSB IMPA1 IMPDH2 INPPL1 GSTK1 PEDS1 MAT2A MDH1 ME1 MGST3 ALDH6A1 ASL ASS1 MTR MMUT ACLY NDUFB8 NME3 NNMT OAT OGDH ALDH7A1 P4HA1 PAFAH1B1 NSDHL PC DCXR HSD17B7 HACD3 ACSL5 INPP5K ATP6V1A PGM3 ATP6V1B2 ATP6V1C1 ATP6V1E1 PLD1 ATP6V0A1 PLOD2 PMM1 ACP5 NANS CYCS UCKL1 SPTLC3 NAGK ACSS2 CMAS ALG1 TIGAR PYGB PYGM SQOR QDPR AASDHPPT RPN1 SDHA SDHB RBKS BLVRA BLVRB NMNAT1 AACS SMPD1 SMS BPGM TK1 TST TYMS UGDH UGP2 UROD NSD2 CA9 GGCT GDPD3 L2HGDH SCD5 UXS1 HKDC1 PDHX ACOX3 PDE5A CBR3 MTMR2 KYNU P4HA2 SELENBP1 PAPSS2 TECR PTGES ATP6V1G1 PRDX6 ACYP1 |

**Suppl table 4: Protein pathways enriched in GO Molecular Function (MF). REACTOME and Wikipathway terms for cladribine deregulated proteins in SW1353 cell line**

| Databases | FDR | nGenes | Pathway Genes | Fold Enrichment | Terms | Genes |
|---|---|---|---|---|---|---|
| GO MF | 0.0123 | 2 | 19 | 72.99 | GO:0035173 histone kinase activity | AURKB CDK1 |
| | 0.0251 | 2 | 38 | 36.49 | GO:0042162 telomeric DNA binding | RPA1 HNRNPD |
| | 0.0123 | 5 | 476 | 7.28 | GO:0003779 actin binding | EPS8L2 ANLN BIN1 INF2 SMTN |
| | 0.0088 | 7 | 849 | 5.72 | GO:0016301 kinase activity | AXL AURKB GNE TK1 CDK1 NRP1 SKP2 |
| | 0.0088 | 8 | 1008 | 5.50 | GO:0016772 transferase activity transferring phosphorus-containing groups | AXL AURKB CMAS GNE TK1 CDK1 NRP1 SKP2 |
| | 0.0088 | 8 | 1053 | 5.27 | GO:0008092 cytoskeletal protein binding | BIN1 EPS8L2 ANLN RAB11FIP5 KIF11 AXL INF2 SMTN |
| | 0.0123 | 9 | 1662 | 3.75 | GO:0005524 ATP binding | LIG1 KIF11 GNE UBE2J2 HSPA4L AXL TK1 CDK1 AURKB |
| | 0.0123 | 9 | 1729 | 3.61 | GO:0032559 adenyl ribonucleotide binding | LIG1 KIF11 GNE UBE2J2 HSPA4L AXL TK1 CDK1 AURKB |
| | 0.0123 | 9 | 1741 | 3.58 | GO:0030554 adenyl nucleotide binding | LIG1 KIF11 GNE UBE2J2 HSPA4L AXL TK1 CDK1 AURKB |
| | 0.0088 | 11 | 2237 | 3.41 | GO:0019899 enzyme binding | HACD2 NRP1 RAB11FIP5 BIN1 HNRNPD INF2 SERPINE1 CCNB1 KIF11 UBE2J2 AURKB |
| | 0.0282 | 9 | 2034 | 3.07 | GO:0035639 purine ribonucleoside triphosphate binding | LIG1 KIF11 GNE UBE2J2 HSPA4L AXL TK1 CDK1 AURKB |
| | 0.0282 | 9 | 2106 | 2.96 | GO:0032555 purine ribonucleotide binding | LIG1 KIF11 GNE UBE2J2 HSPA4L AXL TK1 CDK1 AURKB |
| | 0.0282 | 9 | 2120 | 2.94 | GO:0017076 purine nucleotide binding | LIG1 KIF11 GNE UBE2J2 HSPA4L AXL TK1 CDK1 AURKB |
| | 0.0282 | 9 | 2123 | 2.94 | GO:0032553 ribonucleotide binding | LIG1 KIF11 GNE UBE2J2 HSPA4L AXL TK1 CDK1 AURKB |
| | 0.0282 | 10 | 2553 | 2.72 | GO:0016740 transferase activity | AXL AURKB CMAS GNE UBE2J2 TK1 CDK1 DDB2 NRP1 SKP2 |
| REACTOME | 0.0007 | 2 | 5 | 277.35 | R-HSA-69478 G2/M DNA replication checkpoint | CCNB1 CDK1 |
| | 0.0009 | 2 | 6 | 231.12 | R-HSA-176417 Phosphorylation of Emi1 | CCNB1 CDK1 |
| | 0.0011 | 2 | 7 | 198.10 | R-HSA-2980767 Activation of NIMA Kinases NEK9 NEK6 NEK7 | CCNB1 CDK1 |
| | 0.0017 | 2 | 9 | 154.08 | R-HSA-113507 E2F-enabled inhibition of pre-replication complex formation | CCNB1 CDK1 |

| | | | | | | |
|---|---|---|---|---|---|---|
| | 0.0020 | 2 | 10 | 138.67 | R-HSA-2465910 MASTL Facilitates Mitotic Progression | CCNB1 CDK1 |
| | 0.0034 | 2 | 14 | 99.05 | R-HSA-162658 Golgi Cisternae Pericentriolar Stack Reorganization | CCNB1 CDK1 |
| | 0.0034 | 2 | 14 | 99.05 | R-HSA-75035 Chk1/Chk2Cds1 mediated inactivation of Cyclin B:Cdk1 complex | CCNB1 CDK1 |
| | 0.0039 | 2 | 16 | 86.67 | R-HSA-5358606 Mismatch repair MMR directed by MSH2:MSH3 MutSbeta | LIG1 RPA1 |
| | 0.0006 | 3 | 29 | 71.73 | R-HSA-69205 G1/S-Specific Transcription | TK1 CDK1 RRM2 |
| | 0.0000 | 6 | 153 | 27.19 | R-HSA-69206 G1/S Transition | RPA1 CCNB1 SKP2 TK1 CDK1 RRM2 |
| | 0.0007 | 4 | 108 | 25.68 | R-HSA-174143 APC/C-mediated degradation of cell cycle proteins | CCNB1 SKP2 CDK1 AURKB |
| | 0.0007 | 4 | 108 | 25.68 | R-HSA-453276 Reg. of mitotic cell cycle | CCNB1 SKP2 CDK1 AURKB |
| | 0.0000 | 6 | 172 | 24.19 | R-HSA-453279 Mitotic G1 phase and G1/S transition | RPA1 CCNB1 SKP2 TK1 CDK1 RRM2 |
| | 0.0002 | 8 | 622 | 8.92 | R-HSA-69278 Cell Cycle Mitotic | LIG1 RPA1 CCNB1 SKP2 TK1 CDK1 RRM2 AURKB |
| | 0.0006 | 8 | 774 | 7.17 | R-HSA-1640170 Cell Cycle | LIG1 RPA1 CCNB1 SKP2 TK1 CDK1 RRM2 AURKB |
| Wiki-pathway | 0.0047 | 2 | 23 | 60.29 | WP531 DNA mismatch repair | RPA1 LIG1 |
| | 0.0012 | 3 | 43 | 48.37 | WP4753 Nucleotide excision repair | LIG1 DDB2 RPA1 |
| | 0.0000 | 6 | 90 | 46.22 | WP2446 Retinoblastoma gene in cancer | SKP2 RRM2 ANLN RPA1 CCNB1 CDK1 |
| | 0.0062 | 2 | 31 | 44.73 | WP3982 miRNA reg. of p53 pathway in prostate cancer | SERPINE1 DDB2 |
| | 0.0063 | 2 | 34 | 40.79 | WP5117 Cohesin complex - Cornelia de Lange syndrome | AURKB CDK1 |
| | 0.0063 | 2 | 34 | 40.79 | WP1601 Fluoropyrimidine activity | TK1 RRM2 |
| | 0.0070 | 2 | 37 | 37.48 | WP5269 Genetic causes of PSVD/INCPH | GTF2I SHCBP1 |
| | 0.0076 | 2 | 40 | 34.67 | WP2516 ATM signaling pathway | CCNB1 CDK1 |
| | 0.0024 | 3 | 64 | 32.50 | WP45 G1 to S cell cycle control | RPA1 CCNB1 CDK1 |
| | 0.0024 | 3 | 68 | 30.59 | WP707 DNA damage response | CCNB1 DDB2 CDK1 |
| | 0.0025 | 3 | 74 | 28.11 | WP5114 Nucleotide excision repair in xeroderma pigmentosum | RPA1 LIG1 DDB2 |

| | | | | | | |
|---|---|---|---|---|---|---|
| | 0.0041 | 3 | 98 | 21.23 | WP1530 miRNA reg. of DNA damage response | CCNB1 DDB2 CDK1 |
| | 0.0051 | 3 | 119 | 17.48 | WP4946 DNA repair pathways full network | RPA1 DDB2 LIG1 |
| | 0.0051 | 3 | 120 | 17.33 | WP179 Cell cycle | SKP2 CCNB1 CDK1 |
| | 0.0033 | 4 | 218 | 12.72 | WP5115 Network map of SARS-CoV-2 signaling pathway | RRM2 SERPINE1 CDK1 CCNB1 |

**Suppl table 5: Protein pathways enriched in GO Molecular Function (MF). REACTOME and Wikipathway terms for cladribine deregulated proteins in JJ012 cell line**

| Data bases | FDR | nGenes | Pathway Genes | Fold Enrichment | Terms | Genes |
|---|---|---|---|---|---|---|
| GO MF | 9.77E-29 | 62 | 201 | 5.96 | GO:0003735 structural constituent of ribosome | RPS5 RPS16 RPS19 RPS15 RPS8 RPS14 RPS26 MRPL43 RPL31 RPL6 RPLP0 MRPS18A RPL3 MRPL51 MRPL2 RPL24 MRPL3 MRPL19 RPL21 MRPS2 RPL5 MRPS7 RPL23 RPL36 RPL27 DAP3 MRPL15 RPL13A RPL11 MRPS5 RPL7L1 RPL7 MRPL49 RPL30 MRPL17 RPL26 RPL29 MRPL55 RPL22L1 RPL27A MRPL16 RPL13 MRPL13 RPL4 MRPL11 RPS27 MRPL41 ISG15 RPL14 RPL18A RPL28 RPL34 RPL10A MRPL18 MRPL37 MRPL35 MRPL9 MRPL1 MRPL52 MRPL14 MRPL30 MRPL12 |
| | 5.67E-77 | 305 | 1852 | 3.18 | GO:0003723 RNA binding | UPF1 ELAC2 CSDE1 ZCCHC8 MTREX YTHDC2 POLR2B TPR FAM120A RRP12 EIF2AK2 MRPL43 EIF4B YBX1 THUMPD1 HEATR6 PABPC1 TCOF1 RPL31 BUD23 SMC1A CLNS1A PUM3 BZW1 TNPO1 RPS5 NOA1 CHERP DIMT1 SRRT NOP14 SF3B2 DDX18 XRN2 RPL6 RPLP0 RBM27 PUS7 EZR ARCN1 HSP90AB1 CIRBP POLRMT SF3A1 SNU13 DDX17 RPL3 EIF3D SRSF5 PABPN1 SRP54 PRPF6 CRNKL1 CORO1A EIF3E RPS16 CCDC9 HNRNPUL1 RPS19 NOP53 RPL18A DDX49 SUGP1 GTPBP10 RBM28 EIF4H EDF1 RPL28 DDX5 EFTUD2 SUPT6H CCDC86 EIF4G2 QKI BYSL MRPL2 TCERG1 RPL24 MRPL3 RPS15 SPTBN1 CCT4 NOL10 SRSF7 MRPL37 PRPF3 UTP25 FASTKD2 ALDH6A1 TCP1 UTP20 TARDBP TRMT1L RPL21 ANXA11 RPL5 RBM19 DDX54 ZC3H13 RRP36 UPF3B MRPS7 GTF2F1 RPL23 SNRPD2 SNRPB RRBP1 KTN1 MACF1 MTERF1 DUT RPL36 BST2 NSUN5 LSM4 HELZ2 ASS1 EIF3G DKC1 RBM39 RPL27 ZCCHC9 TOP2A |

| | | | | | | |
|---|---|---|---|---|---|---|
| | | | | | | LGALS3 MYBBP1A GRSF1 ANKRD17 ERAL1 DAP3 TPT1 CCDC59 DHX34 SYNCRIP CKAP4 FLNB DDX56 NCBP1 MRPL15 NUSAP1 HADHB G3BP2 USO1 RBM26 MFAP1 CLTC EIF4A3 RPL13A NOSIP RPL11 RPS8 MRPL9 PARP1 SNRNP200 MRPS5 MANF BTF3 DDX46 G3BP1 NHP2 ABT1 RPL7L1 C7orf50 NOM1 LUC7L2 NONO RPL7 RPP30 MKI67 CELF1 SERPINH1 EIF4E MBNL1 HNRNPDL MRPL39 MOV10 NIFK LARP1 PDIA4 RPL30 EEF1A1 UTP14A SMG1 CCAR2 ADAR FDPS RAVER1 EIF4A1 RPL26 RPL29 FUBP1 RBM15 RBM47 U2SURP RPN1 GNL3 SLBP LSM6 RPS14 DCAF13 STRBP BMS1 RPL27A TRUB2 RPL13 EEF2 TSR1 SF1 COA6 H1-4 MRPL1 PCBP1 RAMAC HNRNPF RSL1D1 MRPL13 DDX23 RPL4 MRPL11 SART1 C11orf68 PUS1 CAVIN1 RPS27 RRS1 FARSA CALR APOBEC3B MRPL14 MRPL41 KPNA2 DDX28 EWSR1 SF3A3 EIF3C H1-10 CCDC137 P4HB GPATCH8 RPS19BP1 RPL14 H1-0 SUPT5H PTPN1 PRPF40A MYO18A PCBP2 RPS26 MAK16 DDX42 RPL10A GIGYF2 RACK1 DDX47 NOL7 RBM12 NOL12 H4C1 DDX20 EIF3I OAS1 MRPS18A OAS3 MRPL18 CPSF3 IFIT3 AARS2 CLUH PRPF4 DNA2 RPL22L1 CPSF2 MRPL16 HEXIM1 YRDC GPKOW RANGAP1 TST DIS3L2 TUT1 RNF20 TRUB1 TYMS DHFR SFSWAP EXOSC5 PAPOLA RBBP7 EARS2 EZH2 RARS1 DNMT1 THUMPD3 RBM17 PDCD4 RANBP2 NABP1 CTU2 EIF1AD IMPDH2 TDRKH RBMXL1 PPME1 STAT3 RPL34 MRPL12 |
| | 9.84E-11 | 54 | 355 | 2.94 | GO:0045296 cadherin binding | ANLN LIMA1 EPN2 BZW1 CHMP2B EPS15 RPL6 ITGA6 EZR HSP90AB1 TBC1D10A PACSIN2 RANGAP1 EIF3E NDRG1 PPP1R13L EIF4H RPL34 EIF4G2 RARS1 RPL24 SPTBN1 STAT1 PRDX6 VASP KTN1 MACF1 ARFIP2 SWAP70 CEMIP2 FLNB NIBAN2 USO1 PHLDB2 BAG3 LARP1 RPL29 GAPVD1 EEF2 PCBP1 RSL1D1 CKAP5 ARHGAP1 EPS8L2 H1-10 RPL14 PTPN1 SPTAN1 RPS26 GIGYF2 RACK1 PPME1 PKP4 CTNNAL1 |
| | 7.62E-18 | 117 | 908 | 2.49 | GO:0005198 structural molecule activity | RPS5 RPS16 RPS19 RPS15 COL4A2 RPS8 RPS14 COL4A1 RPS26 NUP160 TPR MRPL43 RPL31 RPL6 RPLP0 NUP188 MRPS18A RPL3 NUP93 MRPL51 MRPL2 RPL24 MRPL3 MRPL19 RPL21 MRPS2 RPL5 |

| | | | | | | |
|---|---|---|---|---|---|---|
| | | | | | | NUP153 MRPS7 RPL23 NUP214 MACF1 RPL36 TUBG1 RPL27 DAP3 TUBB2A MRPL15 EMILIN1 SEC31A NUP54 RPL13A RPL11 MRPS5 RPL7L1 RPL7 MRPL49 RPL30 MRPL17 RPL26 RPL29 MRPL55 NUP35 RPL22L1 FBN1 RPL27A MRPL16 RPL13 LAMB2 MRPL13 RPL4 MRPL11 CSRP2 TUBB6 RPS27 MRPL41 ISG15 RPL14 MYL6B TUBB3 RPL18A RPL28 RPL34 HMGCL RPL10A EPB41L1 NUP88 MRPL18 SPTBN1 MRPL37 COPA COPB1 MRPL35 H3-3B CLTC MRPL9 COPG2 H1-4 MRPL1 MRPL52 CTBP2 MRPL14 COPG1 COPB2 H1-10 MRPL30 H1-0 H2BC12 MRPL12 H2AC15 H4C1 NDC1 COL6A3 UPF3B MAP1B MAP1A PCOLCE MFGE8 ECM1 HAPLN1 IGFBP7 EDIL3 LMNB1 HMGA1 NUMA1 SMTN SPTAN1 |
| | 2.09E-11 | 84 | 687 | 2.36 | GO:0140640 catalytic activity acting on a nucleic acid | ELAC2 DIMT1 XRN2 NOB1 RECQL UPF1 DNASE1L1 MTREX TDP1 POLR2B RFC2 DDX20 BUD23 MCM6 UNG POLRMT DDX17 MCM5 POLR2C EARS2 DTWD1 POLR2I SMC3 DDX5 MCM3 RARS1 CPSF3 TRMT1L POLM AARS2 MGME1 HELZ2 TERF2 THUMPD3 NARS1 FBH1 ERCC5 DNA2 POLG PTRH2 EIF4A3 SNRNP200 DIS3L2 RFC4 METTL2B MCM7 FARSA PRIM1 TREX1 YTHDC2 MSH2 PLD3 MSH6 DDX54 NSUN5 DNMT1 DKC1 TOP2A PCNA HMGA1 G3BP1 RPP30 TUT1 MOV10 DDX42 DDX18 DDX49 MSH3 HELLS CHD1L DHX34 DDX56 DDX46 CDKAL1 EIF4A1 DDX23 PUS1 DDX28 GTF2F2 MYO18A DDX47 CNOT2 EXOSC5 POLR3G |
| | 1.60E-18 | 228 | 2381 | 1.85 | GO:0000166 nucleotide binding | SDHA ACOX3 HSP90AB1 GSR GTPBP10 ACBD5 NNT UXS1 GBP1 ABCD3 AARS2 RRAS GFER POR TUBG1 CHD1L TUBB2A SQOR RALB QDPR BAG3 DBI AIFM1 CYB5R1 ABCE1 BMS1 CLPX TSR1 TUBB6 ANXA6 ECI2 DHFR TUBB3 ACSS2 MTREX RAB27B RAB27A NT5C2 NOA1 MSH2 SRP54 OAS3 PCYOX1 MSH6 MTHFD1L RAB9A NQO2 LATS1 MORC2 MICAL1 EIF4A3 PPIP5K2 GLUD1 CBR3 COQ8A STK17A RAB8B RAB6A IMPDH2 SULT1A1 MYO18A PRIM1 TXNRD1 UHRF1 CAMKK1 RECQL UPF1 BAZ1B YTHDC2 RFC2 EIF2AK2 RIOK2 DDX20 MYLK ME1 PYGM TRIP13 UBE2D1 SMC1A STK10 MCM6 UBE2T UBE2K AACS ME2 HADHA DNM1L DDX18 OAS1 PAPOLA KIF4A MAST3 DDX17 MCM5 |

| | | | | | | |
|---|---|---|---|---|---|---|
| | | | | | | RP2 KATNAL1 NME3 EARS2 CSK DDX49 BLVRA SMC3 DDX5 EFTUD2 UGDH CHKA MCM3 PTK7 MSH3 RARS1 SMC4 ATP6V1A NRBP1 CCT4 AAK1 DHCR24 KIF14 HELLS TCP1 MASTL PYROXD1 CIT DDX54 CDK2 NAGK UBA2 GNAI1 HELZ2 ASS1 DNMT1 ACLY TOP2A ERAL1 DAP3 SWAP70 IRAK2 CAMK1 DPH6 NARS1 FBH1 DHX34 CDK4 ABCB10 ITM2C DDX56 YME1L1 TOR1B FPGS RAB30 KIF23 KIF11 DNA2 PDE5A SEPTIN11 MMAB AFG3L2 KIF2C PARP1 SNRNP200 DCAF1 DDX46 G3BP1 GNA12 ATP6V1B2 MKI67 TUT1 ATM CHEK1 ACAD8 MRPL39 MOV10 GNAQ EEF1A1 HKDC1 SMG1 AASDH MYO1E GNE EIF4A1 GTPBP8 RFC4 GNL3 HSPA4L HSPA12A ILK IDH3A MCM7 SLFN5 ACSF2 AXL EEF2 TK1 MAT2A MAP2K1 UGP2 CDK1 MFN1 RBKS PC NMNAT1 DDX23 CTBP2 UBE2C YES1 ULK1 AURKB FARSA IDH2 UBA7 DDX28 ACBD3 CAMK1D SEPTIN10 GTF2F2 ACADSB ACSL5 MYO1C ERO1A DDX42 UCKL1 PAPSS2 GK TREX1 DDX47 ALDH6A1 ETFDH DHCR7 ARFIP2 |
| | 1.60E-18 | 228 | 2382 | 1.85 | GO:1901265 nucleoside phosphate binding | SDHA ACOX3 HSP90AB1 GSR GTPBP10 ACBD5 NNT UXS1 GBP1 ABCD3 AARS2 RRAS GFER POR TUBG1 CHD1L TUBB2A SQOR RALB QDPR BAG3 DBI AIFM1 CYB5R1 ABCE1 BMS1 CLPX TSR1 TUBB6 ANXA6 ECI2 DHFR TUBB3 ACSS2 MTREX RAB27B RAB27A NT5C2 NOA1 MSH2 SRP54 OAS3 PCYOX1 MSH6 MTHFD1L RAB9A NQO2 LATS1 MORC2 MICAL1 EIF4A3 PPIP5K2 GLUD1 CBR3 COQ8A STK17A RAB8B RAB6A IMPDH2 SULT1A1 MYO18A PRIM1 TXNRD1 UHRF1 CAMKK1 RECQL UPF1 BAZ1B YTHDC2 RFC2 EIF2AK2 RIOK2 DDX20 MYLK ME1 PYGM TRIP13 UBE2D1 SMC1A STK10 MCM6 UBE2T UBE2K AACS ME2 HADHA DNM1L DDX18 OAS1 PAPOLA KIF4A MAST3 DDX17 MCM5 RP2 KATNAL1 NME3 EARS2 CSK DDX49 BLVRA SMC3 DDX5 EFTUD2 UGDH CHKA MCM3 PTK7 MSH3 RARS1 SMC4 ATP6V1A NRBP1 CCT4 AAK1 DHCR24 KIF14 HELLS TCP1 MASTL PYROXD1 CIT DDX54 CDK2 NAGK UBA2 GNAI1 HELZ2 ASS1 DNMT1 ACLY TOP2A ERAL1 DAP3 SWAP70 IRAK2 CAMK1 DPH6 NARS1 FBH1 DHX34 CDK4 ABCB10 ITM2C DDX56 YME1L1 TOR1B FPGS RAB30 KIF23 |

| | | | | | | |
|---|---|---|---|---|---|---|
| | | | | | | KIF11 DNA2 PDE5A SEPTIN11 MMAB AFG3L2 KIF2C PARP1 SNRNP200 DCAF1 DDX46 G3BP1 GNA12 ATP6V1B2 MKI67 TUT1 ATM CHEK1 ACAD8 MRPL39 MOV10 GNAQ EEF1A1 HKDC1 SMG1 AASDH MYO1E GNE EIF4A1 GTPBP8 RFC4 GNL3 HSPA4L HSPA12A ILK IDH3A MCM7 SLFN5 ACSF2 AXL EEF2 TK1 MAT2A MAP2K1 UGP2 CDK1 MFN1 RBKS PC NMNAT1 DDX23 CTBP2 UBE2C YES1 ULK1 AURKB FARSA IDH2 UBA7 DDX28 ACBD3 CAMK1D SEPTIN10 GTF2F2 ACADSB ACSL5 MYO1C ERO1A DDX42 UCKL1 PAPSS2 GK TREX1 DDX47 ALDH6A1 ETFDH DHCR7 ARFIP2 |
| | 1.56E-20 | 259 | 2743 | 1.82 | GO:0036094 small molecule binding | OAT PYGM SDHA LMAN1 ACOX3 HSP90AB1 PYGB GSR GTPBP10 ACBD5 NNT UXS1 GBP1 ABCD3 AARS2 RRAS GFER POR TUBG1 CHD1L TUBB2A SQOR CRABP2 RALB QDPR BAG3 DDAH1 DBI AIFM1 CYB5R1 ABCE1 BMS1 CLPX TSR1 TUBB6 OSBP2 ANXA6 ECI2 DHFR TUBB3 ACSS2 TYMS STARD3NL NPC1L1 OSBPL5 MTREX RAB27B SLC2A3 RAB27A NT5C2 NOA1 MSH2 SRP54 CLN5 OGDH ENG OAS3 PCYOX1 MSH6 MTHFD1L RAB9A NQO2 LATS1 MORC2 MICAL1 MMAB NPC1 EIF4A3 PPIP5K2 MMUT GLUD1 CBR3 COQ8A STK17A RAB8B RAB6A IMPDH2 RXRA SULT1A1 MYO18A PRIM1 TXNRD1 UHRF1 CAMKK1 RECQL UPF1 BAZ1B YTHDC2 POLR2B RFC2 EIF2AK2 RIOK2 DDX20 MYLK ME1 TRIP13 UBE2D1 SMC1A P4HA2 STK10 MCM6 UBE2T UBE2K AACS ME2 HADHA DNM1L DDX18 OAS1 PAPOLA P3H2 KIF4A MAST3 DDX17 MCM5 RP2 KATNAL1 NME3 EARS2 CDIPT CSK DDX49 BLVRA SMC3 DDX5 EFTUD2 UGDH CHKA SOD2 MCM3 PTK7 MSH3 RARS1 SMC4 ATP6V1A NRBP1 CCT4 KYNU AAK1 DHCR24 MTR P3H1 MTARC2 KIF14 HELLS TCP1 MASTL PYROXD1 P4HA1 CIT DDX54 CDK2 NAGK UBA2 GNAI1 HELZ2 ASS1 DNMT1 ACLY TOP2A ERAL1 DAP3 SWAP70 IRAK2 CAMK1 DPH6 NARS1 FBH1 DHX34 CDK4 ABCB10 ITM2C DDX56 YME1L1 TOR1B FPGS RAB30 KIF23 KIF11 DNA2 PDE5A SEPTIN11 AFG3L2 KIF2C PARP1 SNRNP200 DCAF1 DDX46 G3BP1 GNA12 ATP6V1B2 MKI67 TUT1 ATM CHEK1 ACAD8 PLOD2 RPIA MRPL39 MOV10 GNAQ |

| | | | | | | |
|---|---|---|---|---|---|---|
| | | | | | | EEF1A1 HKDC1 SMG1 AASDH MYO1E GNE EIF4A1 GTPBP8 RFC4 GNL3 HSPA4L HSPA12A ILK IDH3A MCM7 SLFN5 ACSF2 AXL EEF2 TK1 MAT2A MAP2K1 UGP2 CDK1 MFN1 RBKS SPTLC3 PC NMNAT1 DDX23 CTBP2 UBE2C YES1 ULK1 AURKB FARSA IDH2 UBA7 DDX28 ACBD3 CAMK1D SEPTIN10 GTF2F2 ACADSB ACSL5 MYO1C ERO1A DDX42 UCKL1 PAPSS2 GK TREX1 DDX47 ALDH6A1 ETFDH DHCR7 CAV1 ARFIP2 |
| | 6.45E-34 | 407 | 4400 | 1.79 | GO:0003676 nucleic acid binding | UPF1 ELAC2 CSDE1 ZCCHC8 MTREX YTHDC2 POLR2B TPR FAM120A RRP12 EIF2AK2 MRPL43 EIF4B YBX1 THUMPD1 HEATR6 PABPC1 TCOF1 RPL31 BUD23 SMC1A CLNS1A PUM3 BZW1 TNPO1 RPS5 NOA1 CHERP DIMT1 SRRT NOP14 SF3B2 DDX18 XRN2 RPL6 RPLP0 RBM27 PUS7 EZR ARCN1 HSP90AB1 CIRBP POLRMT SF3A1 SNU13 DDX17 RPL3 EIF3D SRSF5 PABPN1 SRP54 PRPF6 CRNKL1 CORO1A EIF3E RPS16 CCDC9 HNRNPUL1 RPS19 NOP53 RPL18A DDX49 SUGP1 GTPBP10 RBM28 EIF4H EDF1 RPL28 SMARCD2 DDX5 EFTUD2 SUPT6H CCDC86 EIF4G2 QKI BYSL MRPL2 TCERG1 RPL24 MRPL3 RPS15 SPTBN1 CCT4 NOL10 SRSF7 MRPL37 PRPF3 UTP25 FASTKD2 ALDH6A1 TCP1 UTP20 TARDBP TRMT1L RPL21 ANXA11 RPL5 RBM19 DDX54 ZC3H13 RRP36 UPF3B MRPS7 GTF2F1 RPL23 SNRPD2 SNRPB RRBP1 KTN1 MACF1 MTERF1 DUT RPL36 BST2 NSUN5 LSM4 HELZ2 ASS1 EIF3G DKC1 RBM39 RPL27 ZCCHC9 TOP2A LGALS3 MYBBP1A GRSF1 ANKRD17 ERAL1 DAP3 TPT1 CCDC59 DHX34 SYNCRIP CKAP4 FLNB DDX56 NCBP1 MRPL15 NUSAP1 HADHB G3BP2 USO1 SMARCC2 RBM26 MFAP1 CLTC EIF4A3 RPL13A NOSIP RPL11 RPS8 MRPL9 PARP1 SNRNP200 MRPS5 MANF BTF3 DDX46 G3BP1 NHP2 ABT1 RPL7L1 C7orf50 NOM1 LUC7L2 NONO RPL7 RPP30 MKI67 CELF1 SERPINH1 EIF4E MBNL1 HNRNPDL MRPL39 MOV10 NIFK LARP1 PDIA4 RPL30 EEF1A1 UTP14A SMG1 CCAR2 ADAR FDPS RAVER1 EIF4A1 RPL26 RPL29 FUBP1 RBM15 RBM47 U2SURP RPN1 GNL3 SLBP LSM6 RPS14 DCAF13 STRBP BMS1 RPL27A TRUB2 RPL13 EEF2 TSR1 SF1 COA6 H1-4 MRPL1 PCBP1 RAMAC |

| | | | | | | |
|---|---|---|---|---|---|---|
| | | | | | | HNRNPF RSL1D1 MRPL13 DDX23 RPL4 MRPL11 SART1 C11orf68 PUS1 CAVIN1 RPS27 RRS1 FARSA CALR APOBEC3B MRPL14 MRPL41 KPNA2 DDX28 EWSR1 SF3A3 EIF3C H1-10 CCDC137 P4HB GPATCH8 RPS19BP1 RPL14 H1-0 SUPT5H PTPN1 PRPF40A MYO18A PCBP2 RPS26 MAK16 DDX42 RPL10A GIGYF2 RACK1 DDX47 NOL7 RBM12 NOL12 H4C1 DNASE1L1 TDP1 DDX20 MCM10 MCM6 NFKB2 RBL1 EIF3I OAS1 TOX4 WDR76 MRPS18A MCM5 GMEB2 CTCF SMC3 NFKB1 OAS3 TIMELESS MRPL18 MCM3 MSH3 SMAD5 STAT1 KDM3A MSH6 CPSF3 IFIT3 TAF12 AARS2 CLUH RPA1 TERF2 ERCC5 PRPF4 DNA2 NUP35 RPL22L1 RYBP CPSF2 MCM7 STAT6 MRPL16 THAP11 STAT3 JMJD1C NABP1 EIF3F BASP1 ZNF114 RXRA HEXIM1 YRDC H2BC12 TFDP1 E2F4 CUX1 H2AC15 GTF3C5 RECQL RTF2 GPKOW UNG CLSPN MSH2 RANGAP1 EZH2 ELK3 PRDX5 TST MAP1S DNMT1 H3-3B PCNA FBH1 ETS1 HMGA1 POLG DIS3L2 TUT1 RNF20 MTERF3 AIFM1 CUL4B TRUB1 FAM111A KDM2A TYMS TBL1XR1 TAF7 MTA1 DHFR UHRF1 RFC2 MTA3 SFSWAP PDCD2 EXOSC5 BRAP PAPOLA KIF4A ORC6 RBBP7 POLR2C EARS2 NUCB1 POLR2I POLD2 NSD2 SOD2 LMNB1 RARS1 SMC4 SET POLM NUP153 SWAP70 THUMPD3 NARS1 RBM17 DNAJC1 AEBP2 PML NCOR1 ZNF593 UHRF2 ATM PDCD4 RANBP2 AHCTF1 RFC4 RAD21 CTU2 EIF1AD IMPDH2 RAD23A TDRKH TCEA1 GTF2F2 NCOR2 NACA HMGN1 RBMXL1 TREX1 PPME1 RPL34 KIF2C MRPL12 |
| | 4.01E-14 | 204 | 2237 | 1.76 | GO:0019899 enzyme binding | PAF1 SPAG9 ATP6V0A1 CUL3 BOD1L1 MTA3 UBE2D1 RANGAP1 FAM83D TIMP1 CAV1 TCIRG1 CAP2 TCERG1 TRAF2 CDC73 CUL4A RPRD1A CUL4B CCAR2 GAPVD1 ARIH1 CAVIN2 RIN1 MTA1 ATP6V0A2 ANKFY1 RPS19BP1 ISG15 CTR9 RACK1 MAP1LC3B2 TPR HACD3 DNM1L SLC9A1 DMXL2 NDRG1 PPP3CB DUSP3 WFS1 DOCK7 CLU ECD ERCC5 ANP32B ITGAV EIF4E UBXN1 PPME1 CUL7 RFC2 DDX20 MCM10 TNPO1 IPO11 TPX2 HSP90AB1 TAB1 DAAM1 SRSF5 DNAJC3 STUB1 CSK PPP6R1 TAX1BP1 DVL1 KAT2A CTSC CCND1 SOD2 LMNB1 ECT2 STAT1 ERRFI1 SLC2A1 CDC20 |

| | | | | | | |
|---|---|---|---|---|---|---|
| | | | | | | POR TOP2A PCNA MICAL1 RAB11FIP5 KCNH1 PARP1 SH3KBP1 UBASH3B CAPN2 STAT3 TAF7 P4HB PTPN1 CYRIA CASP10 NDUFAF7 NPC1L1 YTHDC2 ELMO2 YBX1 PDCD2 SRI UBE2T UBE2K WDR70 MAPRE3 NOP14 MAVS PXN PUS7 WDR76 EZR MSH2 SNU13 HM13 XIAP HNRNPUL1 RPS19 SUFU MSH3 SMAD5 SPTBN1 MSH6 DHCR24 GBP1 PRDX6 KIF14 STBD1 TCP1 CSTA RPL5 CBX3 CIT OPTN SLC12A4 SERPINB6 GTF2F1 RPL23 CD70 UBA2 LATS1 ATP6V1E1 LGALS3 TERF2 AP3B1 CCNB1 GSTM3 ATP6V1G1 HMGA1 CASP1 BCAR3 KIF11 PML POLG NCOR1 CLTC SAE1 RPL11 ECM1 RALB GNA12 LUC7L2 TUT1 UTRN RANBP2 RNF20 EEF1A1 ADAMTS4 RACGAP1 ILK AP1G1 STAT6 STIM1 EEF2 TOR1AIP2 AKAP13 GNB2 PARP14 CTBP2 UBE2C ARHGAP1 YES1 ULK1 AURKB CALR RAD23A KPNA2 RXRA TSPYL1 SUPT5H NCOR2 WDR45 PCBP2 MYO1C PRC1 GSTM2 PPP1CB ITGA1 H2AC15 PAFAH1B1 PABPN1 MASTL ARPP19 FGD4 NRG1 ARFIP2 |
| | 3.64E-16 | 237 | 2630 | 1.74 | GO:0043168 anion binding | SCIN OAT PYGM SDHA ACOX3 HSP90AB1 PYGB GSR GTPBP10 CERT1 UXS1 GBP1 ABCD3 AARS2 RRAS GFER POR TUBG1 TUBB2A SQOR CRABP2 RALB QDPR AIFM1 CYB5R1 ABCE1 BMS1 CLPX TSR1 TUBB6 AKR1C1 WDR45 ANXA6 TUBB3 ACSS2 TYMS MTREX ZFYVE16 RAB27B RAB27A NT5C2 NOA1 MSH2 PACSIN2 SRP54 OGDH OAS3 LANCL1 PCYOX1 MSH6 MTHFD1L RAB9A NQO2 LATS1 MORC2 GSTM3 MICAL1 EIF4A3 PPIP5K2 PTGES GLUD1 CBR3 RACGAP1 COQ8A STK17A ZFYVE1 RAB8B RAB6A RXRA SULT1A1 MYO18A TXNRD1 UQCC3 GSTM2 DHFR ANXA8 CAMKK1 RECQL UPF1 BAZ1B YTHDC2 RFC2 EIF2AK2 RIOK2 DDX20 MYLK TRIP13 UBE2D1 SMC1A P4HA2 STK10 MCM6 UBE2T UBE2K AACS HADHA DNM1L DDX18 OAS1 PAPOLA P3H2 KIF4A MAST3 DDX17 MCM5 RP2 KATNAL1 NME3 EARS2 CSK DDX49 SMC3 DDX5 EFTUD2 CTSC CHKA MCM3 PTK7 MSH3 RARS1 SMC4 ATP6V1A NRBP1 CCT4 KYNU AAK1 DHCR24 P3H1 MTARC2 KIF14 HELLS TCP1 MASTL PYROXD1 P4HA1 CIT DDX54 CDK2 NAGK UBA2 GNAI1 HELZ2 ASS1 ACLY TOP2A CHD1L ERAL1 |

| | | | | | | |
|---|---|---|---|---|---|---|
| | | | | | | DAP3 SWAP70 IRAK2 CAMK1 DPH6 NARS1 FBH1 DHX34 CDK4 ABCB10 ITM2C DDX56 YME1L1 TOR1B FPGS RAB30 KIF23 KIF11 DNA2 PDE5A SEPTIN11 MMAB AFG3L2 KIF2C SNRNP200 NCEH1 DCAF1 DDX46 G3BP1 GNA12 ATP6V1B2 MKI67 TUT1 ATM CHEK1 ACAD8 PLOD2 MOV10 GNAQ EEF1A1 HKDC1 SMG1 AASDH MYO1E GNE EIF4A1 GTPBP8 RFC4 GNL3 HSPA4L HSPA12A ILK MCM7 SLFN5 ACSF2 AXL EEF2 TK1 MAT2A MAP2K1 CDK1 MFN1 RBKS SPTLC3 PC NMNAT1 DDX23 UBE2C YES1 ULK1 AURKB FARSA UBA7 DDX28 CAMK1D SEPTIN10 GTF2F2 ACADSB ACSL5 MYO1C ERO1A DDX42 UCKL1 PAPSS2 GK DDX47 ETFDH ME1 SLC9A1 ARFIP2 |
| | 1.52E-12 | 191 | 2123 | 1.74 | GO:0032553 ribonucleotide binding | HSP90AB1 GTPBP10 ACBD5 GBP1 ABCD3 AARS2 RRAS POR TUBG1 TUBB2A RALB DBI ABCE1 BMS1 CLPX TSR1 TUBB6 ANXA6 ECI2 TUBB3 ACSS2 MTREX RAB27B RAB27A NT5C2 NOA1 MSH2 SRP54 OAS3 MSH6 MTHFD1L RAB9A LATS1 MORC2 EIF4A3 PPIP5K2 GLUD1 COQ8A STK17A RAB8B RAB6A SULT1A1 MYO18A PRIM1 CAMKK1 RECQL UPF1 BAZ1B YTHDC2 RFC2 EIF2AK2 RIOK2 DDX20 MYLK TRIP13 UBE2D1 SMC1A STK10 MCM6 UBE2T UBE2K AACS HADHA DNM1L DDX18 OAS1 PAPOLA KIF4A MAST3 DDX17 MCM5 RP2 KATNAL1 NME3 EARS2 CSK DDX49 SMC3 DDX5 EFTUD2 CHKA MCM3 PTK7 MSH3 RARS1 SMC4 ATP6V1A NRBP1 CCT4 AAK1 KIF14 HELLS TCP1 MASTL CIT DDX54 CDK2 NAGK UBA2 GNAI1 HELZ2 ASS1 ACLY TOP2A CHD1L ERAL1 DAP3 SWAP70 IRAK2 CAMK1 DPH6 NARS1 FBH1 DHX34 CDK4 ABCB10 ITM2C DDX56 YME1L1 TOR1B FPGS RAB30 KIF23 KIF11 DNA2 PDE5A SEPTIN11 MMAB AFG3L2 KIF2C SNRNP200 DCAF1 DDX46 G3BP1 GNA12 ATP6V1B2 MKI67 TUT1 ATM CHEK1 MOV10 GNAQ EEF1A1 HKDC1 SMG1 AASDH MYO1E GNE EIF4A1 GTPBP8 RFC4 GNL3 HSPA4L HSPA12A ILK MCM7 SLFN5 ACSF2 AXL EEF2 TK1 MAT2A MAP2K1 UGP2 CDK1 MFN1 RBKS PC NMNAT1 DDX23 UBE2C YES1 ULK1 AURKB FARSA UBA7 DDX28 ACBD3 CAMK1D SEPTIN10 GTF2F2 |

| | | | | | | |
|---|---|---|---|---|---|---|
| | | | | | | ACSL5 MYO1C DDX42 UCKL1 PAPSS2 GK DDX47 ALDH6A1 ME1 ARFIP2 |
| | 2.38E-12 | 190 | 2120 | 1.73 | GO:0017076 purine nucleotide binding | HSP90AB1 GTPBP10 ACBD5 GBP1 ABCD3 AARS2 RRAS TUBG1 TUBB2A RALB BAG3 DBI ABCE1 BMS1 CLPX TSR1 TUBB6 ANXA6 ECI2 TUBB3 ACSS2 MTREX RAB27B RAB27A NT5C2 NOA1 MSH2 SRP54 OAS3 MSH6 MTHFD1L RAB9A LATS1 MORC2 EIF4A3 PPIP5K2 GLUD1 COQ8A STK17A RAB8B RAB6A SULT1A1 MYO18A CAMKK1 RECQL UPF1 BAZ1B YTHDC2 RFC2 EIF2AK2 RIOK2 DDX20 MYLK TRIP13 UBE2D1 SMC1A STK10 MCM6 UBE2T UBE2K AACS HADHA DNM1L DDX18 OAS1 PAPOLA KIF4A MAST3 DDX17 MCM5 RP2 KATNAL1 NME3 EARS2 CSK DDX49 SMC3 DDX5 EFTUD2 CHKA MCM3 PTK7 MSH3 RARS1 SMC4 ATP6V1A NRBP1 CCT4 AAK1 KIF14 HELLS TCP1 MASTL CIT DDX54 CDK2 NAGK UBA2 GNAI1 HELZ2 ASS1 ACLY TOP2A CHD1L ERAL1 DAP3 SWAP70 IRAK2 CAMK1 DPH6 NARS1 FBH1 DHX34 CDK4 ABCB10 ITM2C DDX56 YME1L1 TOR1B FPGS RAB30 KIF23 KIF11 DNA2 PDE5A SEPTIN11 MMAB AFG3L2 KIF2C SNRNP200 DCAF1 DDX46 G3BP1 GNA12 ATP6V1B2 MKI67 TUT1 ATM CHEK1 MOV10 GNAQ EEF1A1 HKDC1 SMG1 AASDH MYO1E GNE EIF4A1 GTPBP8 RFC4 GNL3 HSPA4L HSPA12A ILK MCM7 SLFN5 ACSF2 AXL EEF2 TK1 MAT2A MAP2K1 CDK1 MFN1 RBKS PC NMNAT1 DDX23 UBE2C YES1 ULK1 AURKB FARSA UBA7 DDX28 ACBD3 CAMK1D SEPTIN10 GTF2F2 ACSL5 MYO1C DDX42 UCKL1 PAPSS2 GK TREX1 DDX47 ALDH6A1 ME1 ARFIP2 |
| | 4.50E-12 | 188 | 2106 | 1.73 | GO:0032555 purine ribonucleotide binding | HSP90AB1 GTPBP10 ACBD5 GBP1 ABCD3 AARS2 RRAS TUBG1 TUBB2A RALB DBI ABCE1 BMS1 CLPX TSR1 TUBB6 ANXA6 ECI2 TUBB3 ACSS2 MTREX RAB27B RAB27A NT5C2 NOA1 MSH2 SRP54 OAS3 MSH6 MTHFD1L RAB9A LATS1 MORC2 EIF4A3 PPIP5K2 GLUD1 COQ8A STK17A RAB8B RAB6A SULT1A1 MYO18A CAMKK1 RECQL UPF1 BAZ1B YTHDC2 RFC2 EIF2AK2 RIOK2 DDX20 MYLK TRIP13 UBE2D1 SMC1A STK10 MCM6 UBE2T UBE2K AACS HADHA DNM1L DDX18 OAS1 PAPOLA KIF4A MAST3 DDX17 MCM5 RP2 KATNAL1 NME3 EARS2 |

| | | | | | | |
|---|---|---|---|---|---|---|
| | | | | | | CSK DDX49 SMC3 DDX5 EFTUD2 CHKA MCM3 PTK7 MSH3 RARS1 SMC4 ATP6V1A NRBP1 CCT4 AAK1 KIF14 HELLS TCP1 MASTL CIT DDX54 CDK2 NAGK UBA2 GNAI1 HELZ2 ASS1 ACLY TOP2A CHD1L ERAL1 DAP3 SWAP70 IRAK2 CAMK1 DPH6 NARS1 FBH1 DHX34 CDK4 ABCB10 ITM2C DDX56 YME1L1 TOR1B FPGS RAB30 KIF23 KIF11 DNA2 PDE5A SEPTIN11 MMAB AFG3L2 KIF2C SNRNP200 DCAF1 DDX46 G3BP1 GNA12 ATP6V1B2 MKI67 TUT1 ATM CHEK1 MOV10 GNAQ EEF1A1 HKDC1 SMG1 AASDH MYO1E GNE EIF4A1 GTPBP8 RFC4 GNL3 HSPA4L HSPA12A ILK MCM7 SLFN5 ACSF2 AXL EEF2 TK1 MAT2A MAP2K1 CDK1 MFN1 RBKS PC NMNAT1 DDX23 UBE2C YES1 ULK1 AURKB FARSA UBA7 DDX28 ACBD3 CAMK1D SEPTIN10 GTF2F2 ACSL5 MYO1C DDX42 UCKL1 PAPSS2 GK DDX47 ALDH6A1 ME1 ARFIP2 |
| | 6.36E-11 | 179 | 2034 | 1.70 | GO:0035639 purine ribonucleoside triphosphate binding | HSP90AB1 GTPBP10 GBP1 ABCD3 AARS2 RRAS TUBG1 TUBB2A RALB ABCE1 BMS1 CLPX TSR1 TUBB6 ANXA6 TUBB3 MTREX RAB27B RAB27A NT5C2 NOA1 MSH2 SRP54 OAS3 MSH6 MTHFD1L RAB9A LATS1 MORC2 EIF4A3 PPIP5K2 GLUD1 STK17A RAB6A MYO18A CAMKK1 RECQL UPF1 BAZ1B YTHDC2 RFC2 EIF2AK2 RIOK2 DDX20 MYLK TRIP13 UBE2D1 SMC1A STK10 MCM6 UBE2T UBE2K AACS DNM1L DDX18 OAS1 PAPOLA KIF4A MAST3 DDX17 MCM5 RP2 KATNAL1 NME3 EARS2 CSK DDX49 SMC3 DDX5 EFTUD2 CHKA MCM3 PTK7 MSH3 RARS1 SMC4 ATP6V1A NRBP1 CCT4 AAK1 KIF14 HELLS TCP1 MASTL CIT DDX54 CDK2 NAGK UBA2 GNAI1 HELZ2 ASS1 ACSS2 ACLY TOP2A CHD1L ERAL1 DAP3 SWAP70 IRAK2 CAMK1 DPH6 NARS1 FBH1 DHX34 CDK4 ABCB10 ITM2C DDX56 YME1L1 TOR1B FPGS RAB30 KIF23 KIF11 DNA2 SEPTIN11 MMAB AFG3L2 KIF2C SNRNP200 DCAF1 DDX46 G3BP1 GNA12 ATP6V1B2 MKI67 TUT1 ATM CHEK1 MOV10 GNAQ EEF1A1 HKDC1 SMG1 AASDH MYO1E GNE EIF4A1 COQ8A GTPBP8 RFC4 GNL3 HSPA4L HSPA12A RAB8B ILK MCM7 SLFN5 ACSF2 AXL EEF2 TK1 MAT2A MAP2K1 CDK1 MFN1 RBKS PC NMNAT1 DDX23 UBE2C YES1 ULK1 AURKB FARSA UBA7 DDX28 CAMK1D SEPTIN10 |

| | | | | | | |
|---|---|---|---|---|---|---|
| | | | | | | GTF2F2 ACSL5 MYO1C DDX42 UCKL1 PAPSS2 GK DDX47 ARFIP2 |
| REACTOME | 2.05E-29 | 55 | 153 | 6.95 | R-HSA-168273 Influenza Viral RNA Transcription and Replication | NUP160 POLR2B TPR NDC1 RPL31 NUP37 RPS5 RPL6 RPLP0 NUP50 AAAS NUP188 RPL3 DNAJC3 NUP93 POLR2C RPS16 POLR2I RPS19 RPL18A RPL28 NUP88 RPL34 RPL24 RPS15 RPL21 RPL5 NUP153 GTF2F1 RPL23 NUP214 RPL36 RPL27 GRSF1 NUP54 RPL13A RPL11 RPS8 PARP1 RPL7 RANBP2 RPL30 RPL26 RPL29 NUP35 RPL22L1 RPS14 RPL27A RPL13 RPL4 RPS27 GTF2F2 RPL14 RPS26 RPL10A |
| | 1.66E-22 | 44 | 129 | 6.59 | R-HSA-156827 L13a-mediated translational silencing of Ceruloplasmin expression | EIF4B PABPC1 RPL31 RPS5 EIF3I RPL6 RPLP0 RPL3 EIF3D EIF3E RPS16 RPS19 RPL18A EIF4H RPL28 RPL34 RPL24 RPS15 RPL21 RPL5 RPL23 RPL36 EIF3G RPL27 RPL13A RPL11 RPS8 RPL7 EIF4E RPL30 EIF4A1 RPL26 RPL29 RPL22L1 RPS14 RPL27A RPL13 RPL4 EIF3F RPS27 EIF3C RPL14 RPS26 RPL10A |
| | 5.19E-31 | 61 | 179 | 6.59 | R-HSA-168255 Influenza Infection | NUP160 POLR2B TPR EIF2AK2 NDC1 RPL31 NUP37 RPS5 RPL6 RPLP0 NUP50 AAAS NUP188 RPL3 PABPN1 DNAJC3 NUP93 POLR2C RPS16 POLR2I RPS19 RPL18A RPL28 NUP88 RPL34 RPL24 RPS15 RPL21 RPL5 NUP153 GTF2F1 RPL23 NUP214 RPL36 RPL27 GRSF1 NUP54 CLTC RPL13A RPL11 RPS8 PARP1 RPL7 RANBP2 RPL30 RPL26 RPL29 NUP35 RPL22L1 RPS14 RPL27A RPL13 RPL4 RPS27 CALR KPNA2 ISG15 GTF2F2 RPL14 RPS26 RPL10A |
| | 2.09E-21 | 43 | 130 | 6.39 | R-HSA-72706 GTP hydrolysis and joining of the 60S ribosomal subunit | EIF4B RPL31 RPS5 EIF3I RPL6 RPLP0 RPL3 EIF3D EIF3E RPS16 RPS19 RPL18A EIF4H RPL28 RPL34 RPL24 RPS15 RPL21 RPL5 RPL23 RPL36 EIF3G RPL27 RPL13A RPL11 RPS8 RPL7 EIF4E RPL30 EIF4A1 RPL26 RPL29 RPL22L1 RPS14 RPL27A RPL13 RPL4 EIF3F RPS27 EIF3C RPL14 RPS26 RPL10A |
| | 2.10E-21 | 44 | 137 | 6.21 | R-HSA-72613 Eukaryotic Translation Initiation | EIF4B PABPC1 RPL31 RPS5 EIF3I RPL6 RPLP0 RPL3 EIF3D EIF3E RPS16 RPS19 RPL18A EIF4H RPL28 RPL34 RPL24 RPS15 RPL21 RPL5 RPL23 RPL36 EIF3G RPL27 RPL13A RPL11 RPS8 RPL7 EIF4E RPL30 EIF4A1 RPL26 RPL29 RPL22L1 RPS14 RPL27A RPL13 RPL4 EIF3F RPS27 EIF3C RPL14 RPS26 RPL10A |

| | | | | | | |
|---|---|---|---|---|---|---|
| | 2.10E-21 | 44 | 137 | 6.21 | R-HSA-72737 Cap-dependent Translation Initiation | EIF4B PABPC1 RPL31 RPS5 EIF3I RPL6 RPLP0 RPL3 EIF3D EIF3E RPS16 RPS19 RPL18A EIF4H RPL28 RPL34 RPL24 RPS15 RPL21 RPL5 RPL23 RPL36 EIF3G RPL27 RPL13A RPL11 RPS8 RPL7 EIF4E RPL30 EIF4A1 RPL26 RPL29 RPL22L1 RPS14 RPL27A RPL13 RPL4 EIF3F RPS27 EIF3C RPL14 RPS26 RPL10A |
| | 4.41E-26 | 62 | 224 | 5.35 | R-HSA-8868773 rRNA processing in the nucleus and cytosol | MTREX RIOK2 THUMPD1 RPL31 BUD23 EXOSC5 RPS5 DIMT1 NOP14 XRN2 RPL6 RPLP0 SNU13 RPL3 RPS16 RPS19 RPL18A DDX49 RBM28 RPL28 RPL34 BYSL RPL24 RPS15 UTP25 UTP20 RPL21 RPL5 RRP36 RPL23 RPL36 DKC1 RPL27 LTV1 TEX10 NOB1 RPL13A RPL11 RPS8 NHP2 RPL7 RPP30 RPL30 UTP14A RPL26 RPL29 RPL22L1 GNL3 RPS14 DCAF13 BMS1 RPL27A RPL13 TSR1 TRMT112 RPL4 RPS27 RPL14 RPS26 RPL10A DDX47 NOL12 |
| | 3.36E-38 | 92 | 336 | 5.29 | R-HSA-72766 Translation | MRPL43 SEC61A1 EIF4B TRAM1 PABPC1 RPL31 RPS5 EIF3I RPL6 RPLP0 MRPS18A RPL3 EIF3D SRP54 EARS2 EIF3E RPS16 RPS19 RPL18A EIF4H RPL28 RPL34 MRPL51 MRPL18 MRPL2 RARS1 RPL24 MRPL3 RPS15 MRPL19 MRPL37 SPCS2 RPL21 MRPS2 RPL5 AARS2 MRPS7 RPL23 RPL36 EIF3G RPL27 MRPL35 ERAL1 DAP3 NARS1 MRPL50 MRPL15 RPL13A RPL11 RPS8 MRPL9 MRPS5 RPL7 MRPL49 EIF4E MRPL39 RPL30 EEF1A1 MRPL17 EIF4A1 RPL26 RPL29 MRPL55 RPL22L1 RPN1 RPS14 RPL27A SEC11C MRPL16 RPL13 EEF2 MRPL1 MRPL13 MRPL52 TRMT112 RPL4 MRPL11 EIF3F AURKAIP1 RPS27 FARSA GADD45GIP1 SSR4 MRPL14 MRPL41 EIF3C MRPL30 RPL14 RPS26 RPL10A MRPL38 MRPL12 |
| | 4.48E-23 | 57 | 214 | 5.15 | R-HSA-6791226 Major pathway of rRNA processing in the nucleolus and cytosol | MTREX RIOK2 RPL31 BUD23 EXOSC5 RPS5 NOP14 XRN2 RPL6 RPLP0 SNU13 RPL3 RPS16 RPS19 RPL18A DDX49 RBM28 RPL28 RPL34 BYSL RPL24 RPS15 UTP25 UTP20 RPL21 RPL5 RRP36 RPL23 RPL36 RPL27 LTV1 TEX10 NOB1 RPL13A RPL11 RPS8 RPL7 RPP30 RPL30 UTP14A RPL26 RPL29 RPL22L1 GNL3 RPS14 DCAF13 BMS1 RPL27A RPL13 TSR1 RPL4 RPS27 RPL14 RPS26 RPL10A DDX47 NOL12 |

| | | | | | | |
|---|---|---|---|---|---|---|
| | 6.66E-23 | 63 | 263 | 4.63 | R-HSA-72312 rRNA processing | ELAC2 MTREX RIOK2 THUMPD1 RPL31 BUD23 EXOSC5 RPS5 DIMT1 NOP14 XRN2 RPL6 RPLP0 SNU13 RPL3 RPS16 RPS19 RPL18A DDX49 RBM28 RPL28 RPL34 BYSL RPL24 RPS15 UTP25 UTP20 RPL21 RPL5 RRP36 RPL23 RPL36 DKC1 RPL27 LTV1 TEX10 NOB1 RPL13A RPL11 RPS8 NHP2 RPL7 RPP30 RPL30 UTP14A RPL26 RPL29 RPL22L1 GNL3 RPS14 DCAF13 BMS1 RPL27A RPL13 TSR1 TRMT112 RPL4 RPS27 RPL14 RPS26 RPL10A DDX47 NOL12 |
| | 2.07E-23 | 66 | 281 | 4.54 | R-HSA-72203 Processing of Capped Intron-Containing Pre-mRNA | NUP160 MTREX POLR2B TPR NDC1 YBX1 GPKOW NUP37 CHERP SRRT SF3B2 PAPOLA NUP50 AAAS NUP188 SF3A1 SNU13 SRSF5 PABPN1 PRPF6 CRNKL1 NUP93 POLR2C POLR2I HNRNPUL1 SUGP1 NUP88 DDX5 EFTUD2 SRSF7 PRPF3 CPSF3 NUP153 UPF3B GTF2F1 SNRPD2 SYMPK SNRPB NUP214 LSM4 RBM17 PRPF4 NCBP1 NUP54 EIF4A3 TXNL4A SNRNP200 DDX46 EIF4E RANBP2 NUP35 U2SURP SLBP LSM6 CPSF2 SF1 CD2BP2 PCBP1 HNRNPF DDX23 SART1 SF3A3 GTF2F2 PRPF40A PCBP2 DDX42 |
| | 3.97E-36 | 146 | 848 | 3.33 | R-HSA-8953854 Metabolism of RNA | UPF1 ELAC2 NUP160 MTREX POLR2B TPR RIOK2 NDC1 EIF4B DDX20 YBX1 THUMPD1 GPKOW PABPC1 RPL31 BUD23 CLNS1A NUP37 EXOSC5 TNPO1 RPS5 CHERP DIMT1 SRRT NOP14 SF3B2 XRN2 RPL6 RPLP0 PAPOLA PUS7 NUP50 AAAS NUP188 SF3A1 SNU13 RPL3 SRSF5 PABPN1 PRPF6 CRNKL1 NUP93 POLR2C RPS16 POLR2I HNRNPUL1 RPS19 RPL18A DDX49 SUGP1 RBM28 RPL28 NUP88 DDX5 EFTUD2 RPL34 CNOT2 BYSL RPL24 PSMD14 RPS15 SRSF7 PRPF3 UTP25 CPSF3 SET UTP20 RPL21 RPL5 RRP36 NUP153 UPF3B GTF2F1 RPL23 SNRPD2 SYMPK SNRPB NUP214 RPL36 LSM4 DKC1 RPL27 RBM17 LTV1 PRPF4 TEX10 NCBP1 NUP54 NOB1 EIF4A3 TXNL4A RPL13A RPL11 RPS8 SNRNP200 DDX46 NHP2 CDKAL1 RPL7 RPP30 EIF4E RANBP2 RPL30 UTP14A SMG1 ADAR EIF4A1 RPL26 RPL29 NUP35 RPL22L1 U2SURP GNL3 SLBP LSM6 RPS14 DCAF13 BMS1 CPSF2 RPL27A RPL13 TSR1 SF1 CD2BP2 PCBP1 HNRNPF TRMT112 CTU2 DDX23 RPL4 SART1 PUS1 RPS27 |

| | | | | | | |
|---|---|---|---|---|---|---|
| | | | | | | APOBEC3B SF3A3 GTF2F2 RPL14 SUPT5H PRPF40A PCBP2 RPS26 DDX42 RPL10A DDX47 NOL12 CD44 |
| | 4.73E-21 | 122 | 896 | 2.63 | R-HSA-2262752 Cellular responses to stress | NUP160 CUL3 CUL7 TPR NDC1 PDIA5 ME1 RPL31 UBE2D1 NUP37 EXOSC5 RPS5 RPL6 RPLP0 BLVRB DNAJB11 NUP50 AAAS NUP188 HSP90AB1 KDELR3 RPL3 HM13 RBBP7 DNAJC3 NUP93 GSR RPS16 RPS19 RPL18A EZH2 BLVRA CA9 RPL28 NUP88 NFKB1 RPL34 WFS1 TCIRG1 SOD2 MRPL18 LMNB1 RPL24 ATP6V1A PSMD14 RPS15 PRDX6 RPL21 RPL5 CDK2 NUP153 RPL23 ID1 PRDX5 NUP214 RPL36 HELZ2 ATP6V1E1 COX4I1 RPL27 RPA1 H3-3B TERF2 LAMTOR5 ETS1 CDK4 ATP6V1G1 TUBB2A HMGA1 SEC31A NUP54 NCOR1 RPL13A RPL11 RPS8 ATP6V1B2 RPL7 SERPINH1 ATM BAG3 HSPB8 RANBP2 ATP6V1C1 MOV10 RPL30 EEF1A1 CCAR2 RPL26 RPL29 NUP35 IGFBP7 RPL22L1 HSPA4L RPS14 HSPA12A RPL27A RPL13 DNAJC7 H1-4 STAT3 CYCS RPL4 UBE2C TUBB6 TBL1XR1 RPS27 CALR P4HB RXRA RPS19BP1 RPL14 H1-0 NCOR2 RPS26 H2BC12 ERO1A TFDP1 GFPT1 TXNRD1 RPL10A TUBB3 H4C1 |
| | 2.05E-29 | 244 | 2197 | 2.15 | R-HSA-392499 Metabolism of proteins | PAF1 PGM3 NUP160 ALG1 CUL3 RAB27B CUL7 TPR MRPL43 SEC61A1 NDC1 EIF4B TRAM1 RAB27A PABPC1 RPL31 LMCD1 UBE2D1 SMC1A LMAN1 NUP37 NFKB2 UBE2T UBE2K RPS5 EIF3I MMP2 MAVS XRN2 RPL6 RPLP0 CLSPN NUP50 AAAS DHPS ARCN1 NUP188 NANS MRPS18A KDELR3 DDX17 RPL3 TAB1 EIF3D RANGAP1 PMM1 SRP54 SEC23A STAG2 RBBP7 USP11 TIMP1 DNAJC3 NUP93 EARS2 EIF3E NUCB1 PPP6R1 RPS16 RPS19 RPL18A EIF4H SMC3 RPL28 FBXL20 NUP88 DDX5 KAT2A RPL34 WFS1 CTSC MRPL51 CMAS MRPL18 MRPL2 SEC24A RARS1 RPL24 MRPL3 PSMD14 RPS15 SPTBN1 MRPL19 CCT4 MRPL37 EDEM3 CDC20 CTSD SPCS2 FBXO30 TCP1 RPL21 MRPS2 COPA RPL5 RAB9A NAGK AARS2 NUP153 MRPS7 RPL23 UBA2 KTN1 NUP214 TRAF2 FBXL12 COPB1 DAD1 RPL36 EIF3G DNMT1 ULBP2 GFPT2 RPL27 TOP2A MRPL35 RPA1 H3-3B ERAL1 PCNA DAP3 DCAF8 DPH2 CCDC59 DPH6 CDC73 NARS1 CKAP4 LMO7 UGGT1 MRPL50 PDCL |

| | | | | | | |
|---|---|---|---|---|---|---|
| | | | | | | TUBB2A RAB30 MRPL15 SEC31A NUP54 USO1 CUL4A USP3 PML MFGE8 PTRH2 SAE1 RPL13A RPL11 RPS8 MRPL9 GALNT2 PARP1 MRPS5 RPL7 UHRF2 ARFGAP2 INCENP MRPL49 EIF4E RANBP2 MRPL39 RNF20 GNAQ RPL30 EEF1A1 CD109 NSMCE2 MRPL17 CUL4B COPG2 ADAMTS4 GNE EIF4A1 RPL26 UBXN1 RPL29 MXRA8 MRPL55 NUP35 IGFBP7 RPL22L1 RPN1 RPS14 RAD21 DCAF13 RAB8B FBN1 RPL27A SEC11C MRPL16 RPL13 EEF2 STAT3 MRPL1 SDC2 CDK1 LAMB2 MRPL13 GNB2 GNG12 MRPL52 TRMT112 RPL4 MRPL11 UBE2C EIF3F RAB6A AURKAIP1 TUBB6 SEC24C RPS27 AURKB FARSA CALR RAD23A GADD45GIP1 SSR4 MRPL14 COPG1 MRPL41 MTA1 NUDT14 EIF3C COPB2 MRPL30 P4HB RXRA COMMD6 RPL14 WDR5 NCOR2 SPTAN1 RPS26 H2BC12 ERO1A CSF2RA GFPT1 CTR9 RPL10A MRPL38 GNG10 TUBB3 MRPL12 SEC22B H2AC15 H4C1 |
| | 3.03E-21 | 231 | 2327 | 1.92 | R-HSA-1430728 Metabolism | CYP51A1 NDUFAF7 GGCT STARD3NL MDH1 OSBPL5 CD44 NUP160 PNPLA6 RETSAT TPR AKR7A2 NDC1 SLC2A3 ACAA1 GPC1 OAT ME1 IDI1 PYGM RPL31 HMMR SDHA HACD3 NUP37 PLD1 NT5C2 FDFT1 AACS ME2 RPS5 HADHA ACOX3 MTMR2 L2HGDH RPL6 RPLP0 BLVRB SLC9A1 PDPR NUP50 AAAS NUP188 CRAT HSP90AB1 TECR GGT1 SLC25A1 RPL3 ABHD4 SEC23A PYGB SMS GLA ACP5 GDPD3 NUP93 NME3 QPRT CDIPT GSR RPS16 PLD3 RPS19 RPL18A OGDH CAV1 BLVRA PTGR1 CA9 ACBD5 RPL28 NUP88 RPL34 UGDH PDHX CHKA LPCAT3 NNT CERT1 SEC24A RARS1 RPL24 ABHD14B PSMD14 RPS15 GLS KYNU DHCR24 MTR SDHB HMGCL SLC2A1 MTARC2 ALDH6A1 MTHFD1L RPL21 RPL5 SLC25A16 NQO2 NUP153 SLC25A19 RPL23 UROD ASL NUP214 ECHS1 POR GNAI1 TST DUT RPL36 HELZ2 ASS1 ACSS2 COX4I1 RPL27 ACLY HSD17B7 INPP5K MORC2 IMPA1 HSD17B4 GSTM3 FPGS SQOR HADHB NUP54 MMAB GALNS NCOR1 RPL13A NOSIP RPL11 RPS8 MGST3 SCD5 PPIP5K2 MMUT NSDHL RPL7 ALAD PTGES GLUD1 AASDHPPT B3GAT3 ACAD8 QDPR RANBP2 RPIA DDAH1 |

| | | | | | | |
|---|---|---|---|---|---|---|
| | | | | | | HS2ST1 DBI SLC26A2 GNAQ RPL30 CPT2 CBR3 MED27 FDPS RPL26 RPL29 BPNT1 NUP35 GNPDA2 RPL22L1 CMBL RPS14 ALDH7A1 PDP1 PGM2L1 INPPL1 NDUFB8 SMPD1 IDH3A RPL27A NNMT ACSF2 RPL13 ATP5PD TK1 MAT2A SDC2 DCXR UGP2 RBKS NAT1 ETFDH CYCS SPTLC3 BPGM GNB2 GNG12 CES2 DHCR7 TRMT112 PARP14 GLRX PC NMNAT1 RPL4 TYMS SEC24C TBL1XR1 RPS27 IMPDH2 SLC25A20 IDH2 RXRA AKR1C1 RPL14 ACADSB NCOR2 SULT1A1 ADA FUT11 ACSL5 GSTK1 RPS26 UCKL1 TXNRD1 PAPSS2 ECI2 RPL10A GK SACM1L GSTM2 PPP1CB DHFR GNG10 HMBS |
| Wiki-pathway | 2.11E-04 | 7 | 15 | 9.02 | WP4240 Reg. of sister chromatid separation at the metaphase-anaphase transition | MAD2L1 SMC1A ESPL1 SMC3 BUB3 CDC20 RAD21 |
| | 1.47E-17 | 33 | 88 | 7.25 | WP477 Cytoplasmic ribosomal proteins | RPL24 RPL27 RPS27 RPL26 RPS26 RPL27A RPL23 RPL30 RPL29 RPL28 RPL13A RPS5 RPL34 RPL3 RPL4 RPS8 RPL5 RPL36 RPL31 RPL10A RPLP0 RPL14 RPL11 RPL13 RPL6 RPL7 RPS15 RPS14 RPL18A RPL21 RPS19 MRPL19 RPS16 |
| | 9.30E-06 | 13 | 42 | 5.98 | WP466 DNA replication | MCM3 MCM5 MCM6 MCM7 RFC2 MCM10 CDK2 PRIM1 PCNA ORC6 RFC4 RPA1 POLD2 |
| | 2.22E-06 | 15 | 50 | 5.80 | WP107 Translation factors | EIF4B EIF4A1 EIF2AK2 EEF2 EIF4E EIF3C EIF3G EIF3D EIF3F EIF3E EIF4H EIF3I EEF1A1 PABPC1 CLUH |
| | 1.17E-03 | 9 | 33 | 5.27 | WP4786 Type I collagen synthesis in the context of osteogenesis imperfecta | P3H1 CRTAP P4HA2 SERPINH1 P4HA1 PLOD2 P4HB P3H2 LOX |
| | 2.22E-13 | 34 | 126 | 5.21 | WP411 mRNA processing | RBM39 EFTUD2 CELF1 NCBP1 NONO SNRPB XRN2 PRPF6 SF3A3 SNRPD2 CD2BP2 PRPF4 PRPF3 TXNL4A YBX1 PAPOLA SF3B2 DDX20 PRPF18 SF3A1 SRSF5 SRSF7 SFSWAP SUPT5H SNU13 CPSF2 PABPN1 CPSF3 SUGP1 RBM17 SRP54 PRPF40A SMC1A PCBP2 |

| | | | | | | |
|---|---|---|---|---|---|---|
| | 2.59E-09 | 24 | 90 | 5.15 | WP2446 Retinoblastoma gene in cancer | MCM3 MCM6 MCM7 KIF4A CDK4 CCND1 CDK2 RBBP7 TYMS PCNA DNMT1 ANLN RPA1 RFC4 MSH6 SMC3 TFDP1 CCNB1 CHEK1 DHFR PRIM1 TOP2A SMC1A CDK1 |
| | 4.41E-05 | 15 | 64 | 4.53 | WP45 G1 to S cell cycle control | MCM3 MCM5 MCM6 MCM7 ATM CDK4 CCND1 CDK2 PCNA RPA1 TFDP1 CCNB1 PRIM1 CDK1 ORC6 |
| | 4.41E-05 | 17 | 81 | 4.06 | WP4016 DNA IR-damage and cellular response via ATR | HUS1 ATM CDK2 UPF1 PCNA TDP1 RPA1 CLSPN CHEK1 SMARCC2 PML USP1 SMC1A PARP1 CDK1 MSH2 RECQL |
| | 4.70E-05 | 18 | 91 | 3.82 | WP3925 Amino acid metabolism | ACLY P4HA2 PDHX GLUD1 GLS OGDH ASS1 MMUT MDH1 RARS1 OAT ACAA1 SDHA HMGCL PC ALDH7A1 SMS GSR |
| | 4.48E-06 | 23 | 120 | 3.70 | WP179 Cell cycle | MCM3 MCM5 MCM6 MCM7 BUB3 ATM CDC20 CCND1 RBL1 SMC3 CCNB1 E2F4 CHEK1 ESPL1 CDK4 CDK2 STAG2 PCNA TFDP1 SMC1A CDK1 ORC6 RAD21 |
| | 5.04E-04 | 19 | 119 | 3.09 | WP4946 DNA repair pathways full network | ATM RPA1 POLD2 MSH6 CHEK1 PARP1 CUL4A RFC2 CUL4B ERCC5 UNG PCNA TERF2 RFC4 USP1 POLM RAD23A MSH2 MSH3 |
| | 1.62E-11 | 63 | 431 | 2.82 | WP3888 VEGFA-VEGFR2 signaling pathway | CLTC LMAN1 FLNB PXN CSRP2 RPL10A CALR IDH2 MYO1C CAPN2 MAP2K1 SLC25A25 ETS1 ITGAV NFKB1 EWSR1 SRP54 GIGYF2 CRIP2 CAV1 PTPN1 ENG COPG1 MMP2 SSR4 VPS39 MOV10 IGFBP7 ATP6V1E1 STAT1 STAT3 ICAM1 SOD2 EPS15 STAT6 SET P4HA2 RPL13A CYCS EZR CCND1 INPP5K P4HB EIF3D EIF3F RPL5 RPL7 RPL18A RPL27 RPL26 QKI NDRG1 GRSF1 RACK1 PABPC1 GPC1 RBM39 PRDX6 EIF4E EIF4G2 GLUD1 CSK FAM120A |
| | 1.10E-04 | 29 | 214 | 2.62 | WP4352 Ciliary landscape | MCM3 MCM5 MCM6 MCM7 YPEL5 CD2BP2 TBC1D4 PAFAH1B1 ANKS3 CTBP2 SMC4 ARHGDIA NFKB1 MCM10 MYL6B MSH2 SNAP29 EFTUD2 RALB AFG3L2 WDR26 GLA MFAP1 DYNC2I2 NUP88 DDX5 ECHS1 LSM4 SSNA1 |
| | 7.15E-04 | 35 | 314 | 2.15 | WP2882 Nuclear receptors meta-pathway | NRG1 CCND1 GGT1 RXRA CAVIN2 ME1 KTN1 ACAA1 SULT1A1 CDK4 FGD4 CPT2 HSP90AB1 DNAJC7 NFKB2 SLC26A2 SMC1A PTGR1 CDK1 GSR PRDX6 CBR3 GSTM3 GSTM2 TXNRD1 DBI BLVRB |

| | | | | | | |
|---|---|---|---|---|---|---|
| | | | | | | SLC2A1 SLC2A3 STAT3 CES2 MFGE8 CAP2 AKAP13 MGST3 |
